# Change-of-mind neuroeconomic decision-making is modulated by LINC00473 in medial prefrontal cortex in a sex-dependent manner

**DOI:** 10.1101/2024.05.08.592609

**Authors:** Romain Durand-de Cuttoli, Orna Issler, Samantha M. B. Pedersen, Benjamin Yakubov, Nusrat Jahan, Aisha Abid, Susanna Kasparov, Kastalia Granizo, Sanjana Ahmed, Scott J. Russo, Eric J. Nestler, Brian M. Sweis

**Affiliations:** Nash Family Department of Neuroscience, Friedman Brain Institute, Icahn School of Medicine at Mount Sinai, New York, NY 10029; Neuroscience Institute, Department of Anesthesiology, Perioperative Care and Pain Medicine, and Department of Neuroscience and Physiology, NYU Langone, New York, NY 10016; CUNY Hunter, New York, NY 10023; Department of Psychiatry, Friedman Brain Institute, Icahn School of Medicine at Mount Sinai, New York, NY 10029

## Abstract

Changing one’s mind involves re-appraisals between past-costs versus future-value and may be altered in psychopathology. Long intergenic non-coding RNA LINC00473 in medial prefrontal cortex (mPFC) can induce stress-resilience in a sex-dependent manner, but its role in cognition is unknown. We characterized decision-making behavior in male and female mice in the neuroeconomic paradigm Restaurant Row following virus-mediated expression of LINC00473 in mPFC. Mice foraged for food among varying temporal-costs and subjective-value while on a limited time-budget. Without affecting primary deliberative decisions, LINC00473 selectively influenced re-evaluative choices in a sex-dependent manner. This included changing how mice (i) cached value with the passage of time and (ii) weighed prior mistakes, which underlie the computational bases of sensitivity to sunk costs and regret. These findings suggest a common value function is shared between these neuroeconomic processes and reveal a bridge between molecular drivers of stress-resilience and psychological mechanisms underlying sex-specific proclivities in negative rumination.

## Introduction

Understanding the mechanisms by which an individual changes one’s mind when making choices is a complex psychological phenomenon that has recently gained heightened attention in the field of decision science ^1^. Change-of-mind decisions involve re-evaluative, secondary processing of recently selected choices. These types of choices may be motivated by several factors, including attempts to correct recent decisions. Investigating these behaviors presents a unique opportunity to peer into how the brain represents internal conflict at multiple levels ^2,3^. Conflict can be engaged at both the economic and psychological levels. The economic level may seek to manage ongoing expenditures while both minimizing potential losses and maximizing future gains ^4^. The psychological level may be more focused on self-criticism of one’s own recent actions, which may be useful for learning but could be perceived as unpleasant or egodystonic ^5–7^. Aspects of these processes could contribute to one’s affective burden and may manifest as changes in both mood and cognition observed in psychiatric disorders ^8,9^.

Depression is a debilitating mental illness characterized by pervasive mood disturbances, altered cognitive capabilities, and maladaptations in motivated behavior ^9^. The incidence of depression is two times higher in women than in men and can manifest different symptomatology, including increased negative rumination in females ^10^. However, the biological mechanisms mediating these sex differences remain unclear ^11–15^. Progress has been made toward developing animal models of aspects of negative rumination. This has included breakthroughs in capturing across species the behavioral and neurophysiological signatures of regret processing [counterfactual representations of missed opportunities] and sunk costs [sensitivity to irrecoverable losses], both of which hinge on change-of-mind behavior and can drive negative rumination ^5,16–18^. However, it remains unclear how psychiatric vulnerabilities in negative rumination depend on the neuroeconomic principles underlying change-of-mind decision-making. Investigating the neurobiology of sex differences in depression using this framework can not only better link changes in mood to behavior, but also provide a refined perspective on decision-making computations. This includes examining how, for instance, regret and sunk cost sensitivity relate to one another.

Progress is being made toward appreciating the neuroanatomical, genetic, epigenetic, and pathophysiological differences between depressed males and females ^19,20^. Clinical and preclinical studies using sex as a biological factor have adopted innovative approaches leveraging unsupervised analyses of large datasets leading to recent breakthroughs in systems biology. This has included identifying key regulatory roles of non-coding elements of the genome ^21^. Recent transcriptomics work carried out in human postmortem brain tissue samples obtained from subjects carrying a diagnosis of major depressive disorder discovered sex-dependent functions of a long intergenic non-coding RNA: LINC00473 ^22^. Overall, long non-coding RNAs represent a substantial percentage of all of the regulated transcripts in human depression and are regulated in a sex- and brain region-specific manner ^22^. Lower LINC00473 expression levels were found in the medial prefrontal cortex (mPFC) of depressed females only, while there were no differences observed in expression levels between healthy males and females ^22^. Some studies have implicated LINC00473 in molecular scaffolding, inter-organelle communication, and transcriptional regulation in non-neuronal tissue, while exhibiting immediate early gene-like properties in cultured neurons. However, LINC00473 function remains largely unknown with no identified role in cognition ^23–28^.

Studies in mice demonstrated that virus-mediated expression of LINC00473 in mPFC neurons promoted resilience to stress in females only ^22^. This included ameliorating depressive- and anxiety-like behavioral abnormalities, while blunting hypothalamic-pituitary-adrenal-axis activity as well as the induction of gene expression changes induced in this brain region by chronic stress ^22^. Beyond this, LINC00473 has not been studied in neural tissue in any other capacity, with no work performed to date on the role of long non-coding RNAs in regulating decision-making. LINC00473 has been shown to be regulated by conserved Ca^2+^/calmodulin signaling pathways and is likely induced by cAMP response element-binding protein (CREB) ^22–24,26^. CREB has long been studied through traditional candidate gene approaches and shown to regulate numerous other genes involved in neuronal excitability and synaptic plasticity as well as stress sensitivity ^29–34^. Like LINC00473, CREB function in the mPFC has been implicated in promoting resilience to stress, however, unlike LINC00473, this has been shown to be true for both males and females ^22,35^. These reports suggest either that there must be some divergence of interacting CREB-LINC00473-molecular pathways between sexes or that prior studies were limited in their ability to resolve more nuanced sex differences at the behavioral level. Recently, we discovered that individual differences in susceptibility vs. resilience to stress were associated with alterations in higher-order neuroeconomic decision-making processes, including sensitivity to regret and sunk costs. This work led to the discovery of two distinct types of regret-related behaviors, one of which depends on change-of-mind decision-making^16^. Heightened sensitivity to sunk costs and change-of-mind-related regret were traits previously found to be associated with stress resilience. Furthermore, CREB function in the mPFC vs. nucleus accumbens could bidirectionally regulate aspects of these processes ^16^. Here, we build upon these findings in order to characterize baseline sex differences in neuroeconomic choice behavior and the impact of LINC00473 function in mPFC in order to identify sex-specific vulnerabilities in decision-making.

Strides made in the field of neuroeconomics, an area focused on understanding how the brain is comprised of multiple valuation algorithms, have leveraged complex approaches in decision neuroscience to quantify how circuits constrain behavior ^36–39^. This encompasses characterizing multifactorial aspects of motivation that operationalize reward value along several dimensions, integrate choice processes with environmental circumstances, and segregate stages of a decision stream into its component parts ^40–43^. This work includes isolating properties of change-of-mind re-evaluative decisions, and points to the mPFC as a critical hub for regulating reward value, self-control, and mood ^4,44–48^. Insights from neuroeconomic principles offer refined approaches to investigate decision-making capable of resolving behavior into discretely measurable computational units in a manner that is biologically tractable and readily translatable across species ^40,49,50^. This approach has yielded substantial efforts developing innovative tasks adapted for use in mice, rats, monkeys, and humans ^5,51–54^. The use of these translational tasks have led to synergistic cross-species discoveries uncovering the behavioral underpinnings of signatures of higher-order cognitive processes previously thought to be unique to humans and supported by physically separable circuits in the brain ^16–18,55,56^. This offers an avenue for improved models of dysfunctional decision-making behavior.

Here, we examined how expression of LINC00473 in the mPFC alters multiple aspects of value-based decision-making behavior in male and female mice. We leveraged our longitudinal neuroeconomic foraging paradigm, Restaurant Row, to characterize complex decision-making behavior in mice transfected with either AAV2-hSyn-LINC00473 or AAV2-hSyn-GFP targeting mPFC neurons. This task has been used previously to access behavioral and neurophysiological mechanisms of multiple decision-making systems in the brain. This includes segregating within the same trial primary decisions vs. secondary re-evaluative change-of-mind decisions ^5,17,57–62^. On this task, rodents must budget 30 minutes to forage for their primary source of food by earning pellets at spatially fixed feeding sites (“restaurants”) that differ in subjective value (i.e., flavors: chocolate, banana, grape, or plain). During each trial, mice are presented with auditory cues whose pitch indicates the temporal delay required to earn a reward (e.g., low pitch corresponds to a short-delay, low-cost offer). Each restaurant is physically divided into a separate offer zone and wait zone. Rodents must make a primary decision to accept or reject an offer in the offer zone. If the offer is accepted by entering the wait zone, the tone cues investment progress by descending in pitch counting down time to earn a reward, during which mice can quit at any point (**Fig. 1a**, **Supplementary Fig. 1**). Behaviors in the offer zone and wait zone comprise separate valuation algorithms that have been previously linked to distinct circuit functions ^5,16,18,43,57–59,61,63,64^. Mice were tested daily over two months across a changing economic landscape during which complex foraging strategies developed. We report sex- and LINC00473-dependent changes in neuroeconomic decision-making behavior specifically tied to change-of-mind decisions in the wait zone, notably with alterations in sensitivity to regret and sunk costs. These data provide deeper insights into the computational processes that could be driving specific aspects of re-evaluative choice. Further, these data highlight a sex-dependent role of LINC00473 function in the mPFC in only certain types of decision-making behaviors.

**Fig. 1.**
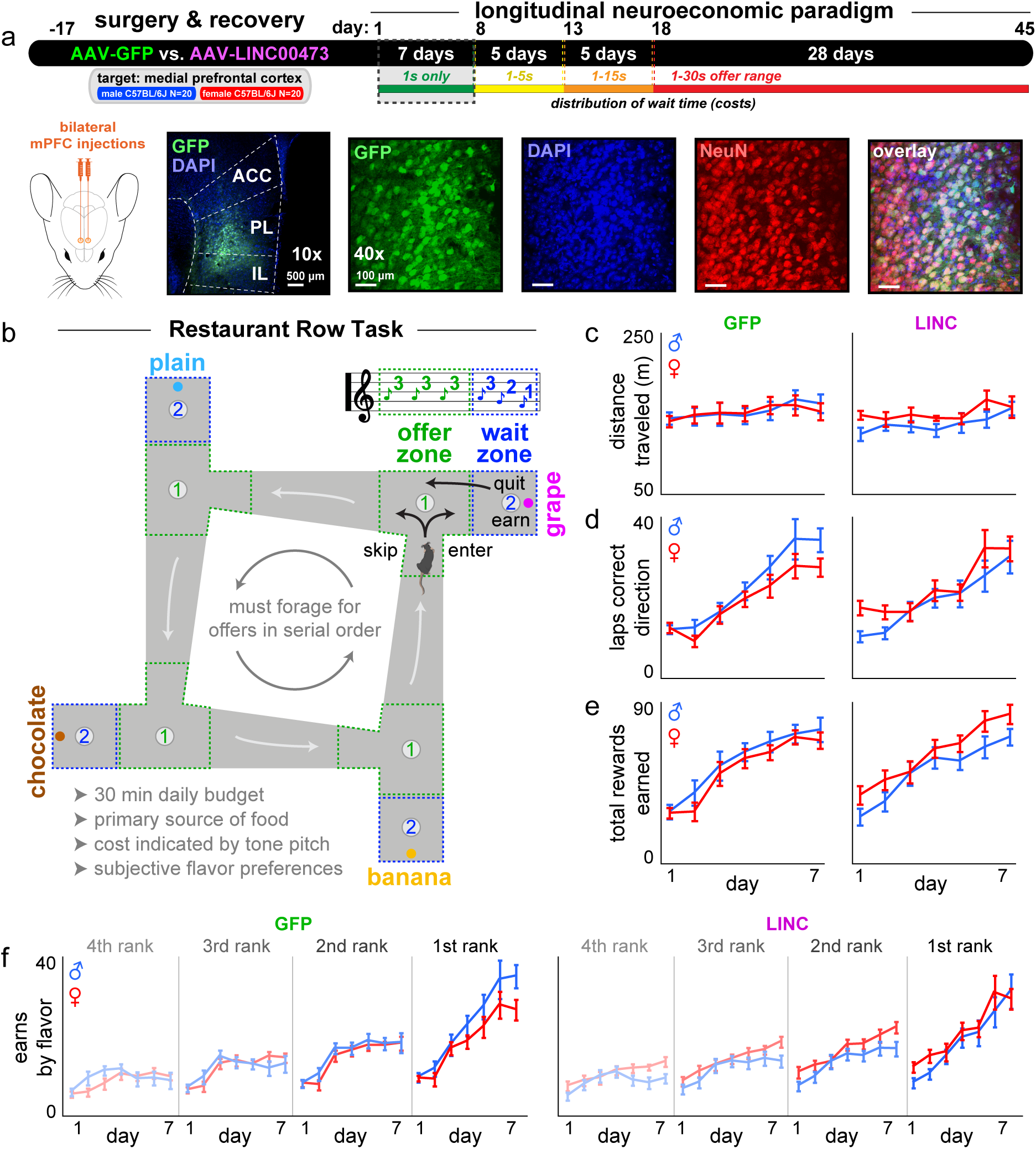
Male and female mice similarly acquire the Restaurant Row task. **a** Timeline. Representative images of surgical targeting and virus transfection of GFP at 40x and 10x magnifications with DAPI and NeuN staining. Mice were allowed to recover before beginning longitudinal testing on the Restaurant Row task for 45 consecutive days. Dashed box indicates time period relevant for this figure: week 1 of testing. **b** Task schematic. Mice were allotted 30 min daily to invest time foraging for their primary source of food in a self-paced manner. Costs to obtain rewards were in the form of delays mice would have to wait near feeder sites. Mice were required to run in a counterclockwise direction encountering offers for different flavors at each “restaurant” in serial order. Each restaurant, separated by hallways, was divided into a T-shaped “offer zone” choice point and a separate “wait zone” that housed the pellet dispenser. Upon offer zone entry from the correct heading-direction, a tone sounded whose pitch indicated the delay mice would have to wait if accepting the offer by entering the wait zone. If entered, tone pitch descended in the wait zone, cuing the indicated delay. Each trial terminated if mice skipped in the offer zone, quit during the countdown in the wait zone, or earned a reward, after which animals were required to proceed to the next restaurant. **c-f** Behavioral metrics across the first week of testing during which all offers were 1 s only (lowest pitch, 4 kHz): (c) distance traveled, (d) laps run in the correct direction, (e) rewards earned, (f) earnings split by flavors ranked from least to most preferred by summing each day’s end-of-session totals in each restaurant. No effect of sex or LINC00473 expression was grossly apparent in these initial metrics. Error bars represent ±1 SEM.

## Results

### Male and female mice acquire the Restaurant Row neuroeconomic task

We performed virus transfection surgeries on all mice and allotted 2-3 weeks for recovery before beginning the longitudinal Restaurant Row paradigm (**Fig. 1a**). During the first week of Restaurant Row testing, all reward offers were fixed at 1 s only (cued by a 4 kHz tone) as mice acquired the basic structure of the task (**Fig. 1a-b**). Mice had to learn to run in the correct direction in order to forage for low-cost food rewards. We food-restricted all mice to the same percentage (~80-85%) of baseline free-feeding body weight (**Supplementary Fig. 2**). Both males and females, regardless of GFP or LINC00473 treatment, grossly traveled similar total distances across the first week (**Fig. 1c**, sex*virus: *F*=0.971, *p*=0.331). All groups learned to run the same number of laps in the correct direction (**Fig. 1d**, day: *F*=334.219, *p*<0.0001; sex*virus: *F*=2.780, *p*=0.104) and increase their total pellets earned (**Fig. 1e**, day: *F*=393.991, *p*<0.0001; sex*virus: *F*=3.062, *p*=0.089) across the first week. All mice displayed subjective flavor preferences, determined by summing end-of-session total earns in each restaurant ranked from least to most preferred (**Fig. 1f**, rank: *F*=255.250, *p*<0.0001). There was a modest upward shift in earns split by flavor ranking in LINC00473-treated females across the first week (**Fig. 1f**, **Supplementary Fig. 3**, sex*virus: *F*=3.334, *p*=0.076). Overall, these data indicate all groups were able to demonstrate the ability to acquire the basic structure of the task and forage for food in a self-paced manner without overt differences in gross locomotor or feeding behavior.

### LINC00473 alters sex- and value-dependent, task-driven conditioned place preference behavior

Before and after the first week of behavioral testing, we characterized exploratory behavior in the maze arena (**Fig. 2a-b**). That is, without any active task, we allowed mice to roam the maze freely on day 0 and again on day 8 of the experimental timeline (**Fig. 2b**). This allowed us to probe a conditioned place preference (CPP)-like response for reward sites that developed throughout the first week of self-paced task performance. We measured the number of entries into and time spent near the feeder site in each restaurant’s wait zone on these two probe sessions (**Fig. 2c-e**). All groups explored the arena, including each restaurant, to similar degrees at baseline (day 0, ANOVA, sex*virus [entries]: *F*=0.259, *p*=0.614; sex*virus [time]: *F*=1.155, *p*=0.290, **Fig. 2c**). Overall, compared to the day 0 pre-test baseline, we found that during the day 8 post-test all mice increased total time spent (*F*=75.818, *p*<0.0001) and – to a lesser degree – entries into the reward sites (*F*=8.501, *p*<0.01, **Fig. 2c**). We then split these metrics by restaurant ranked according to day 7’s task earnings and found that CPP scores (post-pre delta scores) scaled with flavor preferences (rank [entries]: ANOVA, *F*=16.307, *p*<0.0001; rank [time]: *F*=9.829, *p*<0.0001, **Fig. 2f-g**). We found significantly lower CPP scores with respect to time spent but not entries into in the wait zone of the most preferred flavor in LINC00473-treated females only, despite being matched to males for equivalent number of rewards earned on day 7 of the task (sex*virus*rank [time]: ANOVA, *F*=2.844, *p*<0.05, **Fig. 2f-g**). This sex-dependent effect was replicated in an independent cohort of mice separately transfected with a lower titer of the LINC00473-expressing AAV (**Supplementary Fig. 4**). These findings highlight that mPFC LINC00473 expression can have sex-specific effects on certain types of actions during reward-seeking behavior (i.e., time to exit but not enter frequency) that depend on subjective value, explored in further detail below throughout the rest of Restaurant Row testing ^58,65,66^.

**Fig. 2.**
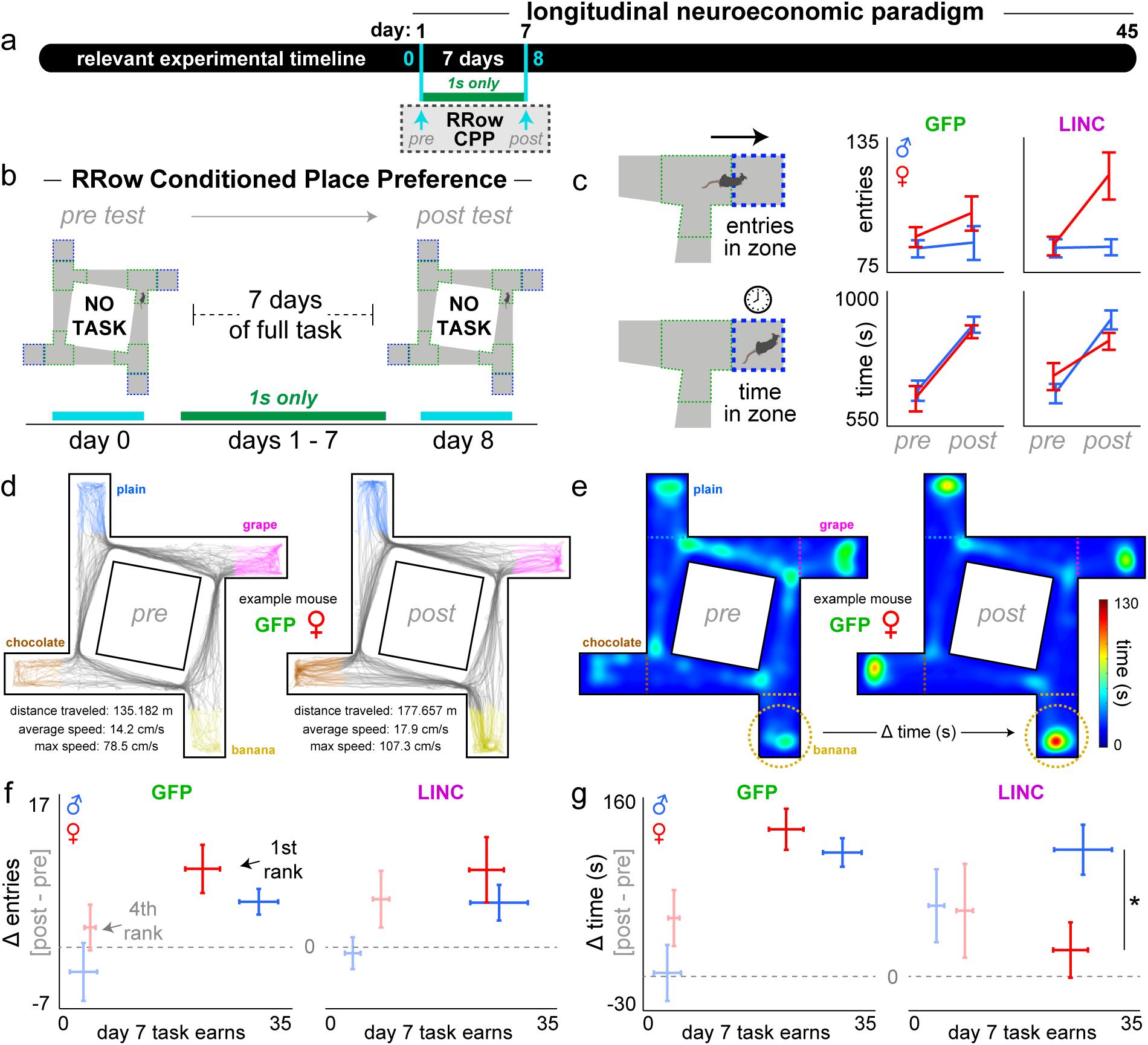
LINC00473 expression in mPFC alters sex- and value-dependent, task-driven conditioned place preference behavior. **a** Timeline. Dashed box indicates time period relevant for this figure: two special testing days marked with cyan arrows. **b** Experimental design. Animals were placed in the arena free to roam for 20 min on day 0 and day 8 of the experimental timeline but with no active task to obtain pre-task baseline and post-task experience-related exploratory behavior. **c** Total number of entries and total time spent in the wait zone, aggregated in all restaurants. **d-e** Representative video tracking data from an example mouse [GFP-treated female] recorded during the pre (day 0) and post (day 8) tests. (d) Track plots displaying body position when traveling between restaurants (gray) and when in the wait zone of each restaurant (colored). (e) Occupancy heat maps of the same example mouse in (d). The change in entries into and time spent in each restaurant’s wait zone can be calculated as a delta score (post minus pre), color of these example heat maps is scaled across sessions to the maximum occupancy (130s). **f-g** Delta scores in change of (f) total entries and (g) total time comparing post (day 8) minus pre (day 0) time points split by restaurants ranked according to rewards earned on the active task on day 7. Only the most preferred (1^st^) and least preferred (4^th^, faint) restaurants are depicted here (see **Supplementary Fig. 4** for full dataset and replication cohort). Horizontal dashed gray line indicates delta score of 0. LINC00473 expression in mPFC abolished CPP behavior in most preferred restaurants only in females and only in the time but not entry domain. **\***Represent significant sex differences. Error bars represent ±1 SEM.

### Escalating reward-scarcity unmasks sex-dependent effects of LINC00473 on foraging profiles

Next, we examined how foraging behaviors changed across the subsequent weeks as we increased the distribution of offer costs (rage of delays encountered) without adjusting the limited time budget (30 min) mice had to forage for food (**Fig. 3a**). Thus, in this “closed-economy” system, animals must adapt to the changing economic landscape of the environment as reward scarcity increased – a model of economic stress we recently characterized ^43^. We increased costs in a stepwise manner across blocks of days (i.e., offer delays ranged from 1-5 s on days 8-12, 1-15 s on days 13-17, and finally 1-30 s on days 18-45, as previously published ^5,43,63^). We randomly sampled costs from a uniform distribution.

**Fig. 3.**
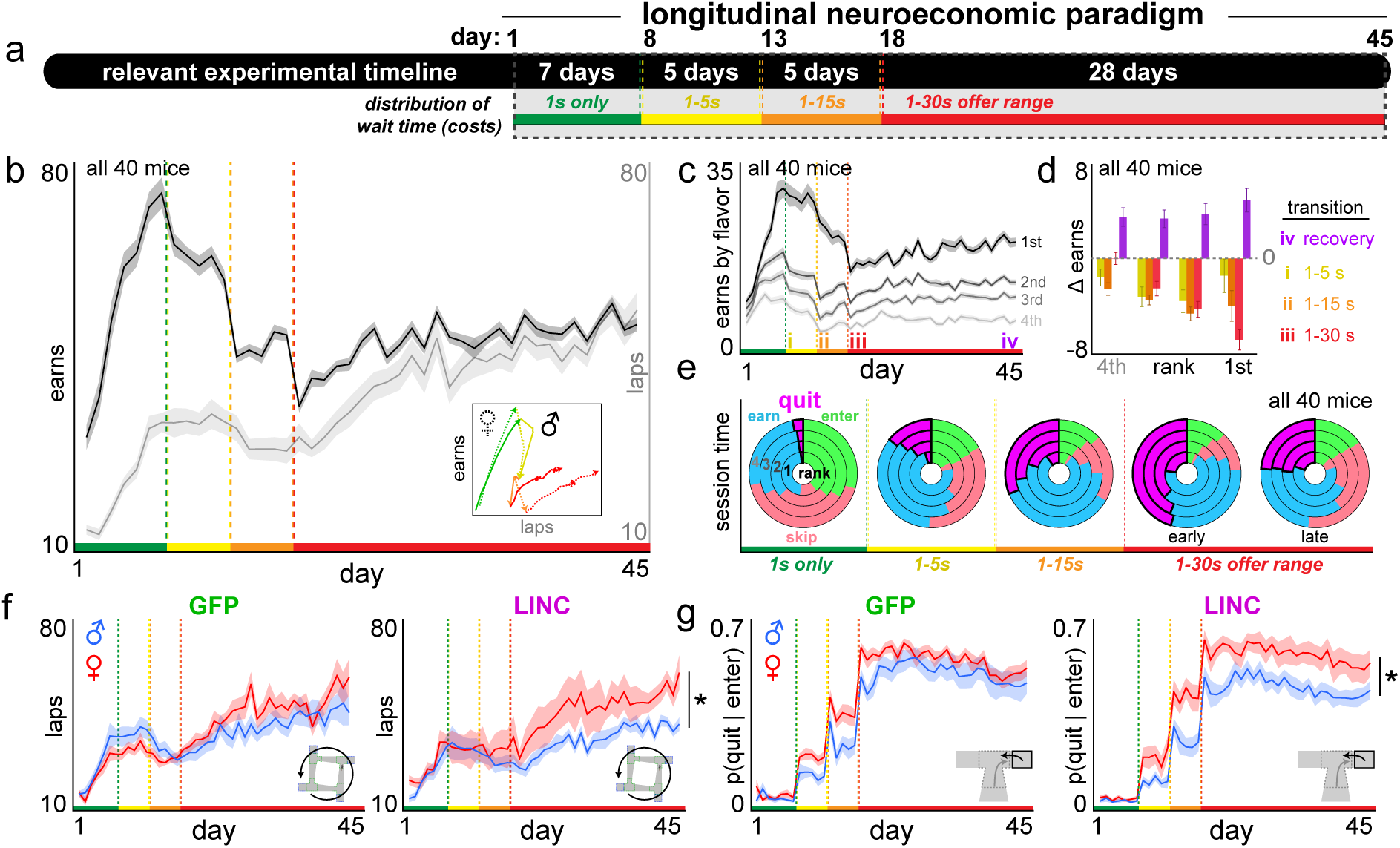
Escalating reward scarcity unmasks sex-dependent effects of LINC00473 expression in mPFC on complex foraging profiles. **a** Timeline. Dashed box indicates time period relevant for this figure: entire 45-day Restaurant Row paradigm. **b** Rewards earned (black) and laps run in the correct direction (gray). All 40 mice. Color bars along the x-axis reflect the distribution of offers illustrated in (a). Vertical dashed green-yellow-red lines indicate transition points of the stepwise schedule and are re-used throughout all other figures as a visual aid. Inset depicts “hysteresis” plot of laps against earns (epochs of days are color-coded). **c** Rewards earned split by flavors ranked from most preferred (1^st^) to least preferred (4^th^, faint). All 40 mice. **d** Change in rewards earned in each restaurant comparing two days at various color-coded transition points (delta scores calculated by subtracting the following: i, day 8-7 (yellow); ii, day 13-12 (orange); iii, day 18-17 (red); iv, day 45-18 (purple). All 40 mice (see **Supplementary Fig. 5** for group comparisons). **e** Portion of total session time engaged in four major choice behaviors: offer zone – enter (green) or skip (red); wait zone – quit (magenta) or earn (blue). Rings split restaurants ranked from most (1^st^, inner ring) to least preferred (4^th^, outer ring). Each of the 5 pie charts represent data from each testing stage, with the early and late epochs during the 1-30 s offer range from the first 5 and last 5 days of that period. **f** Laps run in the correct direction split by sex and treatment. **g** Proportion of enter decisions terminating in a quit outcome once in the wait zone. LINC00473 expression in mPFC drove females to run more laps in response to a worsening economic landscape and quit more frequently. **\***Represent significant sex differences. Shading / error bars represent ±1 SEM.

As expected, we found a decrease in pellets earned in response to each stepwise change in offer costs (sign test [−] on earns: day after minus day before transition, collapsed across epochs: *t*=−12.655, *p*<0.0001, **Fig. 3b**). These deficits in earnings scaled with flavor preferences and were matched between all groups (rank: *F*=6.938, *p*<0.001, timepoint*rank: *F*=3.651, *p*<0.01; sex*virus: *F*=0.016, *p*=0.899, **Fig. 3c-d**, **Supplementary Fig. 5**). Across the final epoch of testing (the 1-30 s offer range period: days 18-45), mice displayed a complex behavioral response profile that depended on running more laps in order to increase and recover total food earnings (laps [days 18-45]: *F*=355.340, *p*<0.0001; earns [days 18-14]: *F*=252.218, *p*<0.0001, **Fig. 3b-d**). All groups were matched in how much food was recovered across the final epoch of testing (sex*virus: *F*=0.234, *p*=0.631, **Fig. 3d**, **Supplementary Fig. 5**). However, we found that overall females displayed a greater increase in the number of laps run compared to males throughout this process (sex*day: *F*=7.848, *p*<0.01, **Fig. 3b,f**, **Supplementary Fig. 5**).

The most striking change in choice behavior across the experiment associated with the changes in laps run and rewards earned was the proportion of session time re-allocated to quitting behavior in the wait zone. This metric significantly increased after each stepwise change in offer costs (timepoint*choice: *F*=226.523, *p*<0.0001, **Fig. 3e**) and more prominently in females (timepoint*sex [quits]: *F*=11.821, *p*<0.0001, **Supplementary Fig. 6**). Interestingly, while sex differences in wait zone quitting behavior disappeared in the final epoch of testing for the GFP control group, LINC00473-treated mice instead displayed a widening of sex differences in quitting behavior (sex*virus: *F*=8.201, *p*<0.01, **Fig. 3g**, **Supplementary Fig. 6**). Furthermore, LINC00473-treated females displayed shorter latencies to quit in the wait zone compared to males, particularly for more preferred flavors (sex*virus*day*rank: *F*=2.805, *p*<0.05), despite no changes in offer zone enter frequency or offer zone reaction times (*F*=1.713, *p*=0.162, **Supplementary Fig. 6**). This effect of LINC00473 in the wait zone is reminiscent of the changes in female behavior during the CPP probe sessions (see **Fig. 2**). Taken together, these data reflect rearrangements in complex foraging patterns that emerged across longitudinal time scales in response to worsening economic demand. Changes in laps run and rewards earned were ultimately driven by more rigorous quitting behavior in LINC00473-treated females. These analyses reveal sex-dependent effects of mPFC LINC00473 expression that alter quit frequency and quit reaction times, which were not unmasked until mice encounter a reward-scarce environment.

### Change-of-mind decision-making patterns reveal unique sex-dependent neuroeconomic policies

To characterize the economic nature of the decisions made in the offer zone and wait zone, we quantified decisions to enter vs. skip (offer zone) or earn vs. quit (wait zone) as a function of both the cued offer cost and flavor identity (**Fig. 4a-c**). All groups were capable of making cost-informed enter vs. skip choices in the offer zone or earn vs. quit choices in the wait zone. Meaning, all mice were able to discriminate tones when making choices (offer*flavor [offer zone]: *F*=13.129, *p*<0.001; offer*flavor [wait zone]: *F*=18.006, *p*<0.001, **Fig. 4c**). Furthermore, how offer costs altered enter vs. skip and earn vs. quit probabilities scaled with flavor preferences in all groups (**Fig. 4b**). To summarize these behaviors, we calculated thresholds of willingness to enter in the offer zone and, separately, willingness to wait to earn a reward in the wait zone by fitting a Heaviside step regression to the trial outcome as a function of the delay that is cued at the start of each trial (note: this metric does not include reaction time or latency to make the choice). We then extracted the inflection point in each restaurant (**Fig. 4d, inset**). Thus, these measures capture the offer cost at which 50% of trials are entered in the offer zone and, separately, earned in the wait zone.

**Fig. 4.**
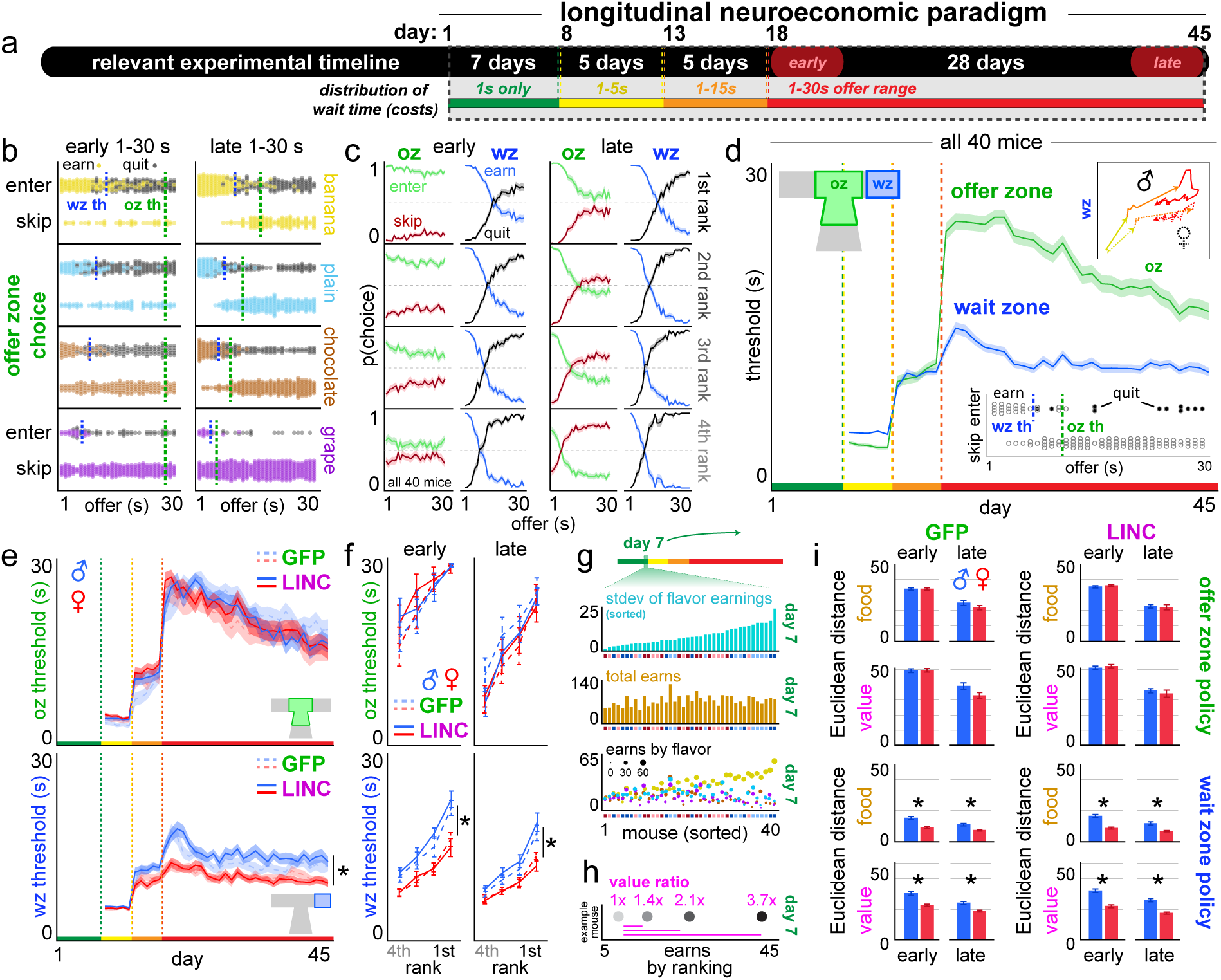
Change-of-mind decision-making patterns reveal unique sex-dependent neuroeconomic policies. **a** Timeline. **b** Example behavior from a single mouse during the 1-30 s epoch (early: days 18-22, late: days 41-45). Example offer zone and wait zone thresholds indicated by the vertical dashed green (oz th) and blue (wz th) lines respectively in each restaurant. **c** Summary data from all 40 mice early or late during the 1-30 s epoch. **d** Thresholds of willingness to accept offers in the offer zone or earn offers in the wait zone. Inset depicts an example session from a single mouse in one restaurant. All 40 mice. Top inset depicts “hysteresis” plot of offer zone against wait zone thresholds (epochs color-coded). **e** Offer zone thresholds (top) and wait zone thresholds (bottom) split by sex and treatment. **f** Thresholds in (e) from early and late 1-30 s epoch. **g-i** Economic analysis of how decision policies during the 1-30 s epoch approximate maximal food vs. subjective value (see **Methods** and **Supplementary Fig. 7** for full explanation). (g) Individual differences in (top) standard deviation of rewards earned across flavors on day 7, (middle) total rewards earned on day 7, and (bottom) earns split by restaurant (flavor: chocolate-brown, banana-yellow, grape-purple, plain-cyan) and size (reward count) on day 7. All data in (g) is sorted by standard deviation data in (top). Small colored boxes along the x-axis indicate sex (male-blue, female-red) and treatment (LINC-dark, GFP-light). **h** Example data from a single mouse on day 7 showing earn ratios between flavors. **i** Data from (g-h) used in a computational modeling analysis to calculate the Euclidean distance between observed and theoretical decision policies for maximal food (brown) or subjective value (magenta) separately for offer zone and wait zone thresholds. **\***Represent significant sex differences. Shading / error bars represent ±1 SEM.

Overall, in all groups, we found that offer zone thresholds significantly differed from wait zone thresholds across epochs of testing (timepoint*zone: *F*=296.648, *p*<0.0001, **Fig. 4d**). Upon transitioning to the 1-30 s epoch (day 18), offer zone thresholds drastically increased in all groups to similar degrees while retaining the ordinal ranking of flavor preferences (rank: *F*=17.247, *p*<0.0001; sex: *F*=0.170, *p*=0.681; sex*virus: *F*=0.031, *p*=0.861, **Fig. 4d-f**). Wait zone thresholds, on the other hand, increased only in male mice following the transition to the 1-30 s epoch (rank: *F*=58.574, *p*<0.0001; sex: *F*=17.776, *p*<0.0001; sex*virus: *F*=1.238, *p*=0.268, **Fig. 4d-f**). Over the subsequent week (approximately days 18-24), wait zone thresholds in males decreased before stabilizing across the remainder of the experiment (approximately days 25-45, **Fig. 4d-f**). In females, wait zone thresholds (i) remained relatively unchanged upon transition to day 18, (ii) were significantly lower than males, and (iii) remained stable across the entire 1-30 s epoch (sex*rank [offer zone]: *F*=0.484, *p*=0.694; sex*rank [wait zone]: *F*=3.689, *p*<0.05, **Fig. 4d-f**). Offer zone thresholds on the other hand remained equal in all groups and gradually decreased across a much slower timescale over several weeks (**Fig. 4d-f**). LINC00473 expression in mPFC did not significantly alter any decision-making thresholds (sex*virus [offer zone]: *F*=1.781, *p*=0.185; sex*virus [wait zone]: *F*=187, *p*=0.175, **Fig. 4d-f**).

In order to determine which groups’ decision policies were most optimal, we leveraged an economic analysis we recently developed that quantifies how mice maximize food and subjective value ^43^. This analysis takes each animal’s offer zone and wait zone thresholds observed on each day and calculates the Euclidean distance between these observed thresholds and theoretical thresholds that yield either the most amount of food or most amount of subjective value (see **Methods** and **Supplementary Fig. 7**). In the 1-30 s epoch, we empirically determined through computer simulations that a threshold of 10 s in any restaurant yields the most amount of food ^43^. Observed thresholds that are either above or below this theoretical threshold thus are suboptimal, in the food-maximizing domain. To determine theoretical thresholds that maximize subjective value, we used individual differences in flavor-specific earnings on day 7 when all costs were 1 s only ^43^. This allowed us to determine idealized thresholds in each restaurant in the 1-30 s epoch that re-approximates on a mouse-by-mouse basis day 7’s ratio of earnings across flavors ^43^. Therefore, observed thresholds that are either above or below this theoretical threshold thus are suboptimal, in the subjective value-maximizing domain. Euclidean distances between observed and theoretical thresholds were calculated separately in the offer zone and wait zone using a 4-dimensional vector of thresholds comprising each restaurant [chocolate, banana, grape, plain].

We found that female decision-making policies in the final epoch of testing were more optimized for both maximizing total food intake as well as maximizing subjective value compared to males (**Fig. 4g-i**, **Supplementary Fig. 7**). This sex-dependent effect was specific to wait zone but not offer zone decision-making policies. LINC00473 treatment had no effect on Euclidean distances. These data highlight a robust sex difference in the optimality of wait zone decision-making policies unique from those in the offer zone ^16,57^. The fact that thresholds are not altered by LINC00473 rules out major confounds of baseline willingness to enter or willingness to wait on higher-order change-of-mind decision-making valuations investigated further below.

### LINC00473 alters the temporal dynamics of change-of-mind decision-making behavior

Wait zone thresholds capture economic indifference points of re-evaluation decisions when deciding to stay or quit. We defined these measures by choice outcome as a function of the offer cost that was cued at trial start, that is, at the onset of the countdown during the delay period. Next, we quantified more dynamically at what timepoint quit decisions were made during the delay period in the wait zone. We calculated the value of each offer as well as the value remaining in the countdown at the moment of quitting. We subtracted the offer or countdown time remaining from an individual mouse’s wait zone threshold in each restaurant (example trial: 18 s offer entered, mouse’s threshold = 10 s, **offer value** = 10 - 18 = −8, latency to quit = 4 s, time remaining at quit = 14 s, **value remaining** at quit = 10 - 14 = **−4**, **Fig. 5a**, **Supplementary Fig. 8**). We found all mice generally made quit decisions for accepted offers that were (i) typically above one’s wait zone threshold (i.e., negatively valued offers) and (ii) had ample time remaining in the countdown such that the remaining time investment required to obtain the reward was still above one’s threshold (i.e., negative value left, **Fig. 5a**). Therefore, most quit decisions are economically advantageous and are efficiently time-saving (**Fig. 5a**). Furthermore, quit decisions are corrective re-evaluations of sub-optimal and impulsive offer zone decisions, consistent with previous reports ^5,16,43,56,58^. Importantly, groups did not differ in how they made enter vs. skip decisions in the offer zone (**Supplementary Fig. 9**).

**Fig. 5.**
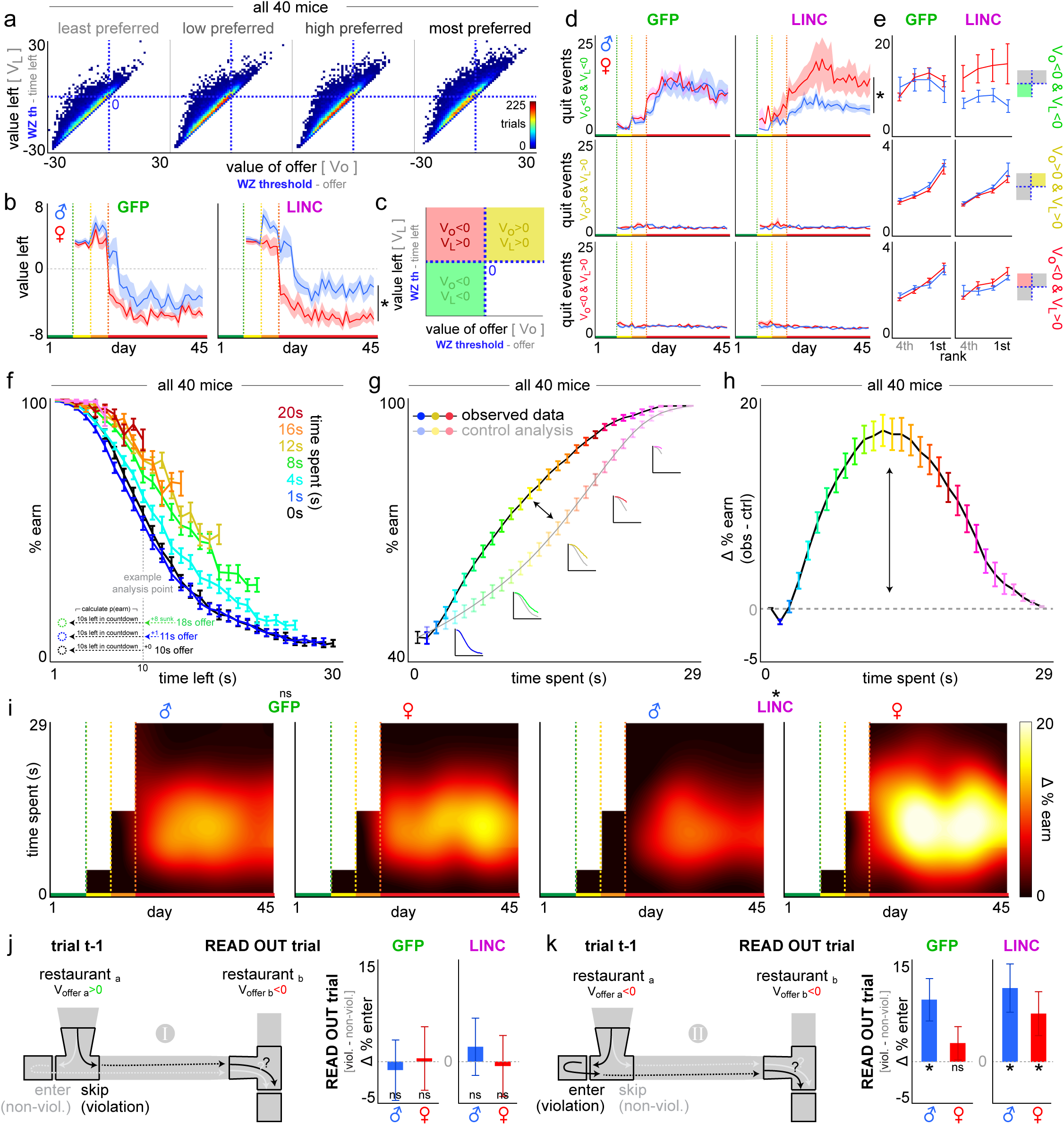
LINC00473 expression in mPFC alters the temporal dynamics of change-of-mind decisions. **a** Distribution of quit trials by offer value or value left from the 1-30 s epoch. **b** Average value left at quit. **c** Three categories of quit events based on offer value and value left. **d** Number of quit events from each quit category in (c). **e** Split by restaurants ranked from least (4^th^) to most preferred (1^st^) from the 1-30 s epoch. **f-h** Sunk cost analysis of staying behavior in the wait zone. (f) Likelihood of staying in the wait zone as a function of time left in the countdown and time already spent waiting. Black 0 s time spent represents just entering the wait zone. Inset llustrates an example analysis point matched at 10 s left. Data from (f) dimensioned reduced in (g) collapsing across time left. Insets in (g) depict data from curves in (f) collapsed into the observed (sunk condition) and control (0 s condition) lines. Difference between curves in (g) are plotted in (h) to summarize the envelope of the overall effect of time already spent on staying in the wait zone. **i** Sensitivity to sunk costs across the entire paradigm. **j-k** Regret-related sequence analyses comparing trial pairs between (j) type 1 scenarios and (k) type 2 scenarios. Read out trials captures a change in behavior depending on choice history of distinct economic violations (black) compared to non-violations (gray) on trial t-1. See **Methods** for a full explanation of these analyses. Prominent sex-dependent effect revealed by LINC00473 expression in mPFC on change-of-mind related sunk costs (enhanced in females) and regret sensitivity (type 2: enhanced in females). **\*** in (b-i) represent significant sex differences. * in (j-k) represent delta score significantly different from 0. Shading / error bars represent ±1 SEM.

We found that females made quit decisions in the wait zone with significantly lower and more negative value remaining in the countdown compared to males (sex*virus: *F*=195.823, *p*<0.0001, **Fig. 5b**). This sex difference was more pronounced in LINC00473-treated animals (sex*virus: *F*=12.957, *p*<0.001, **Fig. 5b**). Furthermore, only LINC0047-treated mice revealed a sex difference in the number of quit decisions that could be characterized as economically advantageous (as described above, sex*virus: *F*=153.236, *p*<0.0001). There was a greater proportion of these events in females relative to males and compared to other quit events that were less common and more disadvantageous (i.e., (i) quitting negatively valued offers [above one’s threshold] with positive value left [below one’s threshold]; or, (ii) quitting positively valued offers [below one’s threshold]; both (i) and (ii) reflect quits passing a timepoint after which mice should have chosen to stay instead, quit type*sex*virus: *F*=13.293, *p*<0.0001, **Fig. 5c-d**, **Supplementary Fig. 8**). Overall, these data indicate that LINC00473-treated females engage in more economically advantageous change-of-mind decisions compared to males that depend on the temporal dynamics and efficiency of quitting behavior.

Next, we characterized sensitivity to sunk costs while mice made continuous re-evaluation decisions in the wait zone as a function of the passage of time (which wait zone thresholds do not capture). We calculated the instantaneous probability of staying and earning a reward in the wait zone as a continuous function of not only (i) the time remaining in the countdown at each snapshot in time but also (ii) the time already spent waiting ^18^. We computed a survival analysis using a sliding 1 s window after binning all trials of varying starting offers into [*time spent*, *time left*] pairs (**Fig. 5f**). We then calculated the probability of earning a reward as a function of time already spent waiting independent of temporal distance to the goal (see **Methods** and **Supplementary Fig. 8** for a detailed and graphical description of sunk cost analyses and control analyses).

In all groups, we found that the passage of time drove an increased likelihood that mice would continue to wait – an escalation of commitment – which grew additively stronger with more time spent (time spent*time left: *F*=891.924, *p*<0.0001, **Fig. 5f-h**). This effect was independent of and orthogonal to temporal distance to the goal measured at any given moment in time. We found a significant sex difference in sensitivity to the amount of time already spent waiting only in LINC00473-treated mice. LINC00473-treated females demonstrated increased sensitivity to sunk costs, while LINC00473-treated males instead demonstrated relatively diminished sensitivity (sex*virus: *F*=15.944, *p*<0.0001, **Fig. 5i**). Sunk cost sensitivity only emerged after mice transitioned to the 1-30 s epoch (time spent in wait zone [1-15 s offers]: *F*=0.174, *p*=0.677, **Fig. 5i**, **Supplementary Fig. 10**). Sunk cost sensitivity uniquely affected wait zone decisions compared to other forms of time spent on this task, including in the offer zone (time spent in offer zone: *F*=0.558, *p*=0.455; time spent*sex*virus: *F*-0.418, *p*-0.518, **Supplementary Fig. 10**, see **Supplementary Text** and **Supplementary Fig. 11–13** for manipulations of task-relevant sensory information).

These data describe how LINC00473-treated females are more sensitive to the passage of time when re-evaluating the decision to quit. This captures a critical duplexity: although LINC00473-treated females are generally more likely to engage in rapid, economically advantageous quit decisions, they also change in this regard more dynamically with the passage of time. That is, LINC00473-treated females cache more value with time spent when deciding to quit that promotes overstaying behavior. This reflects a steepened value function during the re-evaluation process that is capable of overturning quit choices during change-of-mind decision-making, enhanced by mPFC LINC00473 expression in females.

### LINC00473 alters the impact of prior change-of-mind decisions on future behavior

Next, we aimed to characterize how sequences of events surrounding quit decisions influenced subsequent behavior. Because trials are interdependent, a strength of the Restaurant Row foraging task lies in being able to subdivide trial sequences into unique economic scenarios in which animals naturalistically find themselves. We identified trial pair sequences [trial t-1 → trial t] in which animals made atypical, economically suboptimal choices [on trial t-1] and measured the change in economic choice behavior on the next trial [on trial t: the read-out trial] ^5,16,17^. These events were compared to specific alternative sequences that control several additional variables on both trial t-1 and trial t in order to rule out other confounds and isolate the economic impact of choice history on future behavior: (i) the value of the offer presented, (ii) the flavor identity, and (iii) the previous action selected. Sequences were extracted post hoc from natural encounters while foraging amid randomly presented offers and not constructed a priori nor built into the task design. This allowed us to investigate how past economic violations alter future choices as a behavioral readout of sensitivity to regret, as previously demonstrated ^5,16,17^.

We previously defined two distinct sequences of events on this task that each capture the impact of decision history on future choices but differ in the specific actions that give rise to two types of economic violations ^5,16,17^. **Scenario type 1**: situations in which animals inappropriately forgo a high-value offer [on trial t-1] only to subsequently encounter a worse low-value offer on the next trial [trial t] (**Fig. 5j**). This can be classified as a violation sequence and can be compared to the non-violation control sequence in which mice instead appropriately accepted the high-value offer on trial t-1, holding all other variables constant (**Fig. 5j**). Here, we found that all groups were behaviorally insensitive to scenario type 1, with no effect of sex or LINC00473 expression (+ sign test: [male GFP]: *t*=−0.282, *p*=0.608; [female GFP]: *t*=0.101, *p*=0.461; [male LINC]: *t*=0.521, *p*=0.308; [female LINC]: *t*=−0.144, *p*=0.556, **Fig. 5j**). That is, there were no changes in economic choice behavior on the read-out trial when comparing violation sequences to non-violation sequences [thus yielding a delta score of zero for choice probabilities on trial t] (**Fig. 5j**).

**Scenario type 2**: situations in which animals inappropriately accepted a low-value offer [on trial t-1] (**Fig. 5k**). Note that these scenarios typically result in a change-of-mind decision once in the wait zone. This violation sequence can be compared to the non-violation control sequence in which mice instead appropriately skipped the low-value offer on trial t-1, holding all other variables constant and matched for the same low-value offer on trial t as in scenario type 1 (**Fig. 5j-k**). Overall, we found that mice demonstrated behavioral sensitivity to scenario type 2, whereby individuals were more likely to compensate for and accept low-value offers that they typically would not accept on the read-out trial following violation sequences compared to non-violation sequences (**Fig. 5k**). Interestingly, only GFP-treated females did not display behavioral sensitivity to scenario type 2, while all other groups including LINC00473-treated females displayed delta scores significantly greater than zero (+ sign test: [male GFP]: *t*=2.931, *p*<0.01; [female GFP]: *t*=1.123, *p*=0.145; [male LINC]: *t*=3.056, *p*<0.01; [female LINC]: *t*=2.196, *p*<0.05, **Fig. 5k**).

These data reveal how mice can differentially carry the weight of distinct types of past mistakes into future choices. Remarkably, there is a unique enhancement of behavioral sensitivity to type 2 but not type 1 scenarios caused by mPFC LINC00473 treatment in females only. These events depend on change-of-mind decisions on trial t-1 and derive from the same trial pool from which sensitivity to sunk costs is measured, and importantly, enhanced in LINC00473-treated females. Taken together, this suggests a common underlying computational process may be shared between change-of-mind-related regret and sunk cost sensitivity that is engaged by mPFC LINC00473 expression in females, resulting in a steepened value function shared across both processes. Furthermore, these findings provide an example whereby LINC00473 expression in this brain region overcomes a baseline sex difference that is potentially disadvantageous for females. This is especially important considering (i) LINC00473 expression in mPFC carries pro-resilient properties in females and (ii) traits associated with enhanced sensitivity to type 2 regret and sunk costs have been previously linked to stress-resilient individuals ^16,22,43^.

In summary, this collection of findings demonstrates the complexity of change-of-mind decisions that respond to the economic landscape of the environment. This work showcases the multifactorial ways in which re-evaluative choice processes can be uniquely modulated in a sex-dependent manner. Importantly, we highlight a distinction between separate phases along a continuous decision stream captured within the same trial: primary offer zone judgements followed by secondary wait zone re-evaluative decisions. We reveal striking sex differences and sex-dependent effects of LINC00473 expression in mPFC on only wait zone decisions. We provide evidence for LINC00473-induced changes in signatures of two different yet related higher-order decision-making phenomena during quit decisions: sensitivity to the passage of time in the form of sunk costs and change-of-mind related regret – valuation algorithms that comprise critical aspects counterfactual thinking.

## Discussion

In this study, we characterized complex behaviors in mice tested on the neuroeconomic foraging paradigm, Restaurant Row. We explored the role of a non-coding RNA, a relatively poorly understood class of molecules, on motivated behavior. We discovered sex-dependent effects of LINC00473 expression in mPFC neurons on specific aspects of value-based decision-making behavior. This was typified by selectively altering the way in which an individual changes one’s mind during an ongoing re-evaluative decision without affecting primary evaluations. Importantly, we discovered a role for mPFC LINC00473 expression – which we recently found increases stress resilience in simple behavioral screening tests in female mice only ^22^ – in enhancing the contrast of re-evaluation processes in females. Our findings extend from simple conditioned place preference behaviors like those commonly assayed using simpler goal-oriented paradigms ^50,67,68^ to more complex, higher-order neuroeconomic phenomena. These findings build upon our recent work characterizing how complex decision-making processes go awry in depression and interact with molecular regulators of stress resilience ^16^. Here, we provide a deeper understanding of how multiple stages of a continuous decision stream derive from dissociable psychological mechanisms. We explored how perturbations in change-of-mind decision-making could underlie select elements of negative rumination. Our findings have important implications for how molecular determinants of stress-sensitive decision-making processes interact with non-coding regions of the genome differently in the male and female brain.

The distinction between offer zone choices and wait zone choices on the Restaurant Row task is critically important. They reflect a separation of fundamentally distinct decision-making processes that occur within the same trial but are segregated across space and time ^58^. Numerous reports have tested subjects on this paradigm translated for use across species in task variants adapted for mice, rats, monkeys, and humans ^18,51,52^. These studies have shed light on the biological and psychological implications of resolving dissociable, evolutionarily conserved decision-making processes. In the offer zone, individuals engage deliberative systems that represent alternating options of competing actions between which one must choose ^61,62^. This has been shown in rodents to derive in part from hippocampal (HPC)-dependent processes ^57,58^. Population-level activity of neurons in HPC can encode multiple, alternating future paths through the offer zone to potential goals in the current vs. next restaurant ^65,67,69,70^. Similar findings have been observed from fMRI studies in humans tested on a translated version of the Restaurant Row task ^55^. Default mode network activation patterns can encode competing future options during offer zone deliberation, but seldom do such representations include past events or previous restaurants ^2,71^.

Decisions in the wait zone on the other hand are much less understood. The HPC has been implicated in wait zone decisions as well: it switches oscillatory activity from a predominately theta-driven state sensitive to the cued offer costs in the offer zone ^61^ instead to an increase in sharp wave ripple activity sensitive to time spent waiting in the wait zone ^62^. Further, wait zone decision periods also depend on HPC synchrony with the mPFC ^61,62^. Recordings in rodents from the mPFC on Restaurant Row surprisingly found no evidence of encoding of the cued offer costs when animals were in the offer zone ^59^. However, once in the wait zone, mPFC neurons strongly represented offer value independent of time spent waiting and could predict whether or not the trial would terminate in a quit ^59^. Within the mPFC, pairs of neurons also drastically changed excitation-inhibition balance specifically when animals were in the wait zone ^72^. Taken together, these findings indicate a qualitative shift in mental state upon transition from the offer zone to the wait zone that recruits the mPFC preferentially during change-of-mind decisions.

This study represents the first analysis of females in the Restaurant Row task. Nearly all of the sex differences captured here, including those altered by mPFC LINC00473 expression, are restricted to wait zone and not offer zone behaviors (**Fig. 6**, **Supplementary Fig. 14**). Importantly, we found no effects of sex or LINC00473 on how impulsive or deliberative animals behaved in the offer zone. This includes no changes in vicarious trial and error behavior, which is known to correlate with neural representations of episodic future thinking and model-based planning in HPC among other structures ^73^. Recently, we showed that LINC00473 expression diminishes spontaneous excitatory post-synaptic current frequency and amplitude in excitatory mPFC pyramidal neurons of female mice, with no effect seen in males ^22^. Chemogenetic inactivation of the mPFC during Restaurant Row diminishes wait zone quit decisions ^60–62^. Optogenetic induction of long-term depotentiation in excitatory mPFC neurons that project to the nucleus accumbens (NAc) ^57^, where negative dopamine reward prediction error signals have been found to causally precede quit choices on Restaurant Row ^64^, selectively decreases quitting behavior for several days without altering offer zone behavior ^57^. Outside of Restaurant Row, intracranial optogenetic excitatory self-stimulation of mPFC terminals in the NAc can be reinforcing while also promoting enter-then-quitting behavior ^74^. These converging findings suggest that LINC00473 expression is likely mediating sex-dependent effects on change-of-mind decisions by at least in part dampening and restructuring mPFC activity and connectivity with other regions. In depressed patients, neuromodulation interventions that decrease mPFC activity yield therapeutic benefits in mood ^75^. Thus, understanding how these decision-making changes come together can help bridge interpretations between the behavioral and neural computations underlying phenomena like negative rumination and the molecular drivers of stress responses traits.

**Fig. 6.**
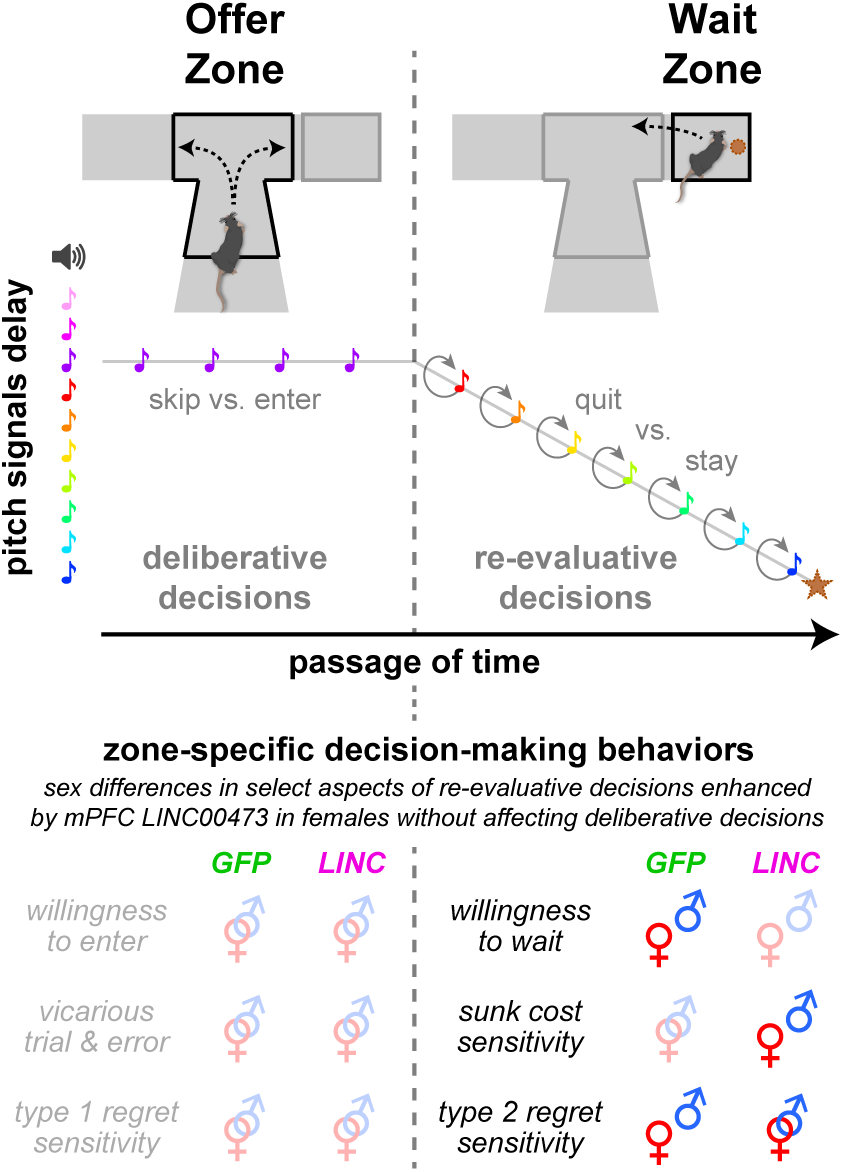
Summary of key findings. Distinction between decision-making processes measured within trial in the offer zone (deliberative choices between skipping vs. entering) and wait zone (re-evaluative choices between quitting vs. continuing to wait before earning a reward, star symbol) as a function of cued delay costs (tone pitch, colored notes) and subjective value (flavor). Analysis of foraging behaviors reveal signatures of dissociable neuroeconomic processes in the offer zone (willingness to enter, vicarious trial and error choice behavior, behavioral sensitivity to type 1 regret scenarios: the impact of skipping high-value offers on subsequent choices) and wait zone (willingness to wait, behavioral sensitivity to sunk costs, and behavioral sensitivity to type 2 regret scenarios: the impact of entering and then quitting low-value offers on subsequent choices). Interlocking male and female symbols (⚤) represent no sex differences while separated symbols (♀ ♂) indicate sex differences, either at baseline (GFP group) or altered by LINC00473 expression in mPFC. See **Supplementary Fig. 14** for an expanded summary table.

A key insight from the present study is a conceptual advance toward developing neuroeconomic principles unifying two critical features of change-of-mind decision-making: sensitivity to sunk costs and regret. Sensitivity to sunk costs captures a well-studied cognitive bias whereby irrecoverable past expenditures can enhance the value of continuing to invest in and perseverate on an ongoing endeavor instead of cutting losses, as we have reported previously captured across species ^18,56^. Sensitivity to regret describes how representations of missed opportunities and alternative actions that could have led to better outcomes can influence future choices ^5,6,16,17^. We theorize that these two phenomena share, at least in some respects, a common computational origin (**Fig. 7**). We previously characterized – behaviorally and neurophysiologically – distinct forms of regret based on two types of economic scenarios in which individuals find themselves ^5,16,17^. Type 1 scenarios describe economically risky decisions to skip high-value offers, followed by poor outcomes. Type 2 scenarios describe economically poor decisions to enter and begin investing time in low-value offers before deciding to quit. Counterfactual thinking during type 2 scenarios likely evokes “should have skipped instead” representations that drive change-of-mind decisions. We posit that sensitivity to sunk costs, which derive from these same type 2 scenarios, is a parallel valuation that stems from the same counterfactual process as type 2 regret and may compete with the decision to quit. With the passage of time, this can promote overstaying behavior and drive sensitivity to sunk costs. However, if an individual does quit, this can also drive altered valuations on the subsequent trial and promote compensatory behavior, measured as type 2 regret sensitivity. Here, we provide evidence that both of these processes – sensitivity to sunk costs and type 2 regret – can be experimentally enhanced in LINC00473-treated females without affecting type 1 regret sensitivity or other lower-order decision processes such as willingness to wait. This suggests that a value function of a common counterfactual process is being selectively enhanced in these animals (**Fig. 7**). Within this conceptual framework, we posit that features of negative rumination could stem from a shared neural substrate governed by these two neuroeconomic processes and may be associated with stress-related traits of the individual.

**Fig. 7.**
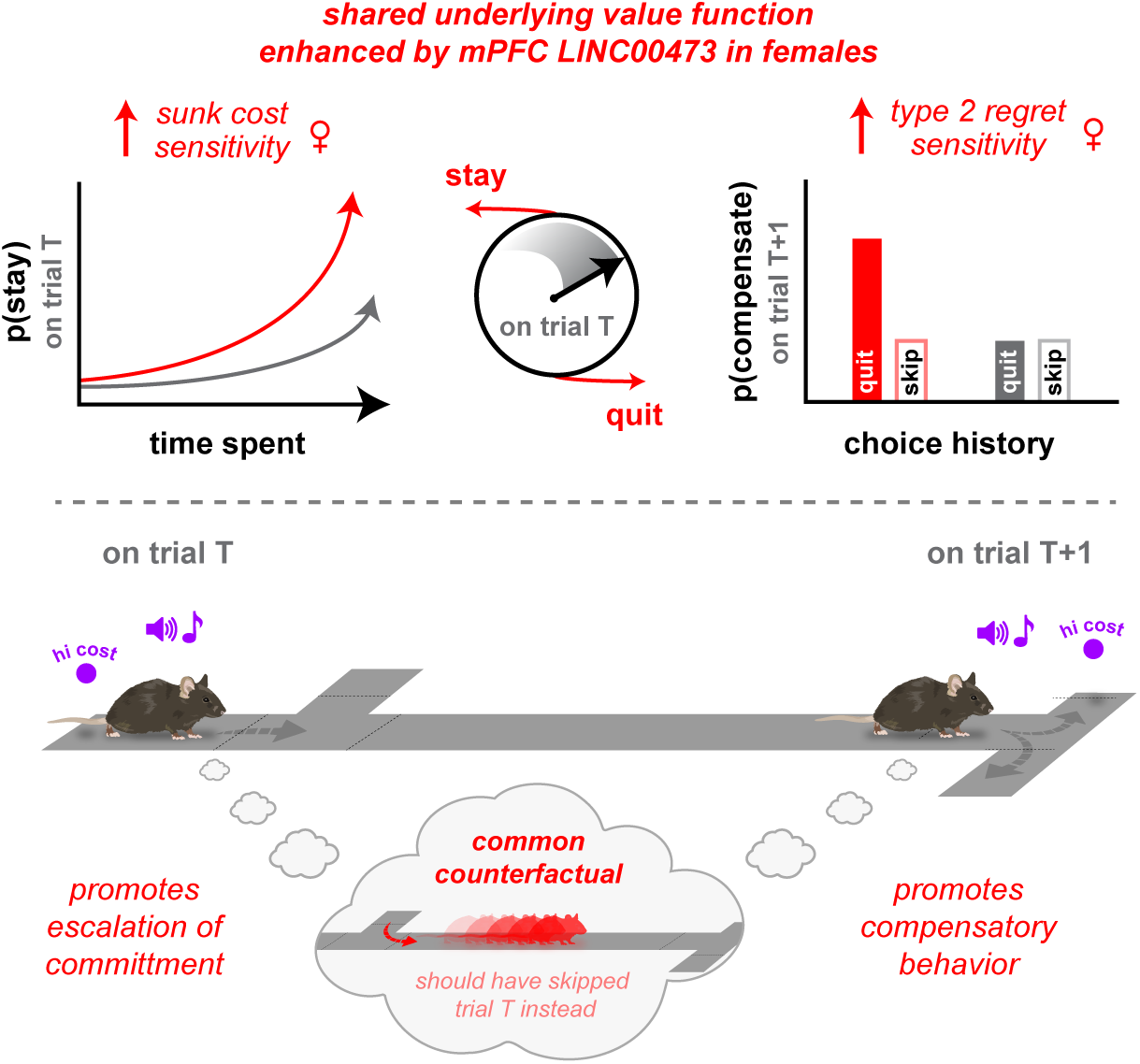
Conceptual neuroeconomic framework. Working model of the link between key wait zone phenotypes enhanced in LINC00473-treated females during change-of-mind decision-making. This schematic depicts the choice conflict after mice accept a high-cost offer – an economic violation in the offer zone – once they arrive in the wait zone and are faced with competing actions to stay vs. quit. Clock emphasizes that this decision process evolves with the passage of time. Quitting rapidly is most economically advantageous. Yet, accruing sunk costs promotes staying as added value is cached over time. If these trials are quit – a defining feature of type 2 regret scenarios – this added value is carried into subsequent trials, augmenting future choices. Red arrows highlight enhanced sensitivity to sunk costs and type 2 regret apparent in LINC00473-treated females compared to baseline (gray). Sunk costs: p(stay) increases as a function of time spent waiting on trial T; Type 2 regret: p(compensate) [increased likelihood of accepting offers typically rejected on trial T+1] is higher following violation sequences [enter-then-quit] relative to non-violation control sequences [skip] for high-cost offers on trial T. Proposed unifying principle linking sensitivity to sunk costs and type 2 regret: Both phenomena stem from the same trial pool on trial T: inappropriately accepting a high-cost offer. Alterations in reward value, driving either sunk cost-induced staying or post-quit regret-related compensatory behavior, may stem from a shared value function. There may be a common underlying counterfactual process where the unselected alternative (i.e., skipping trial T instead) carries added value and influences behavior either during staying or after quitting. This value function in either case minimizes future loss. This work makes explicit predictions about hypotheses for future research investigating the neural correlates of shared counterfactual representations and, clinically, if the content of negative rumination in mood disorders maps onto these representations.

To put this in context, we recently characterized the impact of chronic social defeat stress on Restaurant Row behavior in male mice, in tandem with disruptions of CREB function in mPFC ^16,43,76^. First, we found that mice were required to progress through worsening economic challenges in order to extract any group differences in foraging profiles ^43^. This is consistent with results presented here that emphasize the importance of considering the environmental demand and stressors of a task necessary to unmask disease-relevant phenotypes. Second, we found that stress-resilient mice – defined by a simple social interaction screening test measured outside of the Restaurant Row task – mounted the most robust behavioral response to the transition into the 1-30 s reward-scarce environment in order to recover food earnings, similar to LINC00473-treated female mice as shown here ^43^. Third, without grossly altering decision-making thresholds, we found striking stress-induced changes in sensitivity to sunk costs and regret. Stress-susceptible mice demonstrated increased regret sensitivity to type 1 scenarios. This was a phenotype no group demonstrated in the present study ^16^. Additionally, stress-susceptible mice demonstrated diminished regret sensitivity to type 2 scenarios. Furthermore, stress-resilient individuals instead displayed enhanced sensitivity to type 2 scenarios as well as enhanced sensitivity to sunk costs ^16^. Taken together, this suggests that the females in the present study at baseline may have a decision-making vulnerability specifically in change-of-mind choice processes that is enhanced or restored to male levels by LINC00473 expression in mPFC.

In our prior study, knocking down CREB function had multifactorial effects on decision-making behavior depending on the brain region targeted (mPFC vs. NAc) and whether or not animals were stress-naïve at baseline vs. exposed to subsequent stress ^16^. CREB function in the mPFC was required to maintain suppression of regret sensitivity to type 1 scenarios and, when knocked down, rendered only type 2 scenarios sensitive to subsequent stress. This pattern was distinct from CREB function in the NAc where it is known to drive stress susceptibility ^16,29,77,78^. Importantly, CREB function in mPFC altered not only re-evaluative decisions in the wait zone but also primary decisions in the offer zone, including affecting impulsive choices as well as deliberative vicarious trial and error behavior ^16^, unlike the effects of LINC00473 here which were restricted to wait zone decisions only and did not alter type 1 regret sensitivity.

These data implicate a divergence in how CREB and LINC00473 interact within the mPFC to drive distinct aspects of decision-making behavior. This allows for a new perspective to interpret prior findings in stress research but also opens new questions for future lines of investigation. For instance, it is possible that CREB and LINC00473 effects on neuronal function may diverge in different cell types or sub-populations of neurons that receive different inputs or project to different downstream regions. Prior work implicates involvement of HPC and NAc as two key regions that could be mediating this difference. Sex-dependent effects of LINC00473 also suggest that complex interactions between sexually dimorphic circuits and sex hormones may be critically important in determining how mPFC neurons encodes time, value, and choice. Quantifying similarities in neural signals during the processing of sunk costs vs. distinct regret types remains an open question, with explicit predictions we make here about similarities between sunk costs and type 2 regret. Finally, it is unknown if the representations during the processing of sunk costs or change-of-mind regret overlap with the sort of content patients with depression ruminate over and what features of counterfactual thinking, if any, promote stress-resilience in humans.

Limitations of the current student include that LINC00473’s sex-dependent effects on behavior have only been causally tested in the mPFC to date ^22^. Further, prior CREB knockdown studies during Restaurant Row were conducted in males ^16,43,76^. Although postmortem human tissue studies implicate mPFC as a top region to investigate ^22^, understanding LINC00473’s role in other regions, especially in how it may be interacting with other key molecular players such as CREB in the NAc ^26,29,30^, will be critically important in future studies investigating multiple networks. For instance, LINC00473 has been shown to increase cellular excitability in cultured HPC neurons ^26^, contrary to its effects in mPFC ^22^. It will be critical to record *in vivo* neural activity from these structures in both males and females with and without perturbations of LINC00473 alone, CREB alone, and both, in order to characterize the interplay between these overlapping molecular pathways. Future decision-making studies should combine experiments of LINC00473 manipulations with exposure to other stressors, beyond the economic challenges tested in the present study, such as chronic social defeat or variable stress on decision-making ^22^. Additional work needs to be done investigating the role of gonadal steroid hormones, sex chromosomes, and sex-dependent noncoding RNAs in mediating aspects of these reported sex differences. As just one example, CREB function has been shown to fluctuate with estrus cycle ^79–83^. Additionally, given previous reports that LINC00473 expression in mPFC of stress-naïve male mice can induce behavioral changes in simpler assays (i.e., marble burying) ^22^, it would be fruitful to explore in future work how Restaurant Row decision-making profiles could relate to traits outside of the task itself (e.g., anxiety-related behavior measured on independent tests).

We discovered robust sex differences in neuroeconomic change-of-mind decision-making. We found that LINC00473 expression in the mPFC of females gave rise to enhanced behavioral sensitivity to sunk costs and change-of-mind related regret. These effects on secondary re-evaluative choices were independent of primary decisions. Counterfactual thinking during change-of-mind decisions can elicit “should not have done that” mental representations that map hypothetical outcomes onto alternative actions that could have been selected instead. We propose that the processing of sunk costs and change-of-mind related regret may depend on a common value function and shared form of counterfactual thinking. Such processes could guide aspects of negative rumination and impact mood as well as stress-resilience. However, even if change-of-mind decisions are unpleasant or evoke some degree of cognitive dissonance, enhancing the salience of this process could increase attention paid toward realized losses and carry utility for fictive learning. Collectively, variations in these decision-making patterns are sex-dependent and may alter the lens through which individuals process their past mistakes.

## Methods

### Subjects

Adult male and female C57BL/6J mice (Jackson, 10wks, n=40) were used. Mice were individually housed and maintained on a 12-hr light/dark cycle with ad libitum water, only food-restricted during Restaurant Row testing, conducted during their light phase. Experiments were approved by the Mount Sinai Institutional Animal Care and Use Committee (IACUC; protocol number LA12-00051) and adhered to the National Institutes of Health (NIH) guidelines.

### Viral constructs

To achieve long-term neuronal expression of LINC00473, it was subcloned into an AAV plasmid with human synapsin promoter (hSyn) driving gene expression. LINC00473 transcript variant 1, (NR_026860.1 /AK289375) of 1821bp length was subcloned from p1005+ HSV vector ^22^ into pAAV-hSyn-eGFP-hSny plasmid ^83^ using blunt ligation into a PMEI and EcoRV restriction sites following the second hSyn promoter. The orientation and sequence were validated using Sanger sequencing. Empty plasmid expressing eGFP only was used as control. These vectors were used to generate AAV serotype 2 at the University of Maryland virus vector core. Viral vector placement and expression were confirmed in vivo by a GFP fluorescent flashlight and qPCR.

### RNA extraction, RT, and qPCR

Total RNA was isolated and purified with RNeasy Micro Kit (Qiagen), using QIAzol as a lysis reagent, including DNase treatment. All samples were tested for concentration and purity using a NanoDrop (Thermo Fisher). RNA amount was normalized across samples (400ng), and cDNA was created using iScript (Bio-Rad). Real-time PCR reactions were run in triplicate using PowerUp SYBR Master Mix (Applied Biosystems) in a QuantStudio 7 (Thermo Fisher) qPCR machine. Primer sequences: LINC00473: forward: GCATACTTTGGCGGACCTTTT, reverse TGTGCCTCCCTGTGAATTCTC; HPRT1 forward: GCAGTACAGCCCCAAAATGG, reverse: GGTCCTTTTCACCAGCAAGCT; GFP forward: CATGCCCGAAGGCTACGT, reverse CGATGCCCTTCAGCTCGAT. Raw Ct values are presented and not normalized amounts, as LINC00473 is not endogenously expressed in mice.

### Virus surgery

Mice were anesthetized by intraperitoneal injection with a mixture of Ketamine HCl (100 mg/kg) and Xylazine (10 mg/kg) and positioned on a stereotaxic instrument (David Kopf Instruments). In the medial prefrontal cortex (mPFC, from bregma with an angle of 15 degrees: AP +1.8 mm; ML ±0.75 mm; DV −2.7 mm), 0.7 μL of virus (AAV2-hSyn-GFP or AAV2-hSyn-LINC00473-GFP) was bilaterally infused using 33-Gauge Hamilton needles over 5 min at a rate of 0.1 μL/min, and the needle was left in place for 10-15 min after the injection before retraction. Mice were allotted 2 weeks to recover before beginning food restriction across 3-4 days in preparation for testing on Restaurant Row. An initial virus validation cohort (independent cohort of n=10 mice) was injected with GFP or LINC00473 viruses bilaterally in mPFC at a titer of 1×10^12^ at. These animals were tested on Restaurant Row for only 1 week (offers remained at 1 s only, first epoch, green) after which they were sacrificed for PCR validation on tissue punches obtained from mPFC. Behavioral data from these animals on the conditioned-place preference probe (days 0 and 8) appear in **Supplementary Fig. 4**. A second replication cohort (n=40 mice) was injected with GFP or LINC00473 viruses bilaterally in mPFC at a titer of 5×10^12^ and served as the primary cohort of animals for the main figures that underwent the full 45-day longitudinal behavioral testing. Brain tissue used for immunofluorescence histological visualization of virus transfection was collected in a separate set of mice three weeks post-surgery. At time of collection, animals were deeply anesthetized with peritoneal injections of 500 mg/kg of Fatal Plus (Vortech, Cat #9373), and intracardially perfused with 15 mL 4% PFA (Electron Microscopy Science, Cat #15713-S). Brains were postfixed for 24-72 hr, and subsequently sliced on a Leica VT1000 S vibratome at 40-60 µM sections. Sections were blocked for 1 hr in blocking buffer (10% donkey serum (Jackson Immunoresearch, Cat #017-000-121), 0.3% Triton-X (Sigma, Cat #9284) in PBS), followed by overnight incubation with primary antibody (1:1000 Ch-NeuN, MilliporeSigma, Cat #ABN91) in diluted blocking buffer (1:3 dilution in PBS). Sections were washed three times with diluted blocking buffer (15 min each) before incubation with secondary antibodies (Ch-647: Jackson Immunoresearch, Cat# 703-605-155) for 1 hr. Two additional 15-min washes in diluted blocking buffer and one 15-min wash in PBS. Finally, sections were incubated with 1:10,000 DAPI (Thermofisher, Cat #62248) for 5 min. Sections were mounted with EcoMount (Biocare Medical, Cat #EM897L). Images were acquired on a Zeiss LSM 900 confocal microscope using Zen Blue software with 10x and 40x oil immersion lens at 1:1 digital zoom.

### Neuroeconomic Task

Following surgery, we characterized mice longitudinally in Restaurant Row ^5^. On this task, mice had a limited time period each day to forage for food by navigating a maze with four uniquely flavored (chocolate, banana, grape, and plain) and contextualized (horizontal stripes, dots, triangles, vertical stripes) feeding sites, or “restaurants” (Fig. 1b). Animals were allotted 30 min to forage for their primary source of food for the day. Rewards used on this task were obtained from BioServe (20 mg pellets #F007120). These pellets are full nutrition and not sucrose-only pellets that only differed in flavoring but not nutritional profile (3.6 kcal/gm calories, 59.1% carbohydrate, 18.7% protein, 6.5% ash, 5.6% fat, 4.7% fiber). Choices on this task were interdependent across trials and across days. This makes the task economic in nature, requiring animals to budget their limited time effectively in a self-paced manner in order to earn a sufficient amount of food in a closed-economy system. The cost of earning a reward took place in the form of a delay animals would have to wait cued by the pitch of a tone that sounded upon entry into each restaurant. Each restaurant had a separate offer zone (OZ) and wait zone (WZ). Upon entry into the OZ, a tone (500 ms) sounded whose pitch indicated how long of a delay mice would have to wait in a cued countdown should they enter the WZ. Mice could decide to skip an offer presented in the OZ and proceed down the hallway to the next restaurant. In the OZ, tones repeated every second held at the same pitch to repeatedly cue the offer of the current trial until mice either made a skip or enter decision. If mice decide instead to enter the WZ, this triggered a countdown with descending tone pitch stepping every second until mice either waited out the full delay and earned a food pellet or quit prematurely during the countdown. If mice made a skip or quit decision, the tone terminated, the offer was rescinded, and mice must proceed forward to the next restaurant. Mice quickly learned to only travel in one direction as animals must encounter offers serially navigating the maze in a counterclockwise direction (i.e., nothing happens if mice traverse in the wrong direction other than wasting time). Mice were tested across 45 consecutive days for the main experiment where strategies developed longitudinally as animals learned the structure of the task and the changing economic landscape.

During the first week of testing (block 1, green epoch), all trials consisted of 1 s offers only. 1 s offers were indicated by tones played at the 4,000 Hz pitch. A special probe session was inserted into this timeline immediately before (day 0) and after (day 8) the first week of testing during which mice essentially were allowed to roam the empty maze arena freely without any task components (no tones, no rewards) to examine conditioned place preference behaviors learned through the natural experience of the first week of the task itself. The remainder of the experimental timeline continued thereafter. During the next 5 days of testing (block 2, yellow epoch), offers ranged from 1 to 5 s (4,000 Hz to 5,548 Hz in 387 Hz steps) randomly selected from a uniform distribution. For instance, if a 4 s offer was selected on a given trial, the tone presented in the OZ would be 5,161 Hz for 500 ms, and this would repeat every second indefinitely until a skip or enter decision was made. If the WZ was entered, using this example trial, this 5,161 Hz tone would step down by 387 Hz every second descending in pitch until earning (with the final tone of 4,000 Hz) or quitting prematurely. Block 3 (5 days, orange epoch) consisted of a 1 to 15 s (4,000–9,418 Hz) range. The fourth and final block (red epoch) consisted of offers ranging from 1 to 30 s (4,000–15,223 Hz) and continued for the remainder of the experiment. This block-wise protocol allowed us to examine how mice initially shaped their behavior as they learned the basic structure of the task as well as how they adjusted their foraging strategies across long timescales (days, weeks, and months) as they transitioned from reward-rich to reward-scarce environments in a stepwise manner. That is, blocks 1-3 are relatively reward rich as costs are generally low delays and rewards are easier to earn compared to block 4 which is relatively reward scarce as there are far more expensive reward offers distributed in the environment for which animals must learn to forage efficiently. The rewards used in this task only differ in flavor for which animals have subjective differences in preferences quantified below. All flavor options become increasingly scarce, equally across the longitudinal paradigm.

Rewards were delivered using a custom-built 3D-printed automated pellet dispenser that was triggered by a computer running the task programmed in ANY-Maze (Stoelting), run under dim lighting. The task was run by video-tracking animal position in real-time where all choice contingencies are determined by the centroid of the animal’s body crossing territories in the maze in combination with being required to run in a counterclockwise direction. The key zones in the maze include each restaurant’s separate OZ and WZ as well as the hallways in between restaurants (thus, 12 total main zones). 6 key behavioral epochs make up all possible, mutually exclusive behaviors animals could be engaged in while spending their limited time budget. 1) **Enter time**: defined from OZ entry in the correct heading direction (counterclockwise around the maze) triggering tone onset until the animal physically enters right into the WZ triggering tone countdown. 2) **Skip time**: defined from OZ entry in the correct heading direction triggering tone onset until the animal physically exits left into the hallway. 3) **Quit time**: defined from WZ entry and tone countdown onset until the animal physically exits the WZ prematurely before the countdown completes, rescinding the offer and terminating the countdown. 4) **Earn time**: defined from WZ entry and tone countdown onset until the countdown completes and a pellet is delivered. 5) **Consumption time**: defined from reward delivery onset until the animal physically leaves the reward site and exits the WZ. 6) **Travel time**: defined from previous trial termination (either from a skip, quit, or consumption event as defined above) and hallway entry until the next trial begins, triggered by OZ entry into the next restaurant.

Body weight was measured twice daily, immediately before and after testing on the Restaurant Row task.

### Data analysis

In addition to simple behavioral metrics like rewards earned, laps run, reaction times as described above, a key neuroeconomic behavioral metric is decision thresholds. This metric approximates indifference points in choices made either in the OZ or WZ as a function of cost and flavor. We measured the inflection point of fitting Heaviside step regressions to enter versus skip choices in the OZ or earn versus quit choices in the WZ as a function of cued offer cost in each restaurant. Thresholds determined this way were used for several subsequent analyses, including calculating reward value. The value of an offer can be calculated by subtracting a trial’s offer from one’s WZ threshold as this reflects the relative cost normalized to each animals’ willingness to wait and earn a reward (i.e., offer value <0 indicate offers typically not earned and thus should be skipped in the OZ or quit in the WZ as determined by each animal’s subjective flavor-specific decision policy which remained relatively stable across block 4 of testing). Similarly, when quitting in the WZ, the value left at the moment of quitting can be calculated by subtracting the time remaining in that trial’s countdown from the WZ threshold (i.e., value left <0 indicates an amount of time left that is not worth waiting for and should have been quit whereas value left >0 would have been more advantageous to finish waiting at that point instead of quitting). The amount of time spent remaining in the countdown at the moment of quitting relative to what the starting offer was and how much time was spent also served as the basis of our analysis of sensitivity to sunk costs, as previously published in mice, rats, and humas whereby time spent waiting can decrease the probability of quitting independent of time left in the countdown, see Sweis et al *Science* for the development of this analysis ^16,18,54,56,60^. Importantly, this analysis depends on a control analysis that iteratively re-calculates the probability of earning a reward as a function of time left remaining in the countdown from the 0 s sunk condition (see **Supplementary Fig. 8** for a graphical illustration of this control analysis). This allows all sunk cost metrics to be simplified into a delta score or change in likelihood of continuing to stay, or an escalation of commitment, due purely to the amount of time already spent waiting, independent of temporal distance to the goal, that controls for individual differences in one’s general willingness to wait or level of patience. Thresholds as well as economic choice history serve as the basis and criteria used in regret-related analysis whereby distinct violation vs. non-violation sequences are compared head-to-head in order to quantify the effects of mistake history on influencing choice behavior on the next trial. See Steiner and Redish 2014 *Nature Neuroscience* ^17^, Sweis et al 2018 *PLoS Biology* ^5^, and Durand-de Cuttoli 2022 *Science Advances* ^16^ for the development and full description of these analyses.

Thresholds were also used for an analysis we recently developed to capture how mice develop strategies that optimize food yield while also maximizing subjective value, see Durand-de Cuttoli et al 2023 *Biological Psychiatry* ^43^ for the development of this analysis. In brief, we calculated the Euclidean distance between the coordinates of observed thresholds and theoretical thresholds that would yield maximal food (empirically determined via simulations) versus maximal subjective value (calculated mouse-by-mouse based on revealed preferences determined on day 7 projected across the remainder of testing) on each day from days 18 to 45 during the 1-30 s epoch, calculated separately in both the offer zone and wait zone. We ran a computer Restaurant Row simulation in order to calculate the total number of pellets earned when varying different behavioral parameters. We determined that the ideal threshold required to obtain the theoretical maximum number of pellets when ignoring flavor preferences was empirically determined to be 10 s in all restaurants and all zones when tested in a 1-30 s offer range environment. In order to determine the thresholds that could yield the maximal amount of subjective value, we calculated the ratio of pellets earned on day 7 when all offers were 1s only on a mouse-by-mouse basis. When then multiplied the theoretical food earnings output from our simulation by this mouse-specific ratio in order to transform threshold by food output into a high dimensional subjective value space. From this we could extract what combination of thresholds in all 4 restaurants when tested in a 1-30 s environment could theoretically yield the greatest amount of subjective value for each mouse. From these two sets of theoretical threshold sets (for maximal food or subjective value) we could calculate the Euclidean distance from each animal’s actual observed thresholds on each day from either theoretical maximum.

Data were processed in Matlab with statistical analyses in JMP Pro 17. All data are expressed as mean ± 1 standard error. Statistical significance was assessed using student’s t tests, sign tests, and one-way, two-way, three-way, and repeated measures ANOVAs. No data were excluded as outliers.

## Acknowledgments

We thank Pete Rudebeck and members of the labs of Eric Nestler and Scott Russo for helpful discussion and technical assistance. Open-source illustrations obtained from SciDraw (www.scidraw.io), credit Federico Claudi and Annie Park.

## Funding

National Institute of Mental Health grant R01MH136230 (BMS)

National Institute of Mental Health grant R01MH051399 (EJN)

National Institute of Mental Health grant R01MH129306 (EJN)

National Institute of Mental Health grant R01MH114882 (SJR)

National Institute of Mental Health grant R01MH127820 (SJR)

National Institute of Mental Health grant R01MH104559 (SJR)

National Institute of Mental Health grant L40MH127601 (BMS)

National Institute of Mental Health training grant R25MH129256 (BMS)

National Institute of Mental Health supplement grant R01MH051399-31S1 (BMS)

Leon Levy Scholarship in Neuroscience, New York Academy of Sciences (BMS)

Burroughs Wellcome Fund Career Award for Medical Scientists (BMS)

Animal Models for the Social Dimensions of Health and Aging Research Network via NIH/NIA R24 AG065172 (BMS)

Brain & Behavior Research Foundation NARSAD Young Investigator Award 32856 (BMS)

Brain & Behavior Research Foundation NARSAD Young Investigator Award 31140 (RDC)

## Author contributions

Conceptualization: RDC, OI, EJN, BMS

Methodology: RDC, OI, BMS

Investigation: RDC, OI, SMBP, BY, NJ, AA, SK, KG, SA, SJR, EJN, BMS

Data curation: RDC, OI, BMS

Formal analysis: BMS

Visualization: RDC, OI, SMBP, BMS

Funding acquisition: OI, BMS

Supervision: RDC, BMS

Writing – original draft: RDC, OI, SMBP, BMS

Writing – review & editing: RDC, OI, SBMP, BY, NJ, AA, SK, KG, SA, SJR, EJN, BMS

## Data and materials availability

All data and materials used in the analysis are available in the manuscript, materials and methods section, or supplementary information.

## Supplementary Materials

### Supplementary Text

#### LINC00473 alters responses to limited task information during change-of-mind decisions

We characterized how access to task-relevant sensory information may be critically important for sex­ and LINC00473-dependent differences during change-of-mind decisions. This is important when considering what factors might be guiding on-going continuous re-evaluations and how much of this choice process depends on externally guided sensory information like that contained in the descending tone pitch cuing the countdown in the wait zone. After day 45, we tested mice for an additional 2 days during which tone playback on the task was silenced only in the wait zone but remained audible during offer zone presentations (**Supplementary Fig. 11a**). That is, mice had access to cost information signaled by tone pitch when making initial enter vs. skip decisions in the offer zone but received no countdown information once in the wait zone while making secondary re-evaluative decisions to quit or stay before earning rewards. This allows us to isolate properties of change-of-mind decision-making due to the internal mechanisms of re-evaluation in the absence of sensory information updated from the external world.

Overall, we found that males were more sensitive to this informational manipulation compared to females. Both GFP- and LINC00473-expressing males displayed a modest decreased number of laps run and total number of pellets earned, while female mice remained relatively unchanged between audible and silent conditions (laps [timepoint*sex]: F=10.678, p<0.01; laps [timepoint*virus]: *F=0.449, p=0.507;* earns [timepoint*sex]: F=10.124, p<0.01; earns [timepoint*virus]: *F=0.002, p=0.969,* **Supplementary Fig. 11b-c**). This task manipulation did not alter any overt changes in offer zone choices in any group as animals continued to discriminate the audible tones during cost-informed enter vs. skip decisions (offer: F=1186.818, p<0.0001; offer*timepoint: F=1.365, *p=0.243;* **Supplementary Fig. 11d**). Interestingly, when examining changes in wait zone thresholds, we found a significant sex difference when wait zone countdown information was withheld that was compounded by an additional LINC00473 effect in females only (**Supplementary Fig. 11e**). In males, compared to the audible condition, the lack of auditory information cuing mice on progress during the countdown in the silent wait zone condition drove an increase in wait zone thresholds for most preferred flavors (males: timepoint: F=22.976, p<0.0001; timepoint*virus: F=0.181, p=0.671 **Supplementary Fig. 11e**). Males also displayed a concomitant decrease in wait zone thresholds for least preferred flavors (males: timepoint: F=19.561, p<0.0001; timepoint*virus: F=0.526, *p=0.470,* **Supplementary Fig. 11e**). These data indicate that, without cued auditory guidance in the wait zone, males were more patient waiting for favorite flavors and less patient waiting for least favorite flavors (defined by a change in the indifference point of earn outcomes as a function of the trial’s starting offer). Conversely, GFP-expressing females only displayed a decrease in wait zone thresholds for less preferred flavors, with no change in thresholds for most preferred flavors (**Supplementary Fig. 11e**). LINC00473-expressing females however did not display any changes in wait zone thresholds in any restaurant compared between audible and silent wait zone conditions (females: timepoint [most preferred]: F=2.569, p=0.112; timepoint*virus [most preferred]: F=0.137, p=0.712; timepoint [least preferred]: F=12.185, p<0.0001; timepoint*virus [least preferred]: F=8.901, p<0.01).

These data indicate LINC00473-expressing females valued offers in a consistent manner with or without externally guided information, suggesting that their internal measures of time or how they hold and maintain information from the offer zone in working memory may be more stable. Strikingly, this effect of LINC00473 was specific to when information was manipulated only in the wait zone, and did not appear if information was manipulated in just the wait zone nor in both the offer zone and the wait zone (**Supplementary Fig. 12–13**).

When examining quit times in the wait zone, all mice displayed a modest increased latency to make quit decisions in the silent condition compared to the audible condition (timepoint: F=11.764, p<0.001; sex: F=2.358, p=0.126; virus: F=0.836, p=0.361, **Supplementary Fig. 11f**). Finally, when examining sensitivity of change-of-mind quitting behavior to the passage of time already spent waiting in the wait zone, we found that the silent condition unmasked a sex difference in GFP-treated mice whereby females show increased sensitivity to time expenditures compared to males - a sex difference that was previously only apparent during the audible condition in LINC00473-expressing groups (time spent: F=97.196, p<0.001; sex: F=36.989, p<0.0001; virus: F=1.899, p=0.168; **Supplementary Fig. 11g**). These data highlight how the integration between external cues and internally generated value representations is unique during change-of-mind decisions and is differentially processed in LINC00473-treated females. This implies that mPFC function is critical for different ways in which individuals may rehearse or update information during ongoing re­evaluations with vs. without explicit sensory cues, specifically during change-of-mind decisions.

### Supplementary Figures

**Supplementary Fig. 1.**
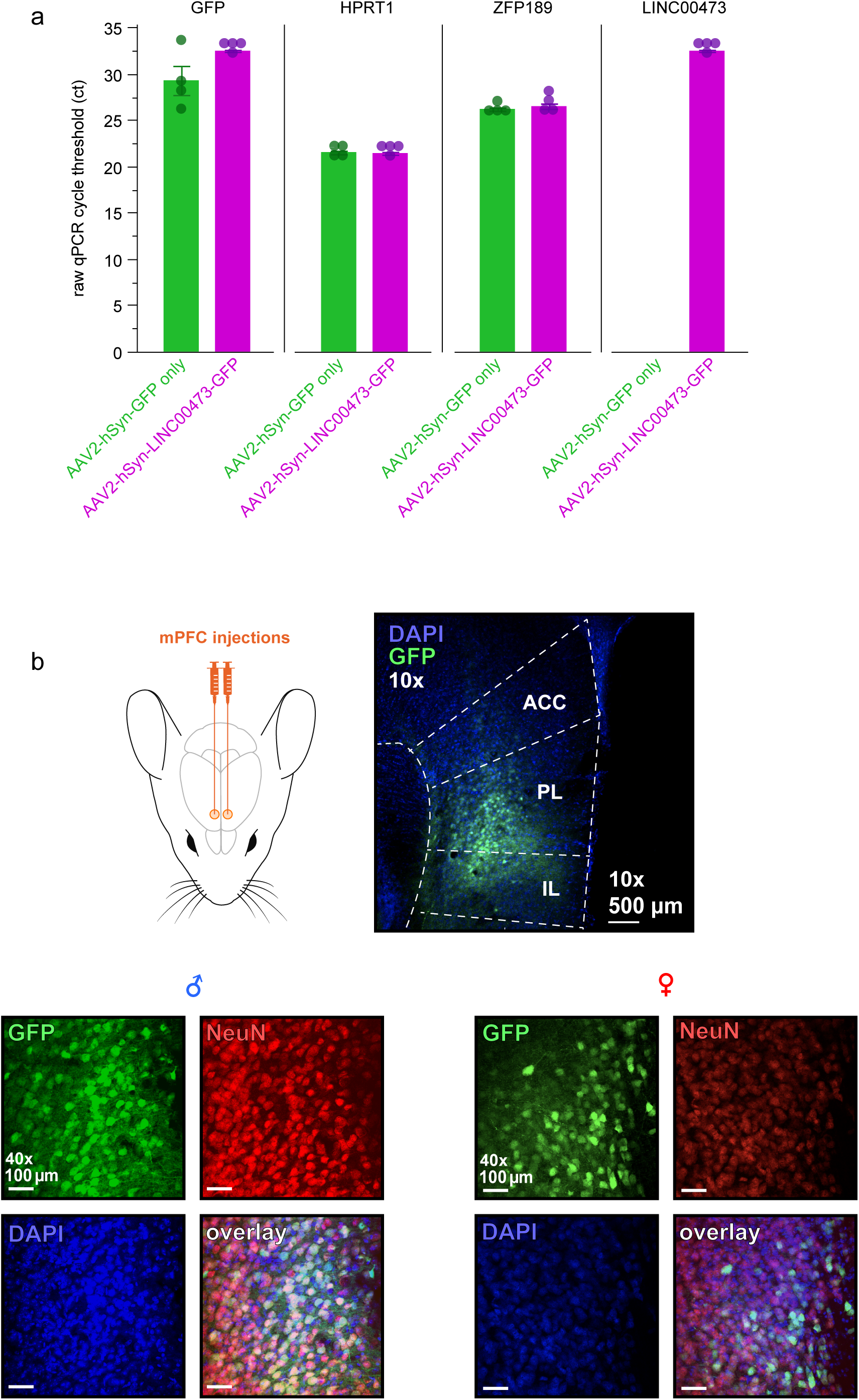
Validation of newly packaged adeno-associated virus expression of LINC00473. **a** Raw qPCR cycle threshold count (ct) obtain from mPFC tissue punches extracted from mice separately transfected with either AAV2-hSyn-GFP (control) or AAV2-hSyn-LINC00473-GFP (treatment). qPCR readouts obtained for expression levels of GFP, HPRT1 (housekeeping), ZFP189 (another known regulator of stress-resilience), and LINC00473. Note similar levels of expression of GFP (*t*=2.03, *p*=0.089), HPRT1 (*t*=0.41, *p*=0.693), and ZFP189 (*t*=0.91, *p*=0.396) between control and treatment viruses. Also note no detectable levels of LINC00473 in mice transfected with the control virus, as this is not endogenously expressed in mice. Dots represent individual samples. Error bars represent ±1 SEM. **b** Representative images of surgical targeting of mPFC bilaterally. 10x magnification image shows representative virus transfection of GFP toward ventral mPFC (generally including both prelimbic and infralimbic subregions). 40x images show representative targeting in both male and female mice depicting virus transfection of GFP, NeuN staining of neurons, and DAPI of cell bodies, both in separate channels and overlaid.

**Supplementary Fig. 2.**
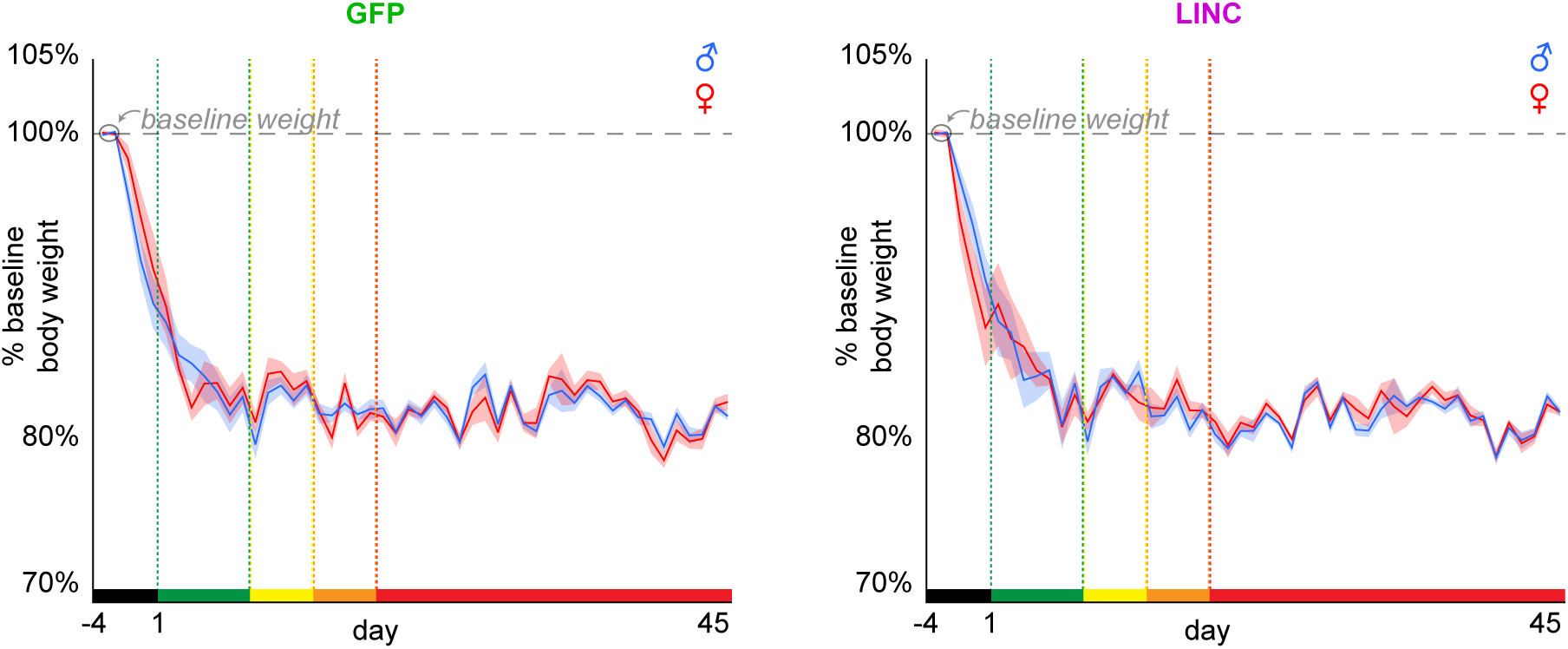
Bodyweight measurements. Percentage of a 2-day average (circled in gray, horizontal dashed line for reference) of baseline body weight obtained 4 days prior (black epoch) to the start of the longitudinal Restaurant Row paradigm. During this period (black epoch), mice were food restricted and fed a limited ration to intentionally decrease weights to approximately 80-85% of baseline that was steadily maintained throughout the remainder of the experiment (days 1-45, see main Fig. 1 for an explanation of the color-coded epochs of testing across the paradigm). These body weights reflect the measurements immediately before task performance on each day of testing. There were no differences in percentage baseline bodyweight between groups (virus: *F*=0.292, *p*=0.593; sex: *F*=0.118, *p*=0.733). Shaded area represents ±1 SEM.

**Supplementary Fig. 3.**
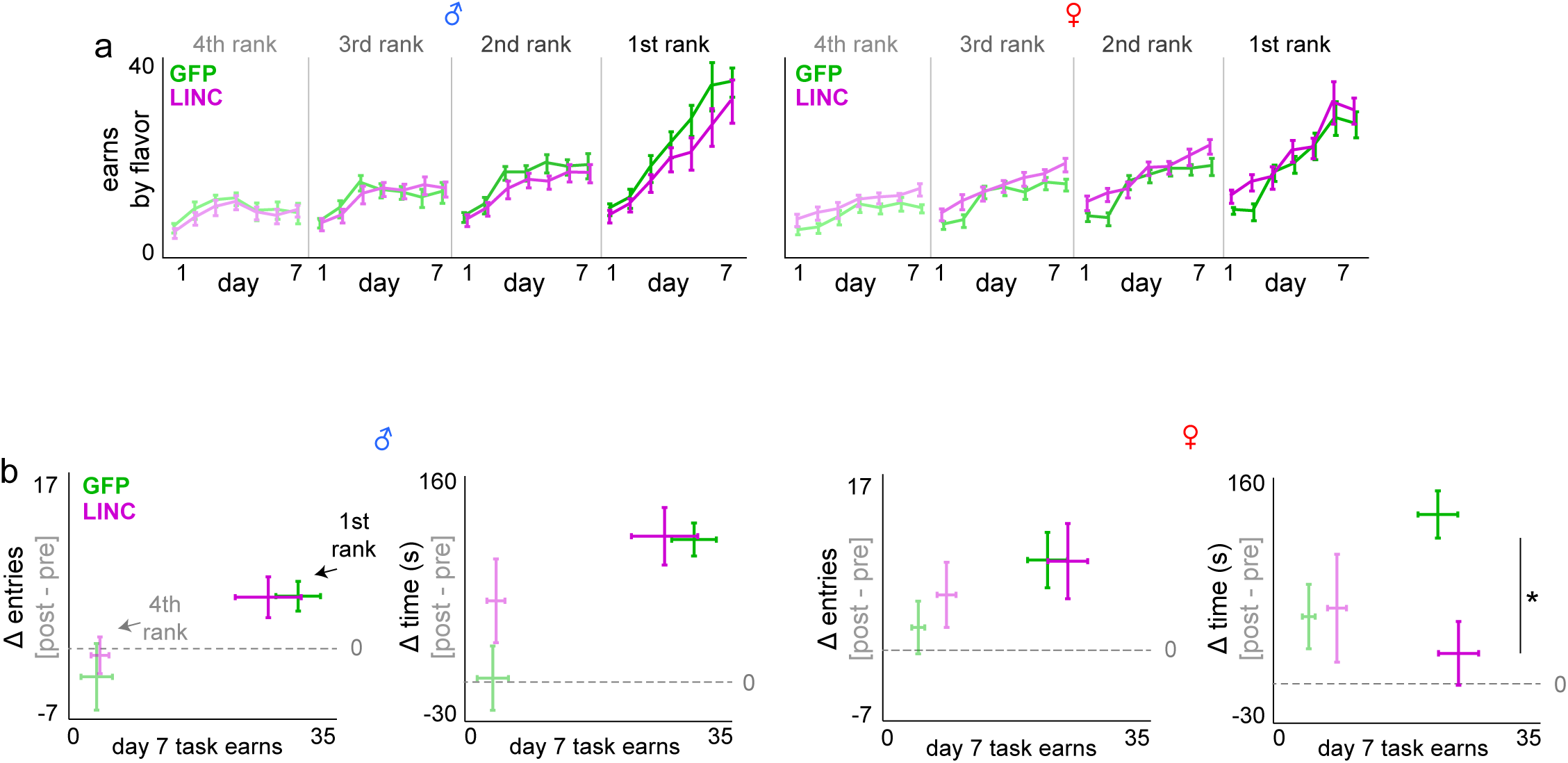
Redisplay of earns by flavor and CPP probe sessions. Figures redisplayed separating sex and superimposing GFP vs. LINC00473 groups. **a** Redisplay from main Fig. 1. Average number of rewards earned across the first week of testing split by flavors ranked from least to most preferred based on each day’s end-of-session totals. **b** Redisplay from main Fig. 2. Delta scores of total number of entries into and total time spent at each reward site subtracting post (day 8) minus pre (day 0) sessions. Two restaurants’ reward sites depicted: most and least preferred restaurants. Error bars represent ±1 SEM.

**Supplementary Fig. 4.**
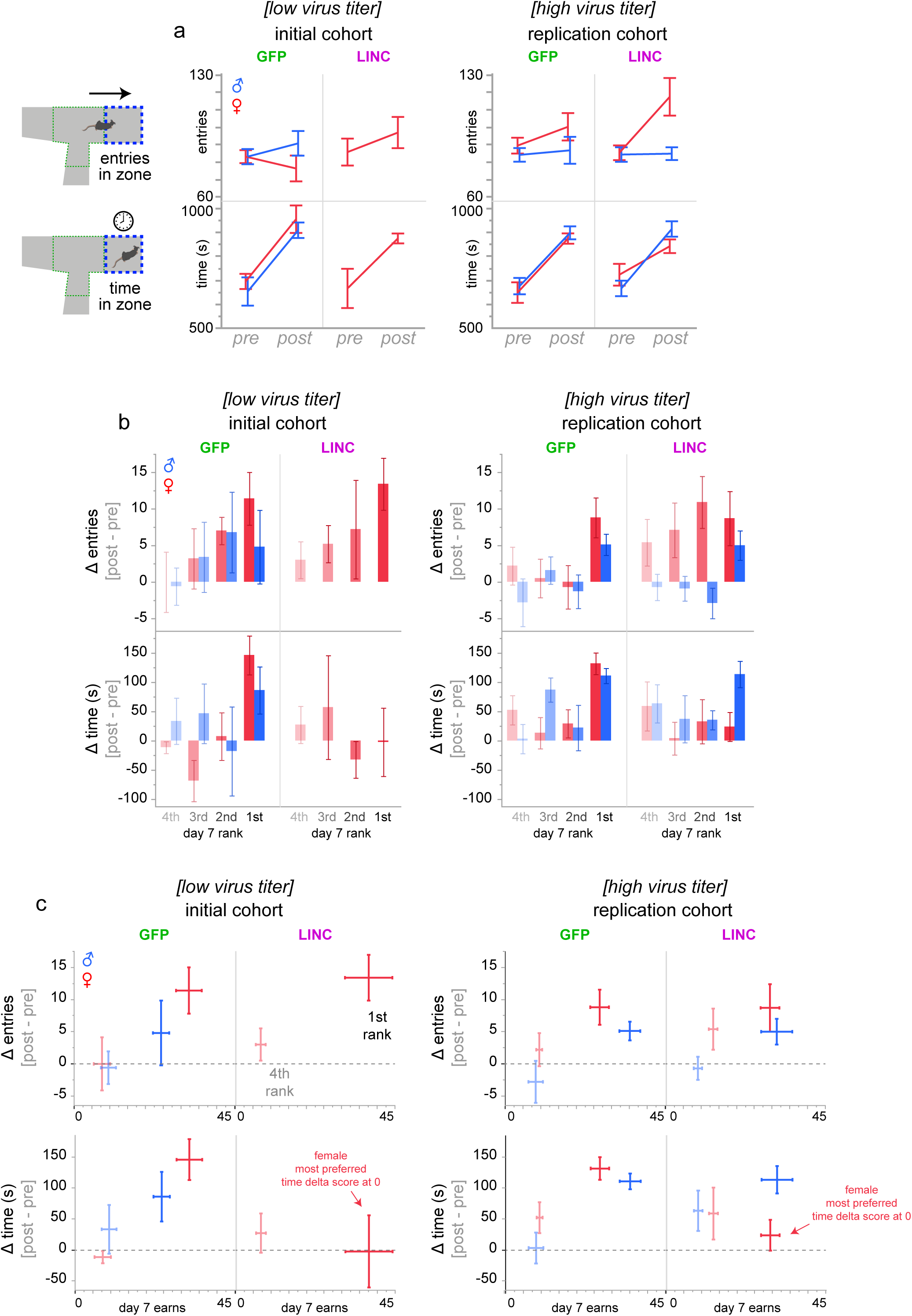
Replication cohort of two different virus titers on sex-specific Restaurant Row conditioned place preference (CPP) findings. Animals were placed in the arena free to roam for 20 min on day 0 and day 8 of the experimental timeline but with no active task to obtain pre-task baseline and post-task experience-related exploratory behavior (see main Fig. 2). **a** Total number of entries and total time spent in the wait zone, aggregated in all restaurants. Depicting findings from independent cohorts: surgeries and behavioral experiments were carried out across separate calendar months and using different virus titers. **b** Delta scores in change of entries and time comparing post (day 8) minus pre (day 0) time points split by restaurants ranked according to rewards earned on the active task on day 7. **c** Data from (b, only 1^st^ and 4^th^ ranking restaurants) replotted with number of rewards earned on day 7 on the x-axis. Horizontal dashed gray line indicates delta score of 0. LINC00473 expression in mPFC abolished CPP behavior in most preferred restaurants only in females and only in the time but not entry domain, replicated in both cohorts. +Sign-test on delta score of time: low virus cohort: male GFP: (*t*=2.142, *p*<0.05); female GFP: (*t*=4.424, *p*<0.01); female LNC: (*t*=0.04, *p*=0.516); high virus cohort: male GFP: (*t*=8.681, *p*<0.0001); male LINC: (*t*=5.087, *p*<0.001); female GFP: (*t*=7.157, *p*<0.0001); female LNC: (*t*=0.962, *p*=0.181). +Sign-test on delta score of entries: low virus cohort: male GFP: (*t*=0.955, *p*=0.197); female GFP: (*t*=3.155, *p*<0.05); female LNC: (*t*=3.766, *p*<0.01); high virus cohort: male GFP: (*t*=3.520, *p*<0.01); male LINC: (*t*=2.493, *p*<0.05); female GFP: (*t*=3.236, *p*<0.01); female LNC: (*t*=2.355, *p*<0.05). Error bars represent ±1 SEM.

**Supplementary Fig. 5.**
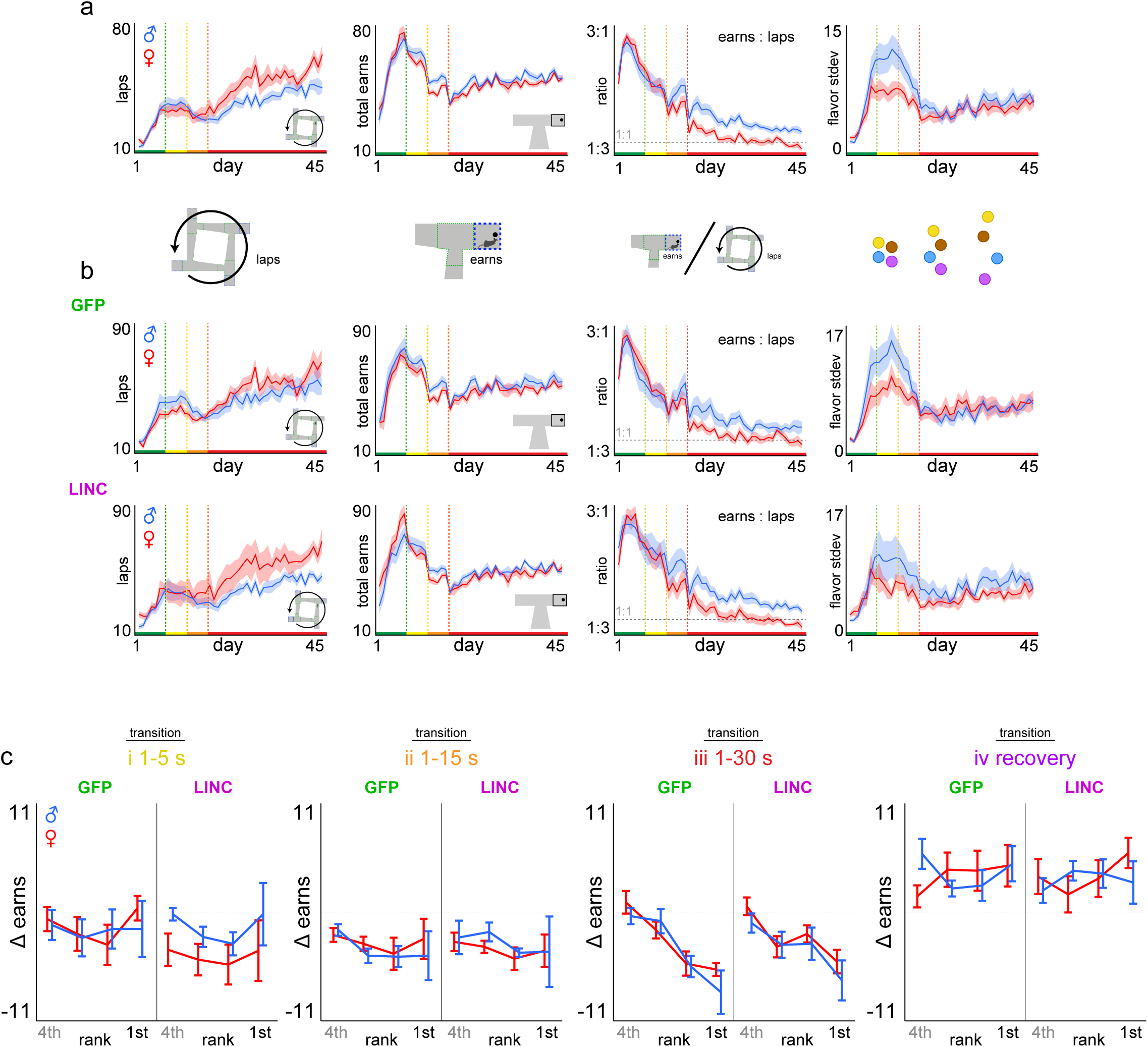
Basic Restaurant Row laps and earnings metrics across testing. **a** Overall sex difference comparisons between males and females (collapsing across GFP and LINC00473 virus treatment within each sex). Average number of laps run in the correct direction (sex*day: *F*=78.040, *p*<0.0001), total rewards earned (sex*day: *F*=1.331, *p*=0.249), ratio of earns / laps (sex*day: *F*=28.155, *p*<0.0001), and standard deviation of earns among flavors (sex*day: *F*=13.014, *p*<0.001). **b** Data from (a) split by GFP and LINC groups: laps (sex*virus*day: *F*=0.008, *p*=0.927), total earns (sex*virus*day: *F*=7.470, *p*<0.01), ratio of earns / laps (sex*virus*day: *F*=1.130, *p*=0.288), standard deviation of earns by flavor (sex*virus*day: *F*=15.486, *p*<0.0001). Statistical tests from (a) and (b) were performed on the entire experimental timeline (days 1-45). **c** Data from main Fig. 3d split by sex and virus groups depicting change in rewards earned in each restaurant comparing two days at each color-coded transition point. Delta scores calculated by subtracting the following: i, day 8-7 (yellow, rank: *F*=1.361, *p*=0.259; sex*virus: *F*=0.017, *p*=0.897); ii, day 13-12 (orange, rank: *F*=1.123, *p*=0.343; sex*virus: *F*=0.086, *p*=0.769); iii, day 18-17 (red, rank: *F*=18.116, *p*<0.0001; sex*virus: *F*=0.076, *p*=0.783); iv, day 45-18 (purple, rank: *F*=0.603, *p*=0.614; sex*virus: *F*=0.375, *p*=0.542). Shaded area and error bars represent ±1 SEM.

**Supplementary Fig. 6.**
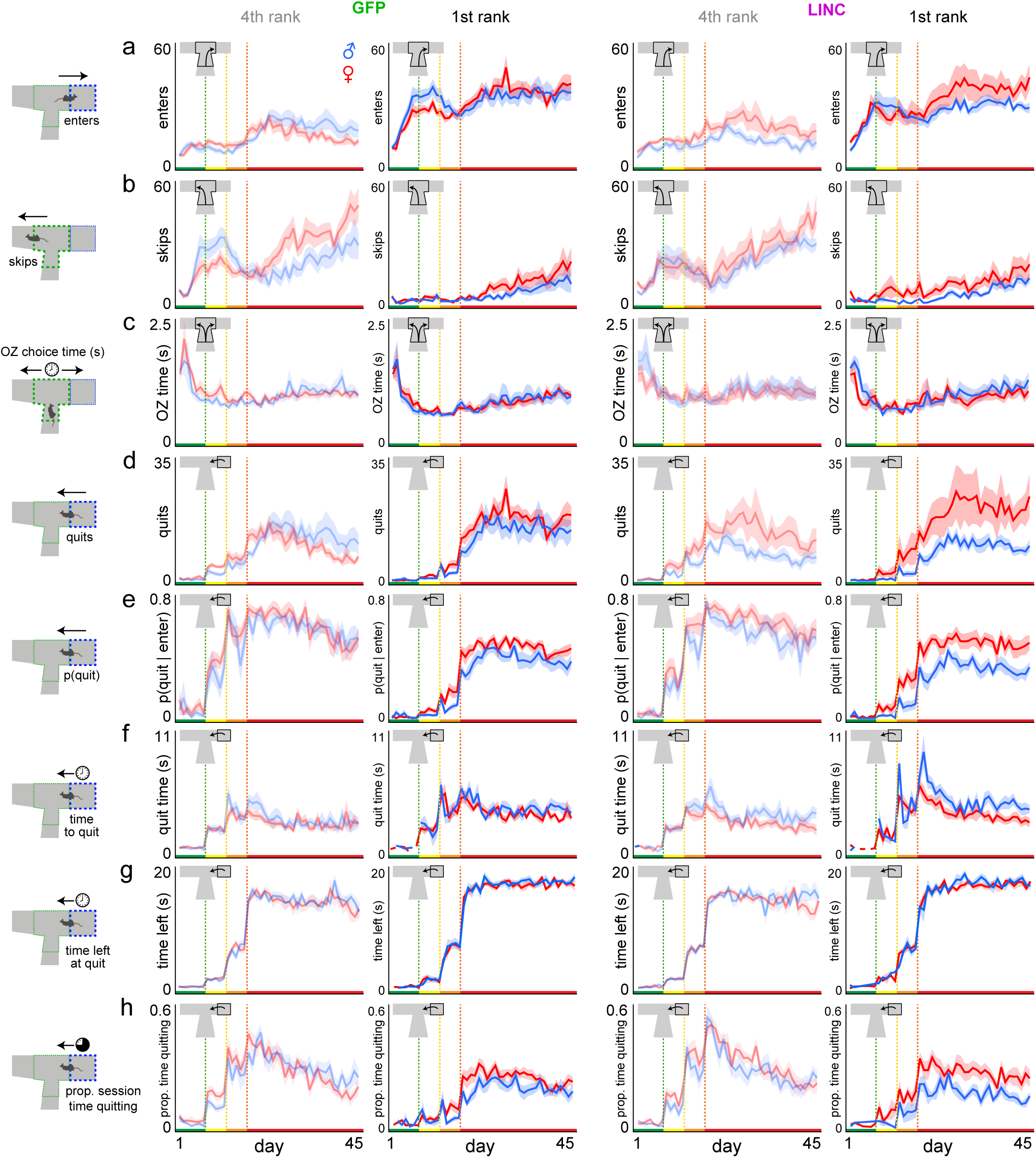
Restaurant Row choice metrics split by flavor ranking across testing. **a** Average number of enter decisions made in the offer zone split by sex and virus treatment groups as well as the 4^th^ and 1^st^ ranking restaurant (sex*rank*day: *F*=16.043, *p*<0.0001; sex*virus*rank*day: *F*=3.967, *p*<0.01). **b** Number of skip decisions made in the offer zone (sex*rank*day: *F*=9.894, *p*<0.0001; sex*virus*rank*day: *F*=2.617, *p*<0.05). **c** Offer zone reaction time before making an enter or skip decision calculated from offer zone entry (tone onset) to either a right turn into the wait zone (enter) or a left turn into the hallway (skip) (sex*rank*day: *F*=0.325, *p*=0.808; sex*virus*rank*day: *F*=2.186, *p*=0.087). **d** Number of quit decisions made in the wait zone during a countdown (sex*rank*day: *F*=11.725, *p*<0.0001; sex*virus*rank*day: *F*=1.216, *p*=0.302). **e** The probability of quitting in the wait zone given the animal entered (sex*rank*day: *F*=4.145, *p*<0.01; sex*virus*rank*day: *F*=0.463, *p*=0.708). This normalizes the likelihood of quitting to enter frequency. **f** Wait zone latency to quit reaction time from countdown onset (wait zone entry) until mice leave the wait zone prematurely (sex*rank*day: *F*=1.427, *p*=0.233; sex*virus*rank*day: *F*=3.160, *p*<0.05). **g** The amount of time left remaining in the countdown required to finish earning a reward at the moment of quitting in the wait zone. **h** Proportion of session time within each restaurant spent engaged in quitting behavior in the wait zone. Statistical tests here were performed on the entire experimental timeline (days 1-45). Shaded area represents ±1 SEM.

**Supplementary Fig. 7.**
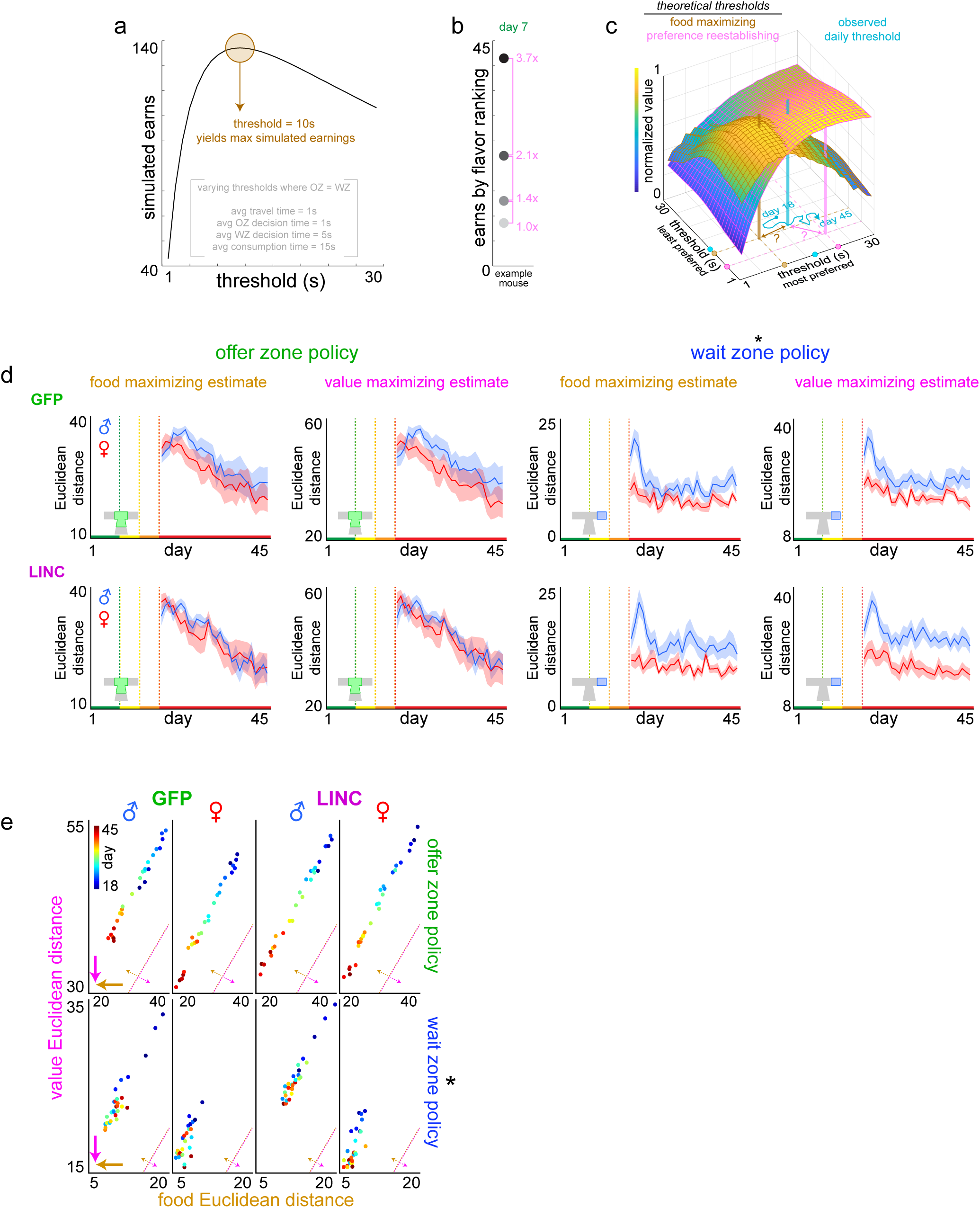
Longitudinal economic analysis of decision policy optimization for food security vs. subjective value. Analysis underlying data from main Fig. 4g-i derived from Durand-de Cuttoli et al 2023 *Biological Psychiatry* ^43^. See **Methods** for more details on this analysis. **a** Computer Restaurant Row simulation of total number of pellets earned. The ideal threshold required to obtain the theoretical maximum number of pellets when ignoring flavor preferences was empirically determined to be 10 s. **b** Relative ratio of earns among restaurants for an example mouse on day 7 between the flavor rankings (i.e., this mouse had a 3.7 : 2.1 : 1.4 : 1 ratio of earns across flavors capturing a summary of idealized relative subjective value for this mouse when all offers were 1 s only). **c** Two intersecting planes of decision policies that yield varying amounts of theoretical value either for maximal food intake (as determined by computer simulations, brown plane) or subjective value (as determined by multiplying simulation output of flavor earnings by day 7’s preference ratios on a mouse-by-mouse basis, pink plane) when in a reward-scarce environment (1 to 30 s offers). Here, only two decision policy dimensions (least preferred and most preferred restaurants) of the four dimensions (all four ranked restaurants) are displayed. Both planes are normalized to each’s min and max values for the purpose of plotting both in the same space while preserving the threshold coordinate locations that achieve either theoretical maximum (brown [location always fixed at threshold coordinates of 10 s] or pink beacons [location of discovered threshold coordinates that vary from mouse-to-mouse]). Actual observed daily decision policies represented by the cyan beacon wander throughout this space from day 18 to 45 in a reward-scarce environment. Trajectories projected to the floor of this display trace out individual mouse decision policy paths. Example coordinates of brown, pink, and cyan beacons illustrated as dots on the x and y axes. Euclidean distance from cyan coordinates to either brown or pink coordinates were calculated (question mark symbols). **d** Euclidean distances from observed decision policies in the offer zone (left) or wait zone (right) to either food (brown) or preference (magenta) theoretical maximum across testing in a reward-scarce environment split by GFP (top) and LINC (bottom) groups and sex. Offer zone: food distance: day: *F*=49.362, *p*<0.0001; sex: *F*=1.115, *p*=0.298; virus: *F*=0.008, *p*=0.929; value distance: day: *F*=42.042, *p*<0.0001, sex: *F*=1.546, *p*=0.221; virus: *F*=0.001, *p*=0.989; Wait zone: food distance: day: *F*=16.774, *p*<0.001, sex: *F*=13.796, *p*<0.001; virus: *F*=0.598, *p*=0.445; value distance: day: *F*=38.046, *p*<0.0001, sex: *F*=18.034, *p*<0.0001; virus: *F*=0.261, *p*=0.613. **e** Data from (d) redisplayed showing the Euclidean distance between observed decision policies each day from days 18-45 (cool to warm colors, each dot indicates group average per day) to theoretical policies that could achieve either maximal food (x-axis) or maximal subjective value (y-axis) for offer zone and wait zone thresholds. Inset diagonal lines indicate reference point for the 1:1 unity line. Bold brown and magenta arrows pointing toward the origin represent direction of reduced Euclidean distance vectors that optimize either food or value, respectively. Dashed brown and magenta arrows pointing off diagonal from the unity line represent if the decision policy is biased to prefer food vs. value. Prominent sex difference, without apparent effect of LINC00473 expression in mPFC, in present in wait zone but not offer zone economic decision policies. Shaded area represents ±1 SEM.

**Supplementary Fig. 8.**
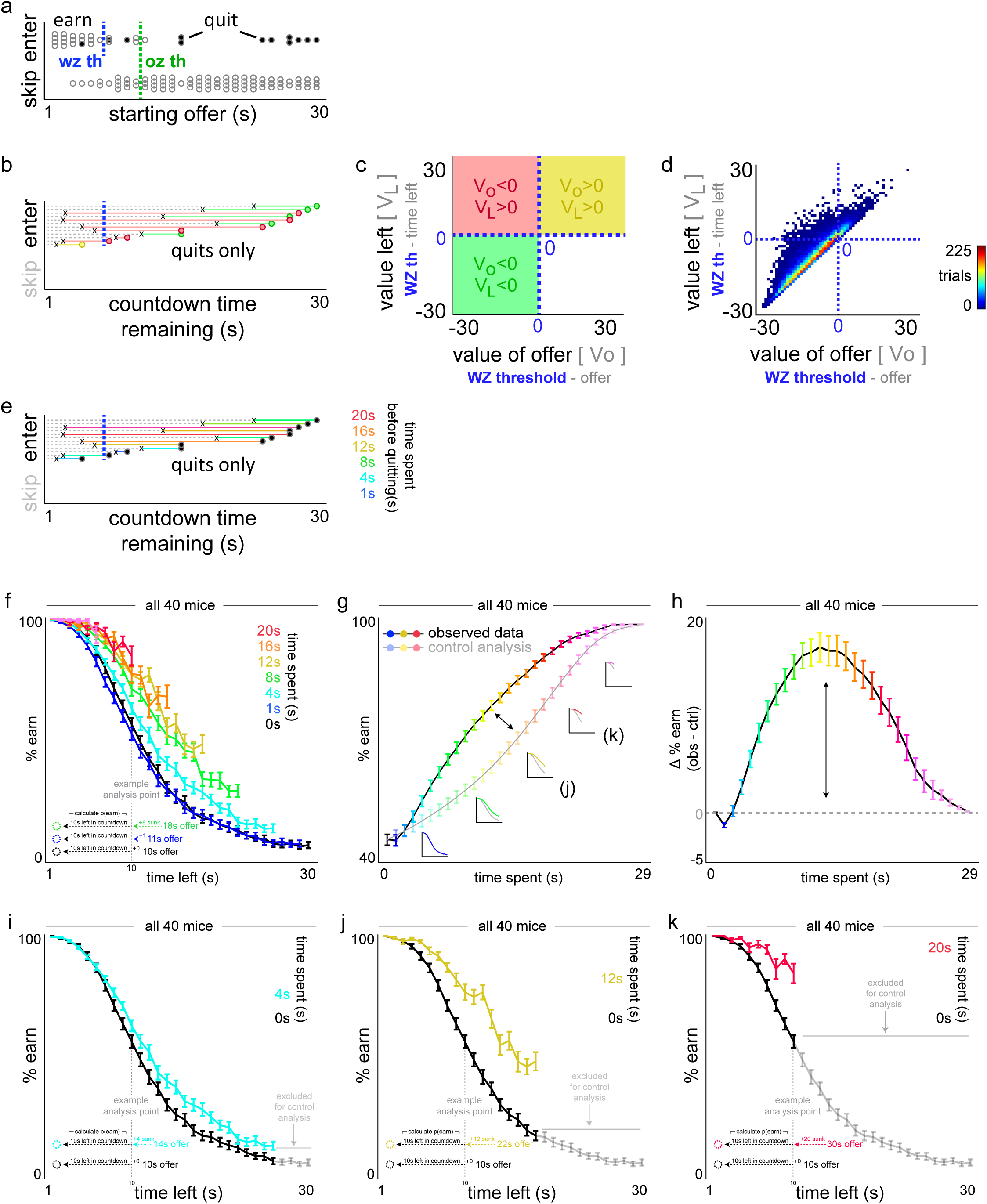
Visual explanation of characterizing quitting behavior. **a** Example choice data from a single mouse, single session, in a single restaurant. Individual dots represent individual trials. Offer zone outcomes plotted on the Y-axis as a function of the cued offer at the start of each trial along the X-axis. Of the enter trials, wait zone outcomes represent earns as open circles and quits as closed circles. Vertical dashed green and blue lines indicate the offer zone and wait zone thresholds, respectively. Note: the only temporal data shown here is the starting delay at the onset of the trial, and how each trial terminates as either a skip, enter then earn, or enter then quit. Only panels b-e include temporal data about how long it took animals to quit, for instance, and conversely, how much time was remaining in the countdown at the moment of quitting. **b** Data from (a) showing only the quit trials. Horizontal solid-colored lines extending from each dot leftward represent how much time was spent waiting before the mouse decided to quit. The point at which the mouse quit is represented by the black “x” symbol. The remaining dashed gray lines extending from the “x” leftward thus represent the amount of time remaining in the countdown at the moment of quitting. The relationship between one’s own wait zone threshold (vertical dashed blue line) and whether or not (i) the starting offer [circle] is to the right or the left of the wait zone threshold and (ii) the amount of time remaining at the moment of quitting [“x”] is to the right or the left of the wait zone threshold determines which quadrant each quit event belongs to in panels (c-d). Thus, each quit event is color coded based on its placement in the quadrants in (c-d). **c-d** Offer value plotted against value left can divide up this quit space into a two-by-two matrix based on the sign of each value term [i.e., if the starting offer was above or below one’s own wait zone threshold [represented by the dashed blue 0 lines] and if the time left at the moment of quitting was above or below one’s own wait zone threshold]. Data can only possibly exist in the three colored green, yellow, and red quadrants. Schematic in (c) and actual example data from main Fig. 5a re-displayed here. **e** Lastly, the time spent waiting before quitting is color-coded in yet another way, here based on how much time was already invested prior to the quit decision being made. Thus, this can separate rapid quit events from longer latencies to quit. These measures form one dimension of the data used in the sunk cost analysis, which also factors in varying amounts of time left remaining (horizontal gray dashed lines that represent the residual time at the moment of quitting). **f-k** Explanation of sunk cost and control analysis. (f-h) Redisplay of data from main Fig. 5f-h. Sunk cost analysis of staying behavior in the wait zone, demonstrated using all mice. (f) The likelihood of staying in the wait zone and earning a reward (e.g., not quit) is plotted as a function of time left in the countdown along the x-axis and time already spent waiting orthogonally in color. Note the black 0 s time spent curve represents animals having just entered the wait zone from the offer zone. Inset vertical dashed gray line illustrates an example analysis point comparing three sunk cost conditions originating from different starting offers but matched at 10 s left. Data from (f) dimensioned reduced in (g) collapsing across time left, instead highlighting the grand mean of each time spent sunk cost condition (color and x-axis). Insets depict data from curves in (f) are collapsed into the observed (sunk condition) and control (0 s condition) lines. Difference between curves in (g) are plotted in (h) in order to summarize the envelope of the overall effect of time already spent on escalating the commitment of staying in the wait zone. Horizontal dashed line represents a delta score of 0. (i-k) Redisplay of data from (f) but for three sunk cost conditions indicated in the insets in (g, 4 s cyan in i, 12 s gold in j, 20 s red in k) with the 0 sunk cost condition (black curve) repeated in each panel. Because each colored sunk cost curve derives from trials where the starting offer cost is higher than the matched time left value of the 0 s cost condition, some data points do not exist and are missing from the sunk cost curve on the rightward end. Thus, when reducing dimensions, to control for the inflated summary calculations of % earn due to missing data alone, the control analysis iteratively excludes matched data from the black 0 s sunk curve (highlighted in gray) before collapsing data to yield the resultant control curve in (g). Error bars represent ±1 SEM.

**Supplementary Fig. 9.**
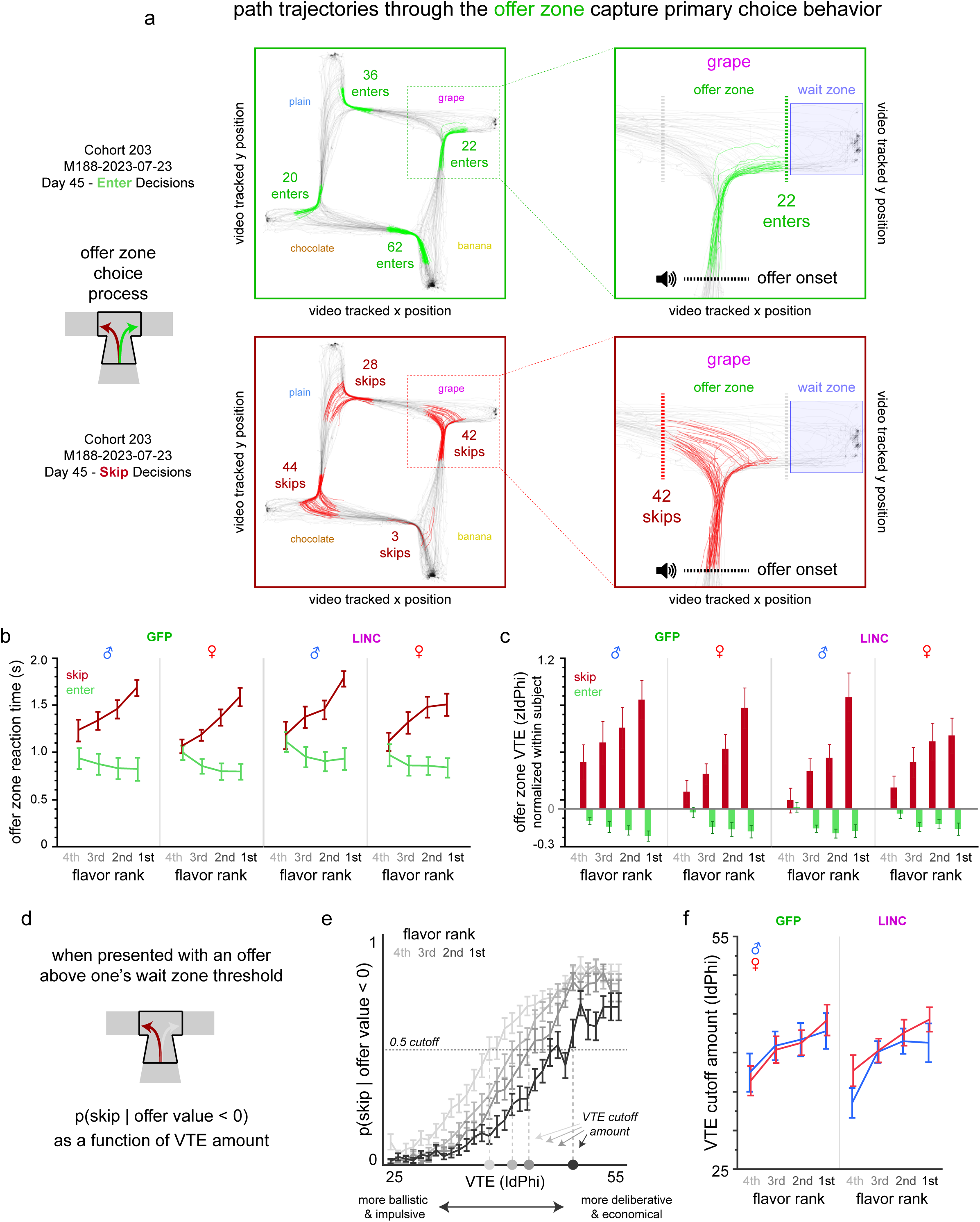
Video-tracking analysis of choice behavior in the offer zone. **a** Example video tracking data during a full 30-min session obtained from one example mouse (M188) taken from day 45 of testing. Track plots display X-Y body centroid position data of the mouse at 30 fps as the animal navigates the maze arena performing the task. All path trajectories through the offer zone on each trial beginning at trial onset (triggered by body centroid position crossing the stem of the entrance into the restaurant, which also triggers the auditory cue onset) until animals turn right and enter the wait zone (triggered by body centroid position crossing the wait zone boundary, which then triggers the countdown onset) are color-coded in green. All other position data are color-coded in gray. The number of enter decisions made in each restaurant are displayed. Zoomed-in example shows a closer depiction of the grape restaurant. The same data is redisplayed again but for skip decisions color-coded in red. Note the significantly greater heterogeneity in path trajectories through the offer zone, including a greater mix of heading-direction reorientation events before ultimately registering a skip outcome (triggered by body centroid position crossing the left boundary of the offer zone as animals enter the hallway connecting to the next restaurant) compared to enter decisions that are more uniformly ballistic and without path reorientations. **b** Offer zone reaction time measures plotted as a function of restaurant flavor preferences ranked from least to most preferred and split by whether or not mice made a skip or enter decision. Latencies to skip or enter were measured from trial and cued offer onset (traversing the stem entrance of each restaurant) until body centroid position crosses the rightward wait zone entry point or leftward hallway entry point. Significant interaction between choice outcome (enter vs. skip) and flavor ranking (*F*=14.201, *p*<0.0001) but no significant effects of sex (*F*=1.170, *p*=0.280) or virus (*F*=0.253, *p*=0.615). Reaction time alone does not necessarily capture individual differences in how ballistic vs. tortuous path trajectories can be through the choice point. Vicarious trial and error (VTE) behavior has been previously shown to capture heading-direction reorientation events and correlates with neural representations of deliberative decision-making processes (high VTE events) vs. the lack thereof (low VTE events). The physical “hemming and hawing” characteristic of VTE is best measured by calculating changes in velocity vectors of discrete body *X* and *Y* positions over time as *dx* and *dy*. From this, we can calculate the momentary change in angle, *Phi*, as *dPhi*. When this metric is integrated over the duration of the pass through the offer zone, VTE is measured as the absolute integrated angular velocity, or *IdPhi*, until either a skip or enter decision was made. This measure importantly is normalized within subject by z-scoring *IdPhi* values across all choices within a given session on a mouse-by-mouse basis, and then subsequently splitting the data by choice outcome and restaurant in order to appreciate within subject differences of how different types of choices and in different restaurants elicit varying amounts of either high or low VTE events. Significant interaction between choice outcome (enter vs. skip) and flavor ranking (*F*=19.819, *p*<0.0001) but no significant effects of sex (*F*=0.181, *p*=0.671) or virus (*F*=0.374, *p*=0.541). **d-f** Analysis of VTE behavior depicting the probability of skipping in the offer zone a negatively valued offer (offer above one’s wait zone threshold) as a function of binned VTE amounts. (e) Skipping probability given offer value <0 as a function of VTE amount split by restaurants ranked by flavor preferences for all 40 mice. Horizontal dashed line indicates a probability cutoff of 0.5 used to determine the X-intercept (vertical dashed lines shaded per restaurant), or amount of VTE displayed in order to reliably skip an economically disadvantageous offer. (f) VTE cutoff amount split by flavor rank, sex, and LINC00473 treatment. Significant effect of flavor ranking (*F*=9.372, *p*<0.0001) but no significant effects of sex (*F*=0.464, *p*=0.497) or virus (*F*=1.330, *p*=0.251). Data in (b-f) collapsed across the 1-30 s epoch (days 18-45). These data indicate that LINC00473 treatment nor sex has an impact on offer zone choice processes, including leaving intact deliberative behaviors that interact with the type of decision being made and counteract pre-potent impulsive responses to enter depending on the degree of flavor preference.

**Supplementary Fig. 10.**
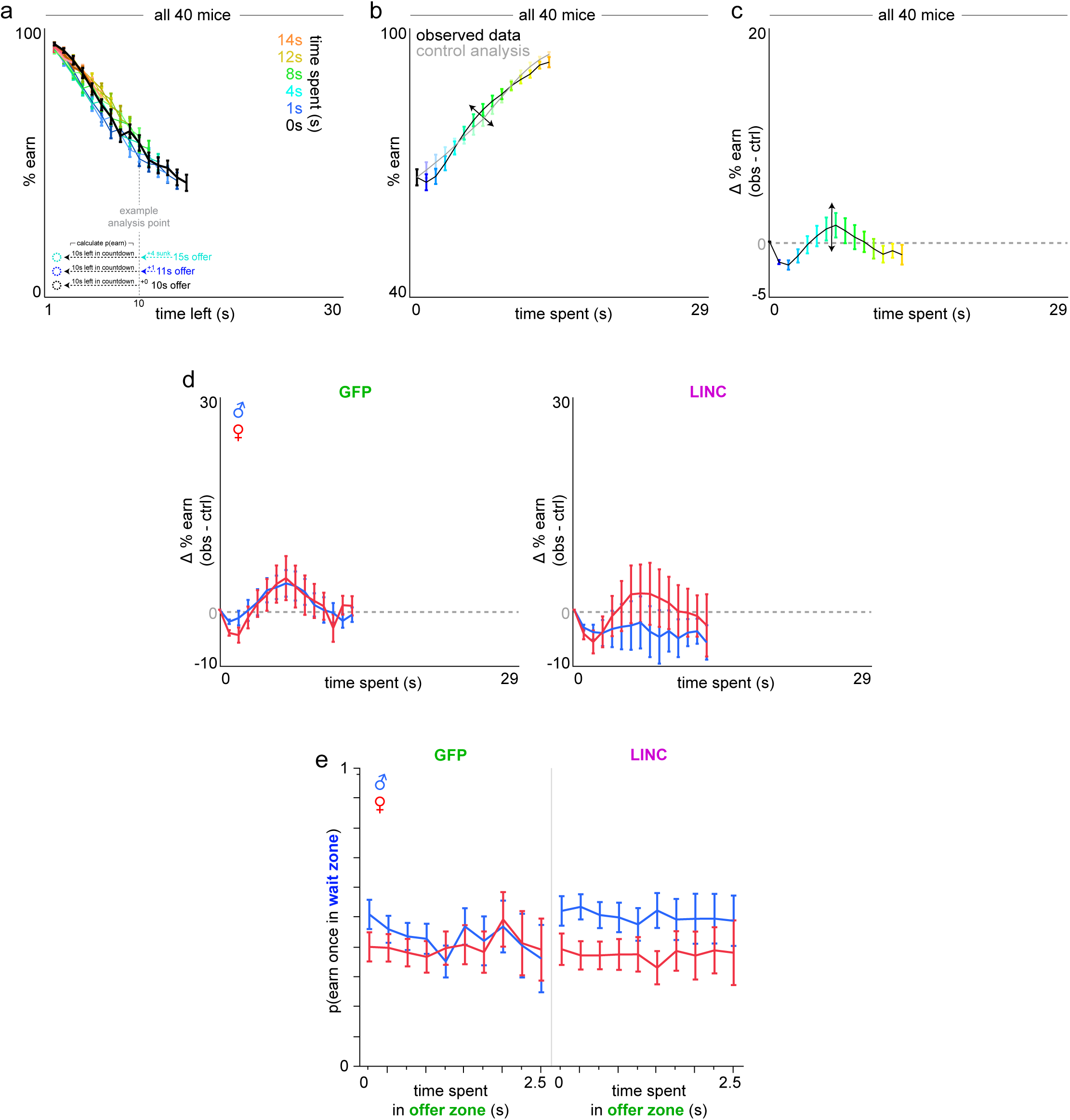
Sensitivity to sunk costs analysis during the 1-15 s testing epoch. Mice do not display sensitivity to sunk costs during days 13-17 in a relatively reward rich environment compared to days 18+ (when offers range 1-30 s, as in main Fig. 5). **a-c** Sunk cost analysis of staying behavior in the wait zone, demonstrated using all 40 mice. (f) The likelihood of staying in the wait zone and earning a reward (e.g., not quit) is plotted as a function of time left in the countdown along the x-axis and time already spent waiting orthogonally in color. Note the black 0 s time spent curve represents animals having just entered the wait zone from the offer zone. Inset vertical dashed gray line illustrates an example analysis point comparing three sunk cost conditions originating from different starting offers but matched at 10 s left. Data from (a) dimensioned reduced in (b) collapsing across time left, highlighting the grand mean of each time spent sunk cost condition (color and x-axis). Difference between curves in (b) are plotted in (c) in order to summarize the envelope of the overall effect of time already spent on escalating the commitment of staying in the wait zone. Horizontal dashed line represents delta score of 0. **d** Data in (c) split by sex and virus treatment groups depicting no effect of time spent during the countdown in the wait zone on changing the probability of quitting vs. staying to earn a reward (time spent: *F*=0.522, *p*=0.470; sex*virus*time spent: *F*=0.809, *p*=0.369). **e** To compare to time spent waiting in the wait zone, data here depict a different type of time spent on the task: time spent in the offer zone prior to accepting an offer and entering the wait zone does not influence the probability of quitting vs. staying to earn a reward once in the wait zone (time spent in offer zone: *F*=1.574, *p*=0.210; sex*virus*time spent: *F*=0.916, *p*=0.339). These data are from the 1-30 s reward-scarce epoch (days 18-45) to demonstrate the unique value of time spent waiting in the wait zone uniquely conferring sunk cost sensitivity. Error bars represent ±1 SEM.

**Supplementary Fig. 11.**
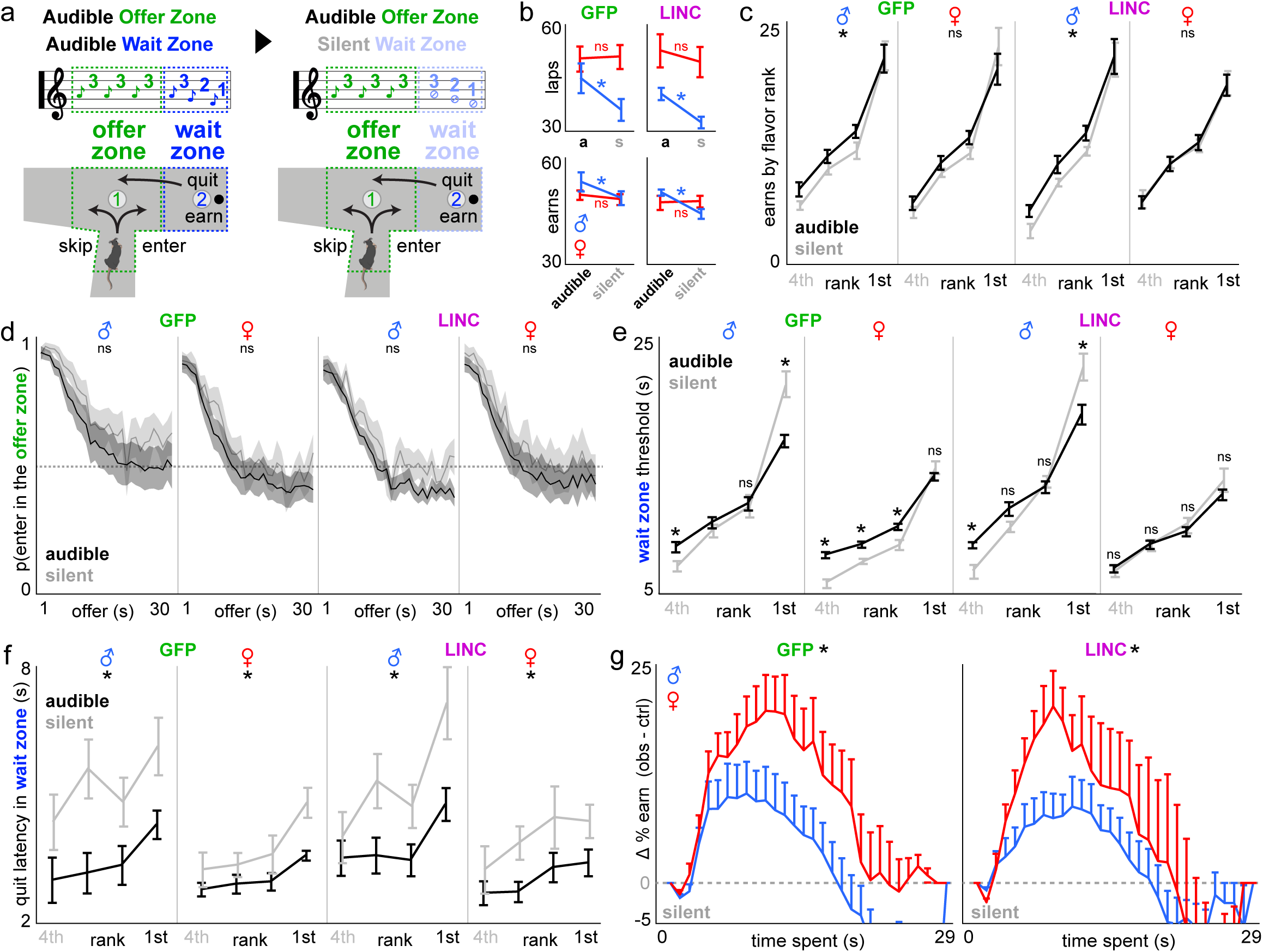
Sex- and LINC00473-dependent wait zone behaviors are differentially modulated by access to specific task information. **a** Informational manipulation schematic: (left) standard Restaurant Row task followed by (right) two days of tones silenced only in the wait zone, but not offer zone (remains audible as before). **b** Total number of laps run in the correct direction (top) and total pellets earned aggregated among restaurants (bottom) across the audible and silent wait zone task conditions (x-axis). **c** Rewards earned in each restaurant ranked from least (4^th^) to most preferred (1^st^) across the audible (black) and silent wait zone (gray) task conditions. **d** Overall offer zone choice probabilities to make enter decisions as a function of audible cued offer costs collapsed across all flavors to illustrate no effect of task condition on auditory discriminatory behaviors. Horizontal dashed line indicates change at 0.5. **e** Wait zone thresholds in each restaurant across task conditions. **f** Latency to quit reaction times once in the wait zone in each restaurant across task conditions. **g** Sensitivity to sunk costs in the wait zone depicted here only in the silent condition as a function of time already spent (x-axis). Y-axis reflects delta earn probabilities as depicted in main Fig. 5h. Prominent sex-dependent effects of withholding wait zone information on overall task performance (observed in males), wait zone economic decision policies (bidirectional in males, unidirectional in GFP-females, and no effect in LINC00473-females), and sunk cost sensitivity (sex difference revealed for the first time GFP groups unlike in main Fig. 5). Horizontal dashed line indicates delta score of 0. **\*** in (b-f) represent significant effects of task condition within group. * in (g) represent significant sex differences. Shading / error bars represent ±1 SEM. See **Supplementary Text**.

**Supplementary Fig. 12.**
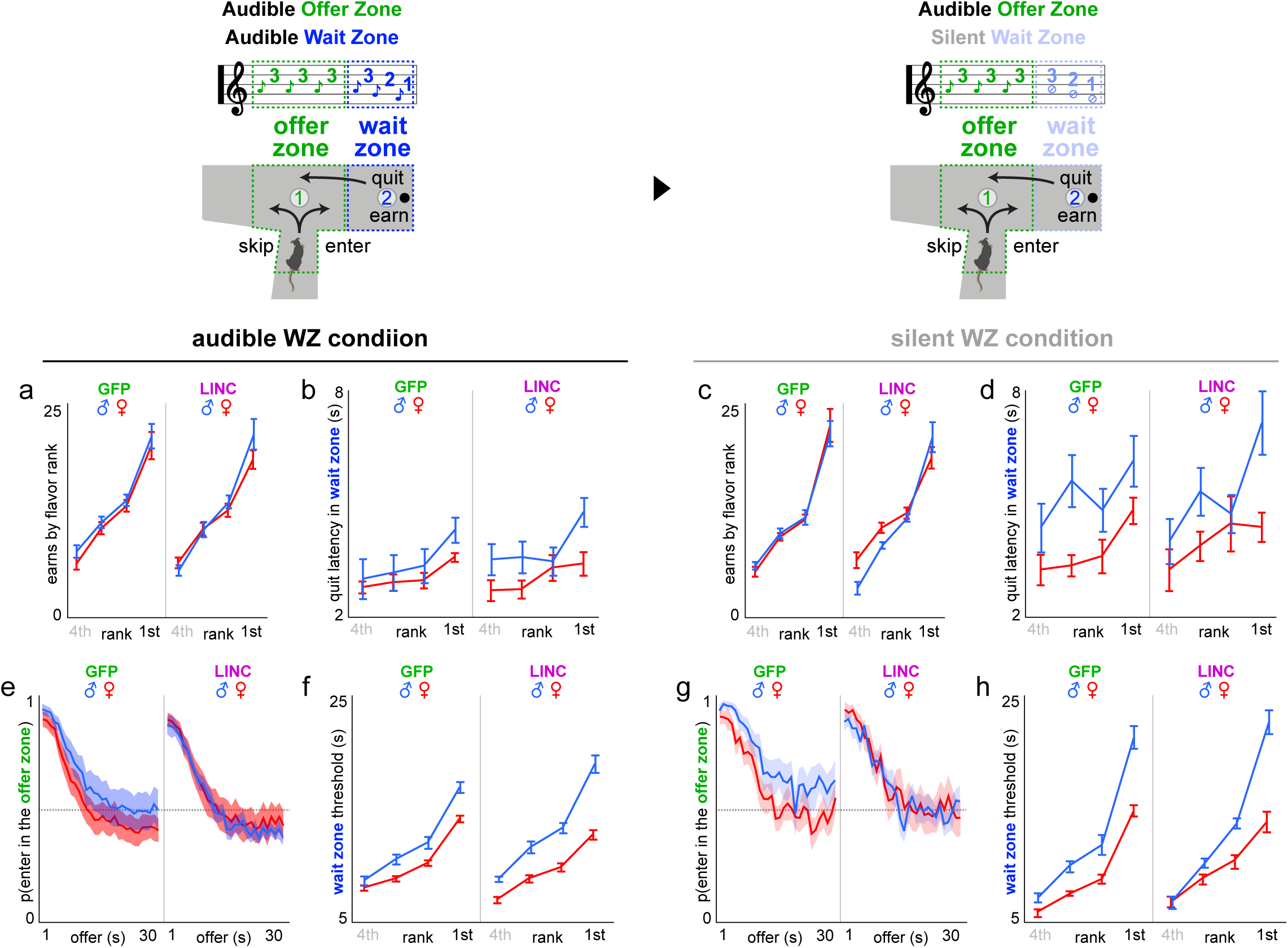
Redisplay of main metrics obtained from task information manipulation. Figures redisplayed separating task manipulation conditions (baseline audible wait zone (WZ) vs. silent WZ conditions) and superimposing sex. Redisplay from Supplementary Fig. 11. **a,c** Average number of rewards earned split by flavors ranked from least to most preferred based on end-of-session totals. **b,d** Average latency to quit after accepting an offer once in the wait zone. **e,g** Average proportion of trials accepted by making an enter decision from the offer zone into the wait zone as a function of cued offer cost in the offer zone. **f,h** Average wait zone threshold calculated from the inflection point of Heaviside-step function fits to wait zone outcome (earn vs. quit) as a function of the starting delay of each trial. Error bars and shading represent ±1 SEM.

**Supplementary Fig. 13.**
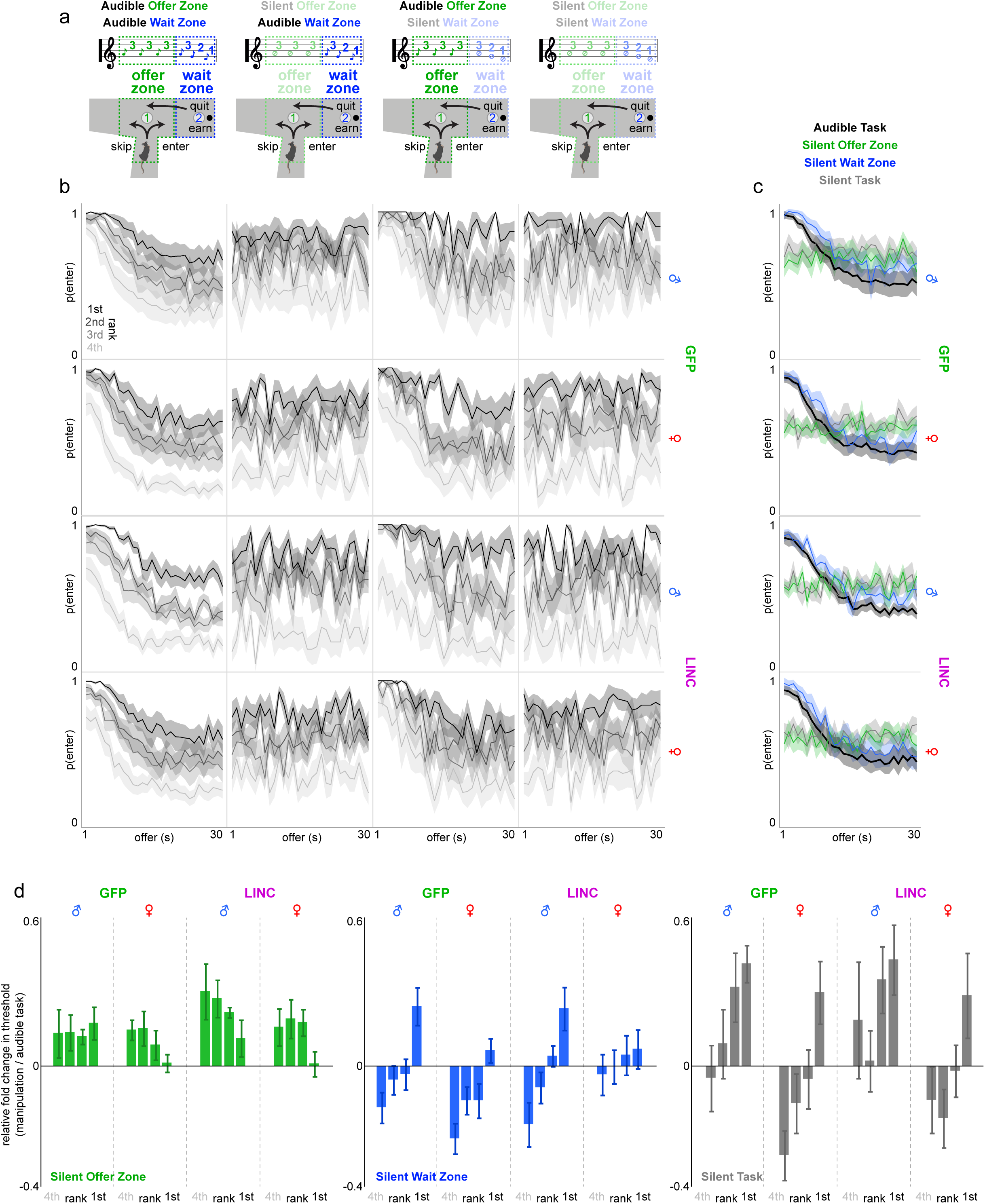
Full set of task information manipulations on Restaurant Row. **a** After day 45, mice were challenged with special probe days on three different task informational manipulations each for 2 days. Manipulations took the form of silencing and withholding tone presentation either in the offer zone, wait zone, or both: (i) silent offer zone but audible wait zone [days 46-47], (ii) audible offer zone but silent wait zone [days 48-49], and (iii) silent offer zone and silent wait zone [days 50-51]. **b** Offer zone choice probability to enter as a function of offer cost split by restaurants ranked from least (4^th^) to most (1^st^) preferred separating sex and virus treatment groups (rows) across the 4 different task information conditions (columns). **c** Data in (b) collapsed across all four restaurants showing summary enter probabilities as a function of offer cost. Note that enter probabilities are largely unaffected when the offer zone tones are audible comparing baseline (black, regular task) to silent wait zone (blue) conditions, however, as intended, offer zone choice probabilities change into a flat horizontal line in both the silent offer zone (green) and silent task (gray) conditions, as animals do not have any tone information in the offer zone during these conditions (audible task: offer: *F*=2017.115, *p*<0.0001; sex*virus*offer: *F*=1.912, *p*=0.167; silent offer zone task: offer: *F*=1.783, *p*=0.182; sex*virus*offer: *F*=0.227, *p*=0.634; silent wait zone task: offer: *F*=1004.552, *p*<0.0001; sex*virus*offer: *F*=0.147, *p*=0.702; fully silent task: offer: *F*=0.723, *p*=0.395; sex*virus*offer: *F*=1.723, *p*=0.190). **d** Wait zone thresholds plotted as a relative fold change comparing each condition to the regular task audible baseline separated across the three information manipulations (silent offer zone, left, green; silent wait zone, middle, blue; silent task, right, gray) split by restaurants ranked from least (4^th^) to most (1^st^) preferred, sex, and virus treatment groups. Note the difference in patterns in changes in wait zone thresholds across restaurants between sex/virus groups and across task information manipulation conditions. Sign tests: silent offer zone: male GFP: least preferred: *t*=+1.365, *p*=0.103; most preferred: *t*=+2.639, *p*<0.05; male LNC: least preferred: *t*=+2.699, *p*<0.05; most preferred: *t*=+1.538, *p*=0.079; female GFP: least preferred: *t*=+3.638, *p*<0.01; most preferred: *t*=+0.360, *p*=0.363; female LNC: least preferred: *t*=+2.125, *p*=<0.05; most preferred: *t*=+0.188, *p*=0.427; silent wait zone: male GFP: least preferred: *t*=−2.684, *p*<0.05; most preferred: *t*=+3.191, *p*<0.01; male LNC: least preferred: *t*=−2.624, *p*<0.05; most preferred: *t*=+2.685, *p*<0.05; female GFP: least preferred: *t*=−4.733, *p*<0.001; most preferred: *t*=+1.325, *p*=0.109; female LNC: least preferred: *t*=−0.411, *p*=0.345; most preferred: *t*=+0.886, *p*=0.199; silent task: male GFP: least preferred: *t*=−0.358, *p*=0.364; most preferred: *t*=+5.554, *p*<0.001; male LNC: least preferred: *t*=+0.794, *p*=0.225; most preferred: *t*=+3.049, *p*<0.001; female GFP: least preferred: *t*=−3.584, *p*<0.01; most preferred: *t*=+2.345, *p*<0.05; female LNC: least preferred: *t*=−1.009, *p*=0.170; most preferred: *t*=+1.671, *p*=0.064. Shaded area and error bars represent ±1 SEM. See **Supplementary Text**.

**Supplementary Fig. 14.**
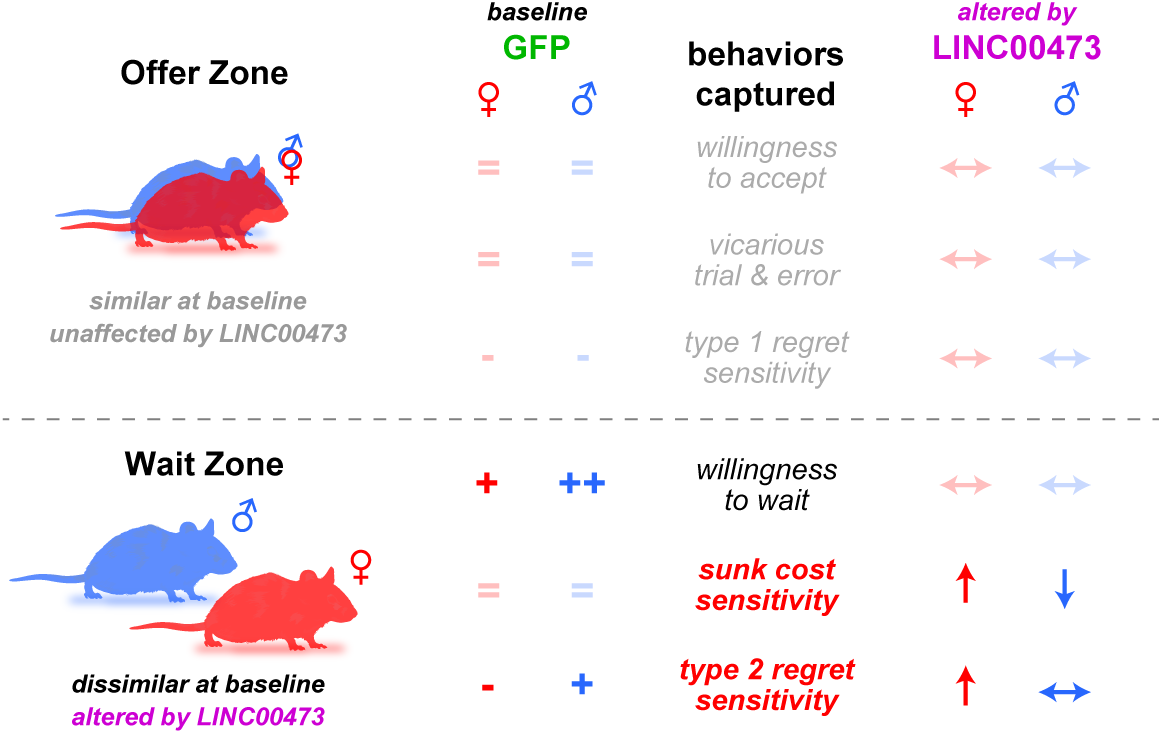
Expansion of summary of key findings. Expanded table from main Fig. 6 describing in more detail the direction of baseline effects (GFP group) with equal signs (=) if no sex differences, plus signs (+ or ++) to represent a sex difference in magnitude, or minus signs (−) to indicate baseline insensitivity to a given behavioral metric. Metrics altered by mPFC LINC00473 expression are represented by arrow symbols showing either an increase (↑), decrease (↓), or no change (↔) relative to baseline. Bolded / opaque symbols highlight several key wait zone decision-making behaviors that are different between sexes at baseline or altered by LINC00473 expression, none of which appear in the offer zone.

