## Supplementary Information for "Change-of-mind neuroeconomic decision-making is modulated by LINC00473 in medial prefrontal cortex in a sex-dependent manner"

#### Supplementary Figures 1-14

Supplementary Figure 1

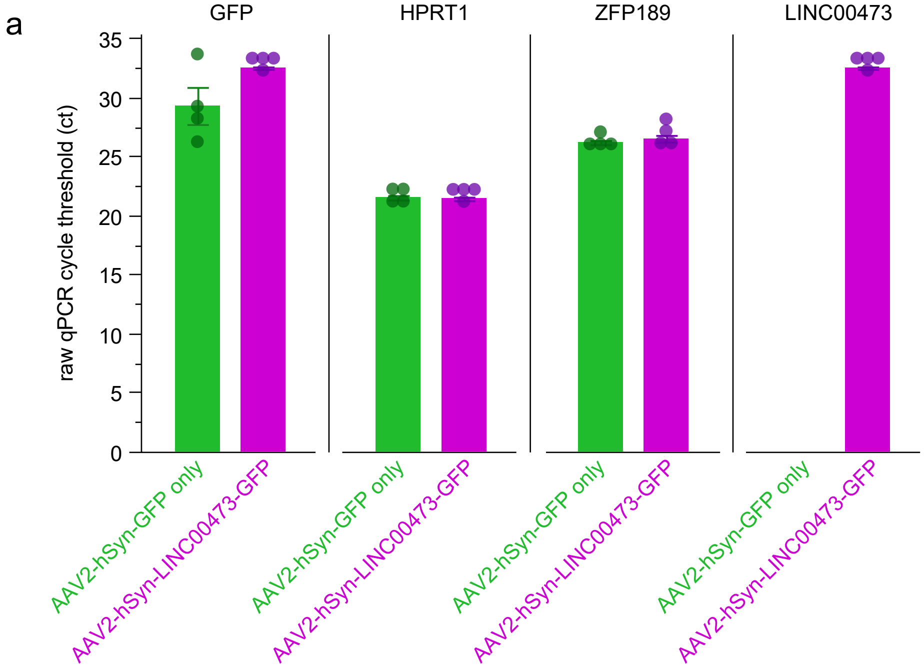


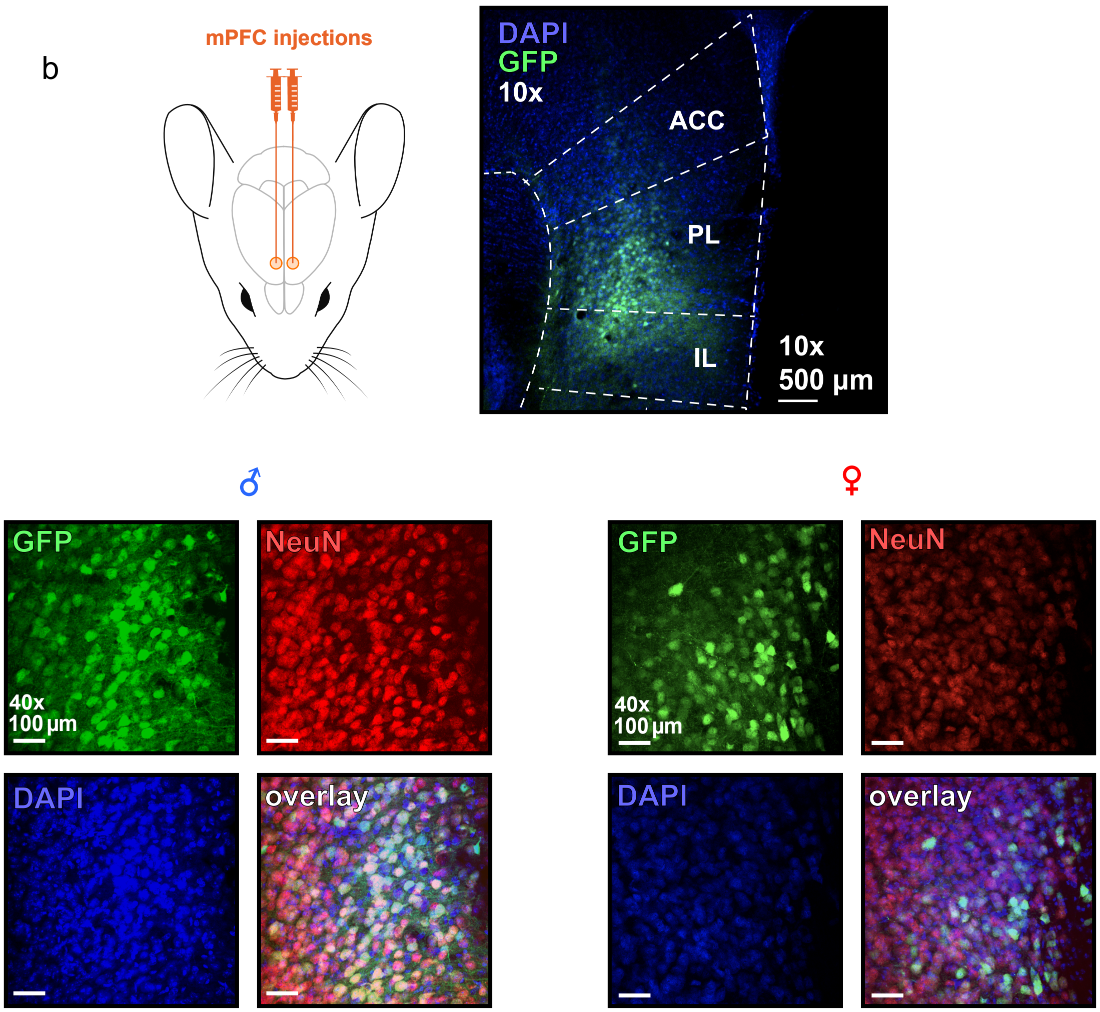

**Supplementary Fig. 1 | Validation of newly packaged adeno-associated virus expression of LINC00473. a** Raw qPCR cycle threshold count (ct) obtain from mPFC tissue punches extracted from mice separately transfected with either AAV2-hSyn-GFP (control) or AAV2-hSyn-LINC00473-GFP (treatment). qPCR readouts obtained for expression levels of GFP, HPRT1 (housekeeping), ZFP189 (another known regulator of stress-resilience), and LINC00473. Note similar levels of expression of GFP ($t$=2.03, $p$=0.089), HPRT1 ($t$=0.41, $p$=0.693), and ZFP189 ($t$=0.91, $p$=0.396) between control and treatment viruses. Also note no detectable levels of LINC00473 in mice transfected with the control virus, as this is not endogenously expressed in mice. Dots represent individual samples. Error bars represent ±1 SEM. **b** Representative images of surgical targeting of mPFC bilaterally. 10x magnification image shows representative virus transfection of GFP toward ventral mPFC (generally including both prelimbic and infralimbic subregions). 40x images show representative targeting in both male and female mice depicting virus transfection of GFP, NeuN staining of neurons, and DAPI of cell bodies, both in separate channels and overlaid.

Supplementary Figure 2

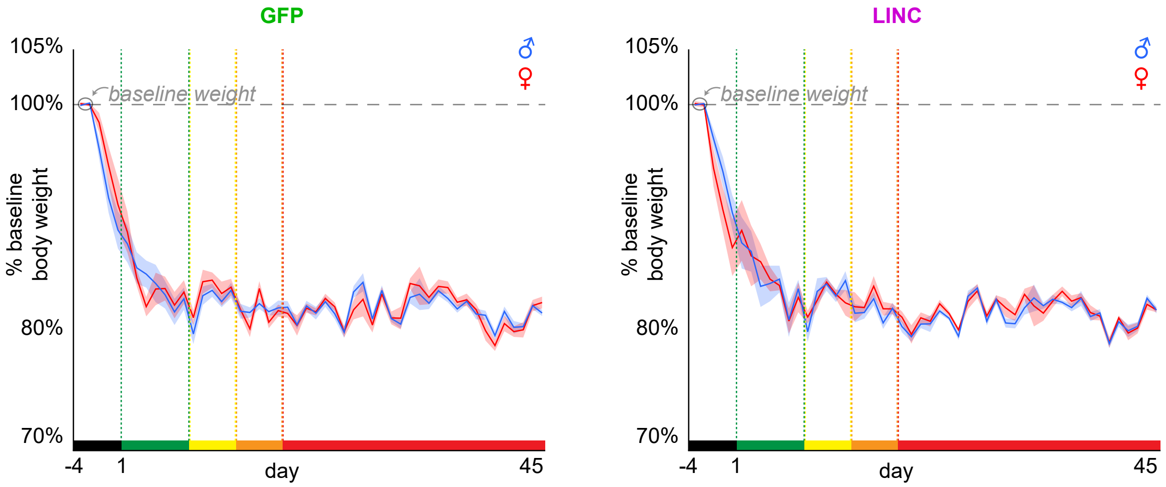

**Supplementary Fig. 2 | Bodyweight measurements.** Percentage of a 2-day average (circled in gray, horizontal dashed line for reference) of baseline body weight obtained 4 days prior (black epoch) to the start of the longitudinal Restaurant Row paradigm. During this period (black epoch), mice were food restricted and fed a limited ration to intentionally decrease weights to approximately 80-85% of baseline that was steadily maintained throughout the remainder of the experiment (days 1-45, see main Fig. 1 for an explanation of the color-coded epochs of testing across the paradigm). These body weights reflect the measurements immediately before task performance on each day of testing. There were no differences in percentage baseline bodyweight between groups (virus: $F$=0.292, $p$=0.593; sex: $F$=0.118, $p$=0.733). Shaded area represents ±1 SEM.

Supplementary Figure 3

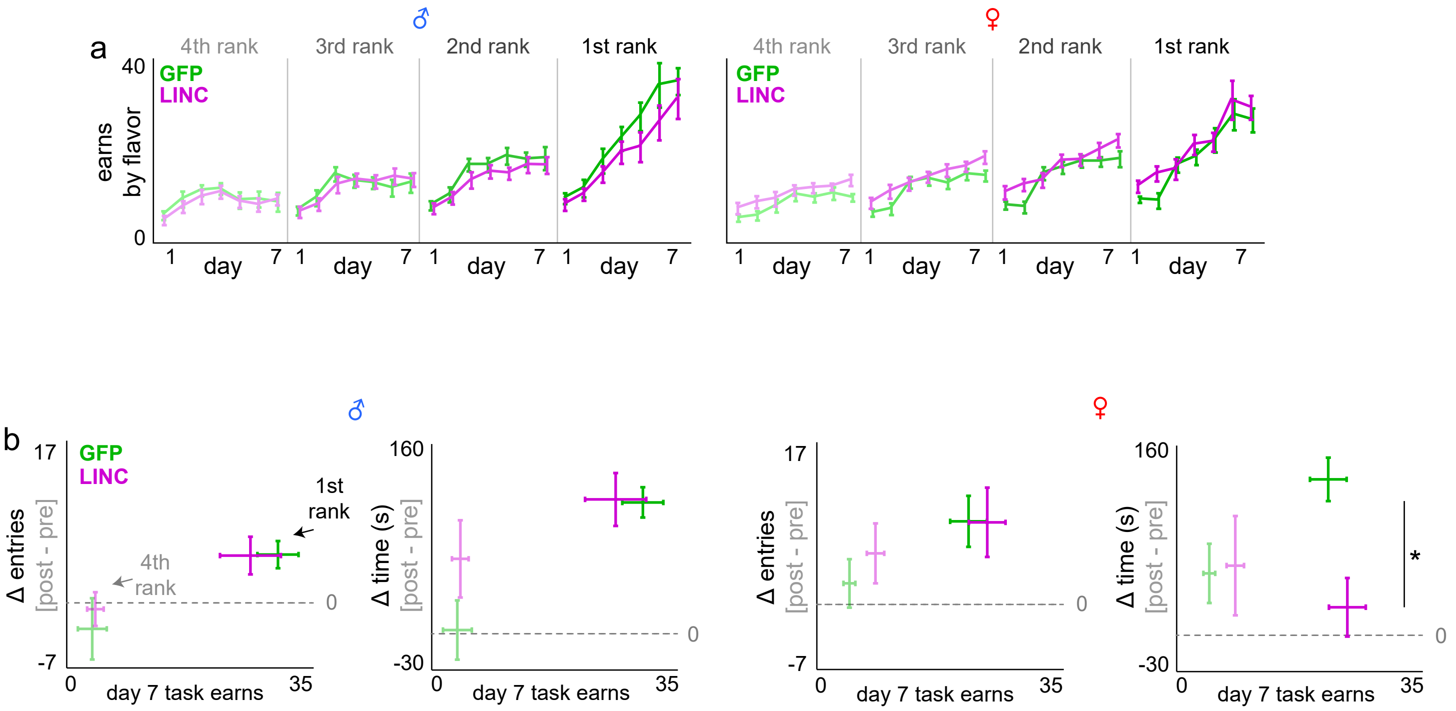

**Supplementary Fig. 3 | Redisplay of earns by flavor and CPP probe sessions.** Figures redisplayed separating sex and superimposing GFP vs. LINC00473 groups. **a** Redisplay from main Fig. 1. Average number of rewards earned across the first week of testing split by flavors ranked from least to most preferred based on each day's end-of-session totals. **b** Redisplay from main Fig. 2. Delta scores of total number of entries into and total time spent at each reward site subtracting post (day 8) minus pre (day 0) sessions. Two restaurants' reward sites depicted: most and least preferred restaurants. Error bars represent ±1 SEM.

Supplementary Figure 4

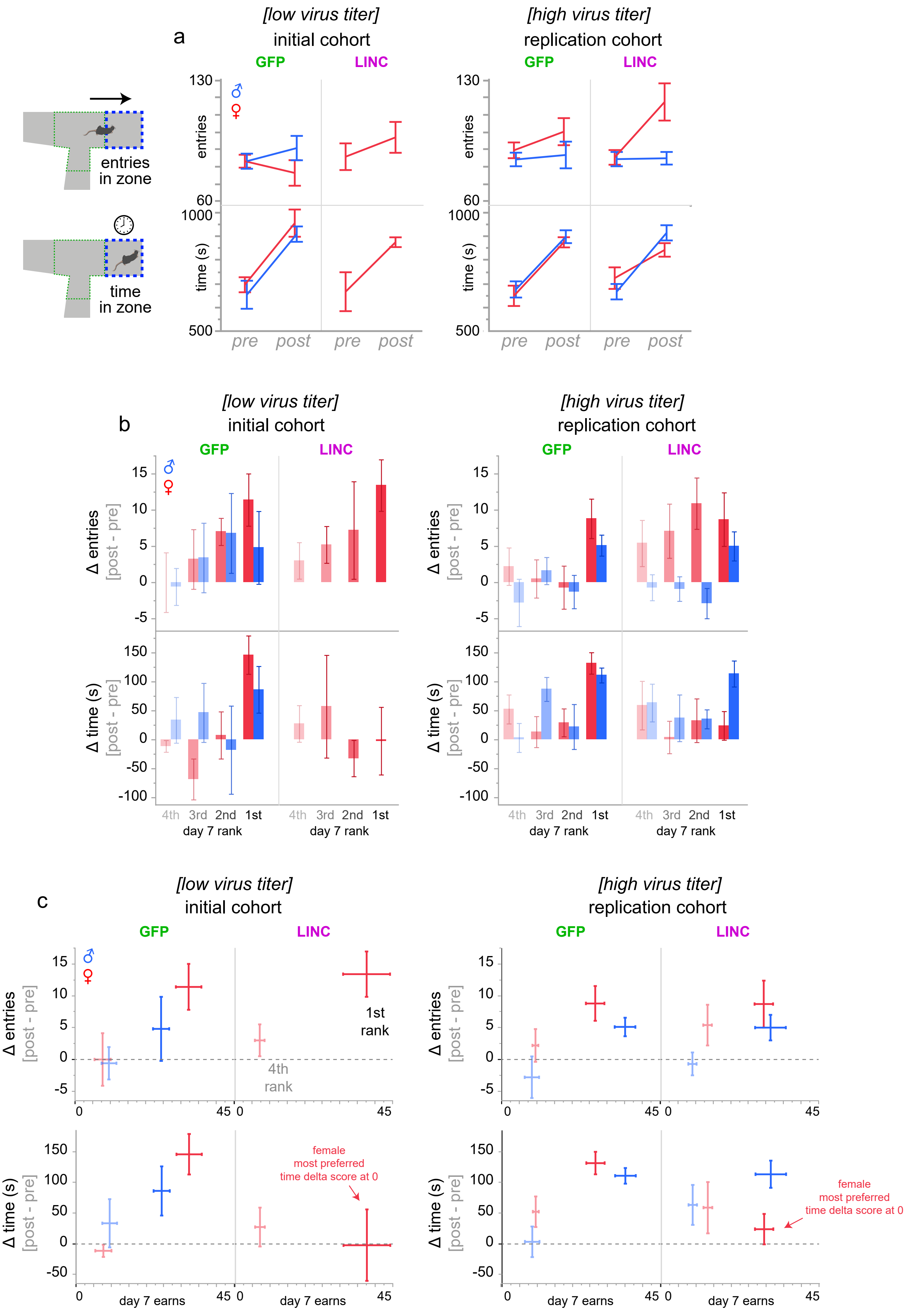

**Supplementary Fig. 4 | Replication cohort of two different virus titers on sex-specific Restaurant Row conditioned place preference (CPP) findings.** Animals were placed in the arena free to roam for 20 min on day 0 and day 8 of the experimental timeline but with no active task to obtain pre-task baseline and post-task experience-related exploratory behavior (see main Fig. 2). **a** Total number of entries and total time spent in the wait zone, aggregated in all restaurants. Depicting findings from independent cohorts: surgeries and behavioral experiments were carried out across separate calendar months and using different virus titers. **b** Delta scores in change of entries and time comparing post (day 8) minus pre (day 0) time points split by restaurants ranked according to rewards earned on the active task on day 7. **c** Data from (b, only 1$^{st}$ and 4$^{th}$ ranking restaurants) replotted with number of rewards earned on day 7 on the x-axis. Horizontal dashed gray line indicates delta score of 0. LINC00473 expression in mPFC abolished CPP behavior in most preferred restaurants only in females and only in the time but not entry domain, replicated in both cohorts. +Sign-test on delta score of time: low virus cohort: male GFP: ($t$=2.142, $p$<0.05); female GFP: ($t$=4.424, $p$<0.01); female LNC: ($t$=0.04, $p$=0.516); high virus cohort: male GFP: ($t$=8.681, $p$<0.0001); male LINC: ($t$=5.087, $p$<0.001); female GFP: ($t$=7.157, $p$<0.0001); female LNC: ($t$=0.962, $p$=0.181). +Sign-test on delta score of entries: low virus cohort: male GFP: ($t$=0.955, $p$=0.197); female GFP: ($t$=3.155, $p$<0.05); female LNC: ($t$=3.766, $p$<0.01); high virus cohort: male GFP: ($t$=3.520, $p$<0.01); male LINC: ($t$=2.493, $p$<0.05); female GFP: ($t$=3.236, $p$<0.01); female LNC: ($t$=2.355, $p$<0.05). Error bars represent ±1 SEM.

Supplementary Figure 5

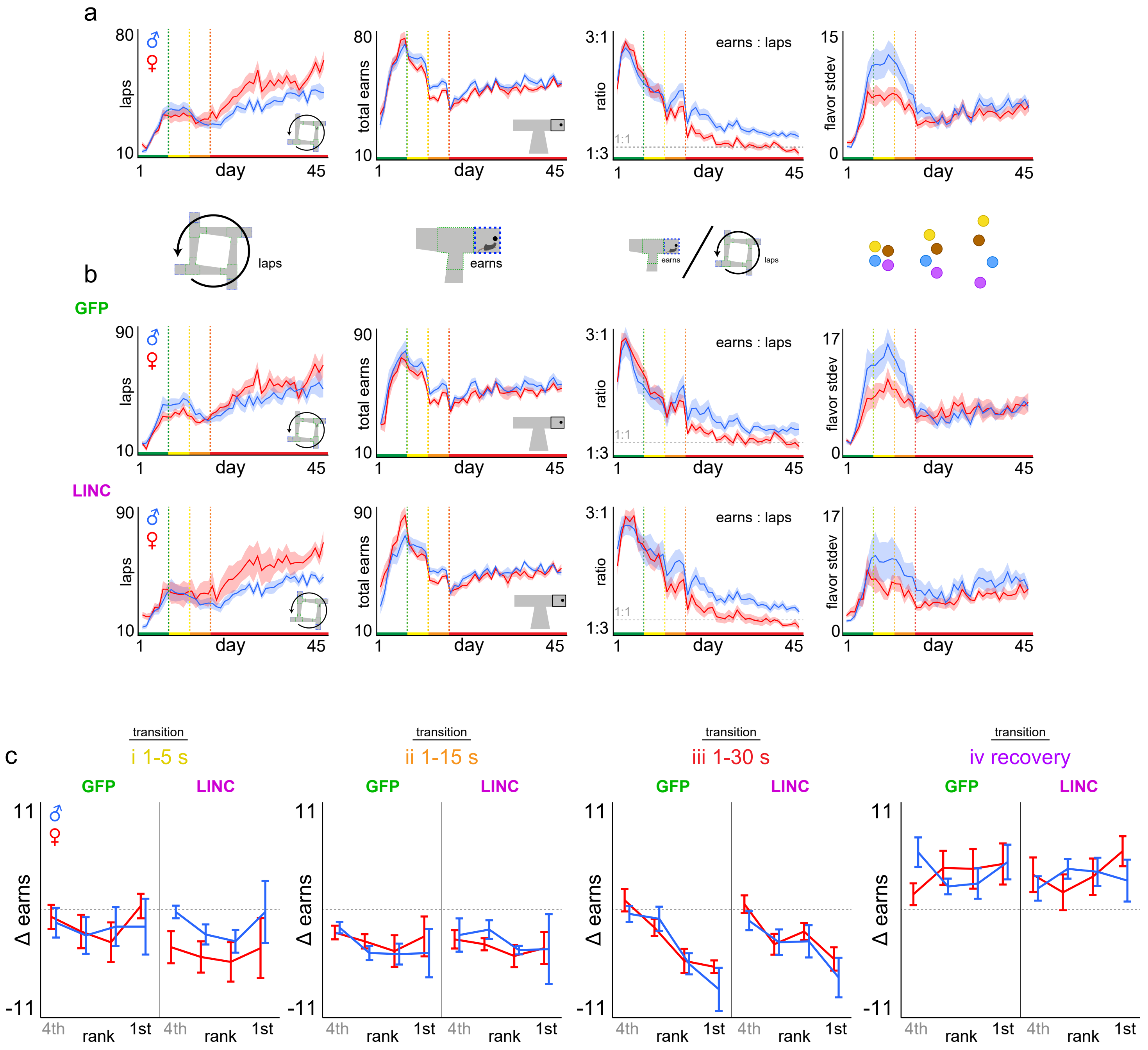

**Supplementary Fig. 5 | Basic Restaurant Row laps and earnings metrics across testing. a** Overall sex difference comparisons between males and females (collapsing across GFP and LINC00473 virus treatment within each sex). Average number of laps run in the correct direction (sex*day: $F$=78.040, $p$<0.0001), total rewards earned (sex*day: $F$=1.331, $p$=0.249), ratio of earns / laps (sex*day: $F$=28.155, $p$<0.0001), and standard deviation of earns among flavors (sex*day: $F$=13.014, $p$<0.001). **b** Data from (a) split by GFP and LINC groups: laps (sex*virus*day: $F$=0.008, $p$=0.927), total earns (sex*virus*day: $F$=7.470, $p$<0.01), ratio of earns / laps (sex*virus*day: $F$=1.130, $p$=0.288), standard deviation of earns by flavor (sex*virus*day: $F$=15.486, $p$<0.0001). Statistical tests from (a) and (b) were performed on the entire experimental timeline (days 1-45). **c** Data from main Fig. 3d split by sex and virus groups depicting change in rewards earned in each restaurant comparing two days at each color-coded transition point. Delta scores calculated by subtracting the following: i, day 8-7 (yellow, rank: $F$=1.361, $p$=0.259; sex*virus: $F$=0.017, $p$=0.897); ii, day 13-12 (orange, rank: $F$=1.123, $p$=0.343; sex*virus: $F$=0.086, $p$=0.769); iii, day 18-17 (red, rank: $F$=18.116, $p$<0.0001; sex*virus: $F$=0.076, $p$=0.783); iv, day 45-18 (purple, rank: $F$=0.603, $p$=0.614; sex*virus: $F$=0.375, $p$=0.542). Shaded area and error bars represent ±1 SEM.

Supplementary Figure 6

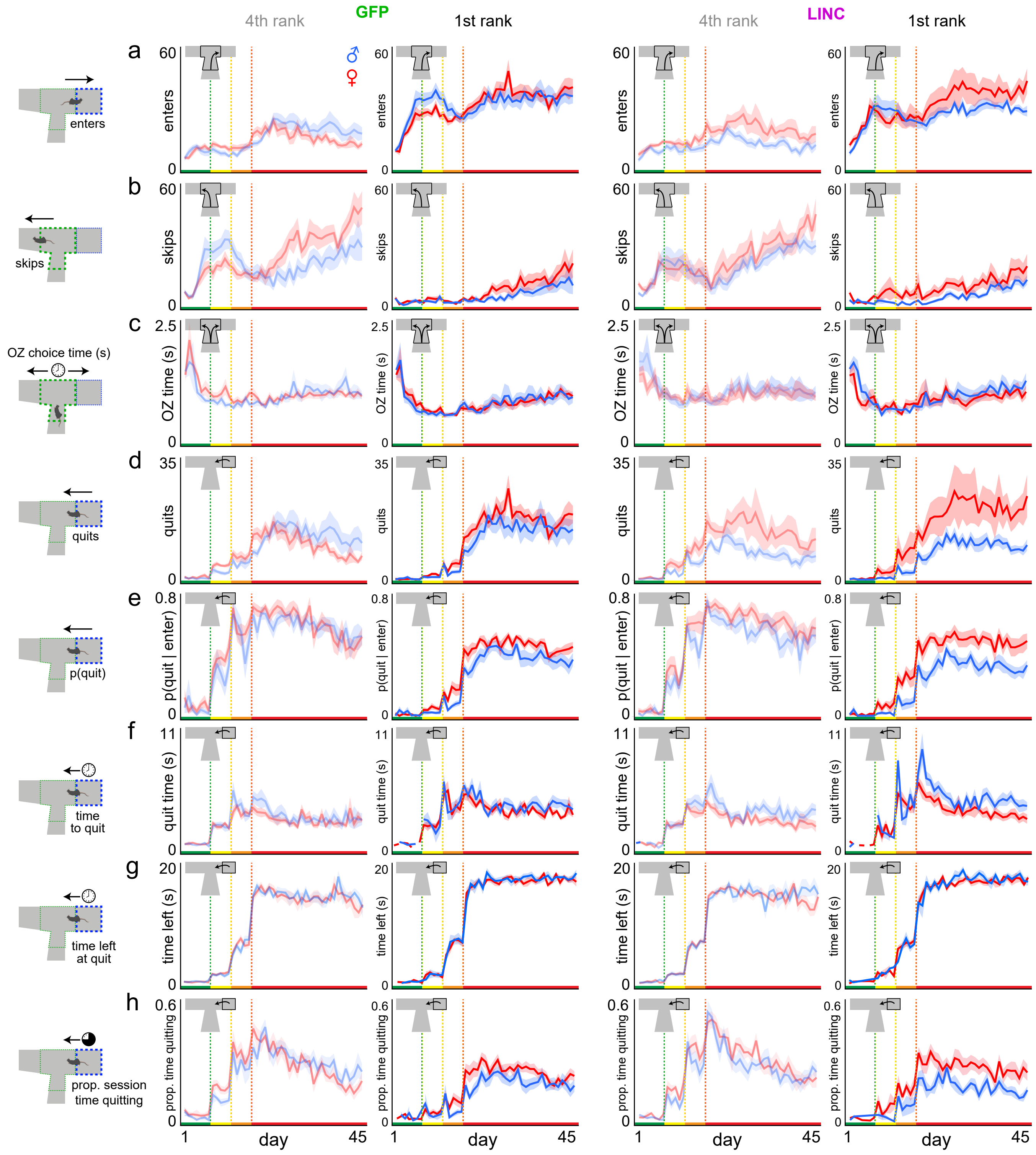

**Supplementary Fig. 6 | Restaurant Row choice metrics split by flavor ranking across testing. a** Average number of enter decisions made in the offer zone split by sex and virus treatment groups as well as the 4$^{th}$ and 1$^{st}$ ranking restaurant (sex*rank*day: $F$=16.043, $p$<0.0001; sex*virus*rank*day: $F$=3.967, $p$<0.01). **b** Number of skip decisions made in the offer zone (sex*rank*day: $F$=9.894, $p$<0.0001; sex*virus*rank*day: $F$=2.617, $p$<0.05). **c** Offer zone reaction time before making an enter or skip decision calculated from offer zone entry (tone onset) to either a right turn into the wait zone (enter) or a left turn into the hallway (skip) (sex*rank*day: $F$=0.325, $p$=0.808; sex*virus*rank*day: $F$=2.186, $p$=0.087). **d** Number of quit decisions made in the wait zone during a countdown (sex*rank*day: $F$=11.725, $p$<0.0001; sex*virus*rank*day: $F$=1.216, $p$=0.302). **e** The probability of quitting in the wait zone given the animal entered (sex*rank*day: $F$=4.145, $p$<0.01; sex*virus*rank*day: $F$=0.463, $p$=0.708). This normalizes the likelihood of quitting to enter frequency. **f** Wait zone latency to quit reaction time from countdown onset (wait zone entry) until mice leave the wait zone prematurely (sex*rank*day: $F$=1.427, $p$=0.233; sex*virus*rank*day: $F$=3.160, $p$<0.05). **g** The amount of time left remaining in the countdown required to finish earning a reward at the moment of quitting in the wait zone. **h** Proportion of session time within each restaurant spent engaged in quitting behavior in the wait zone. Statistical tests here were performed on the entire experimental timeline (days 1-45). Shaded area represents ±1 SEM.

Supplementary Figure 7

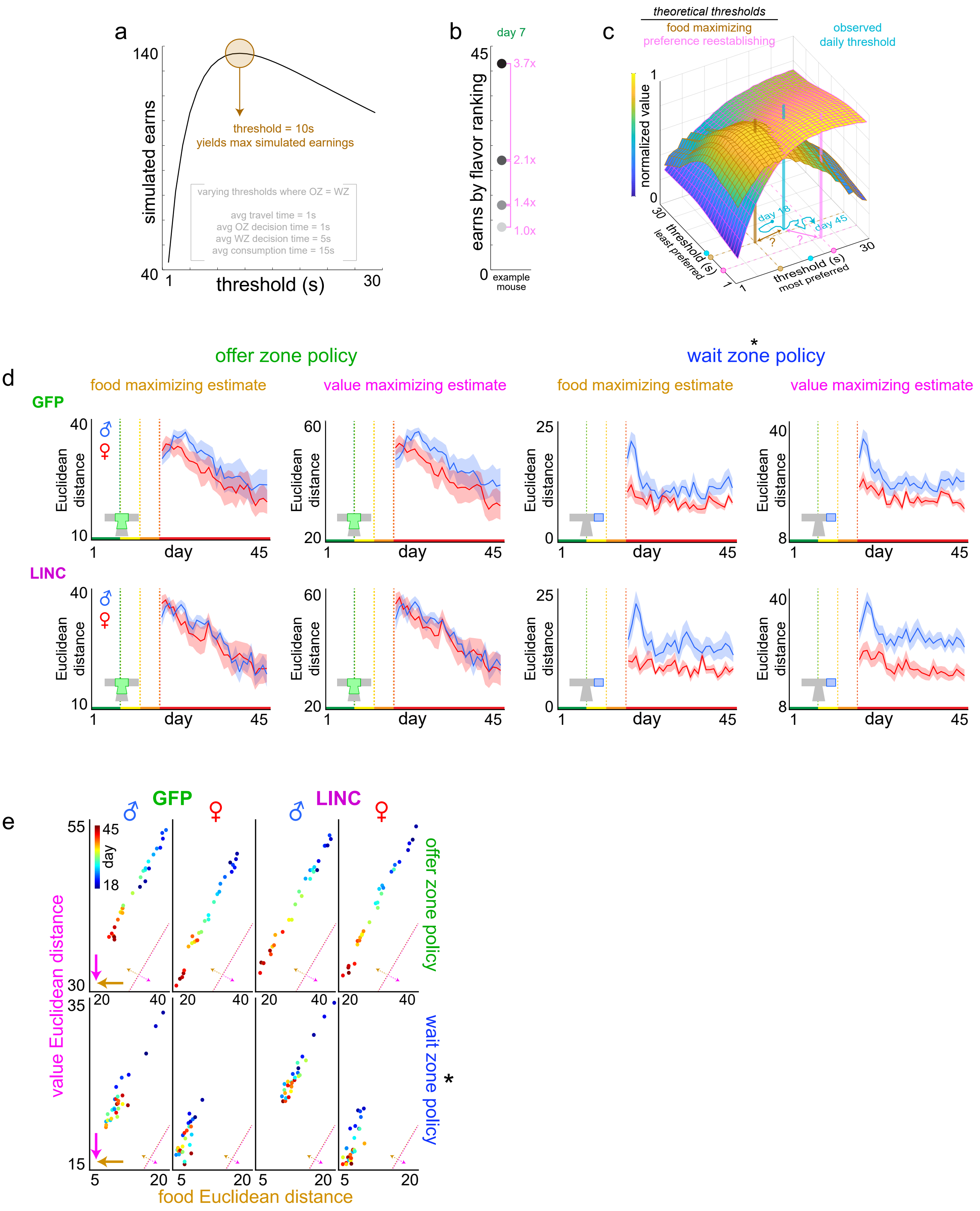

**Supplementary Fig. 7 | Longitudinal economic analysis of decision policy optimization for food security vs. subjective value.** Analysis underlying data from main Fig. 4g-i derived from Durand-de Cuttoli et al 2023 *Biological Psychiatry* [43]. See **Methods** for more details on this analysis. **a** Computer Restaurant Row simulation of total number of pellets earned. The ideal threshold required to obtain the theoretical maximum number of pellets when ignoring flavor preferences was empirically determined to be 10 s. **b** Relative ratio of earns among restaurants for an example mouse on day 7 between the flavor rankings (i.e., this mouse had a 3.7 : 2.1 : 1.4 : 1 ratio of earns across flavors capturing a summary of idealized relative subjective value for this mouse when all offers were 1 s only). **c** Two intersecting planes of decision policies that yield varying amounts of theoretical value either for maximal food intake (as determined by computer simulations, brown plane) or subjective value (as determined by multiplying simulation output of flavor earnings by day 7's preference ratios on a mouse-by-mouse basis, pink plane) when in a reward-scarce environment (1 to 30 s offers). Here, only two decision policy dimensions (least preferred and most preferred restaurants) of the four dimensions (all four ranked restaurants) are displayed. Both planes are normalized to each's min and max values for the purpose of plotting both in the same space while preserving the threshold coordinate locations that achieve either theoretical maximum (brown [location always fixed at threshold coordinates of 10 s] or pink beacons [location of discovered threshold coordinates that vary from mouse-to-mouse]). Actual observed daily decision policies represented by the cyan beacon wander throughout this space from day 18 to 45 in a reward-scarce environment. Trajectories projected to the floor of this display trace out individual mouse decision policy paths. Example coordinates of brown, pink, and cyan beacons illustrated as dots on the x and y axes. Euclidean distance from cyan coordinates to either brown or pink coordinates were calculated (question mark symbols). **d** Euclidean distances from observed decision policies in the offer zone (left) or wait zone (right) to either food (brown) or preference (magenta) theoretical maximum across testing in a reward-scarce environment split by GFP (top) and LINC (bottom) groups and sex. Offer zone: food distance: day: $F$=49.362, $p$<0.0001; sex: $F$=1.115, $p$=0.298; virus: $F$=0.008, $p$=0.929; value distance: day: $F$=42.042, $p$<0.0001, sex: $F$=1.546, $p$=0.221; virus: $F$=0.001, $p$=0.989; Wait zone: food distance: day: $F$=16.774, $p$<0.001, sex: $F$=13.796, $p$<0.001; virus: $F$=0.598, $p$=0.445; value distance: day: $F$=38.046, $p$<0.0001, sex: $F$=18.034, $p$<0.0001; virus: $F$=0.261, $p$=0.613. **e** Data from (d) redisplayed showing the Euclidean distance between observed decision policies each day from days 18-45 (cool to warm colors, each dot indicates group average per day) to theoretical policies that could achieve either maximal food (x-axis) or maximal subjective value (y-axis) for offer zone and wait zone thresholds. Inset diagonal lines indicate reference point for the 1:1 unity line. Bold brown and magenta arrows pointing toward the origin represent direction of reduced Euclidean distance vectors that optimize either food or value, respectively. Dashed brown and magenta arrows pointing off diagonal from the unity line represent if the decision policy is biased to prefer food vs. value. Prominent sex difference, without apparent effect of LINC00473 expression in mPFC, in present in wait zone but not offer zone economic decision policies. Shaded area represents ±1 SEM.

Supplementary Figure 8

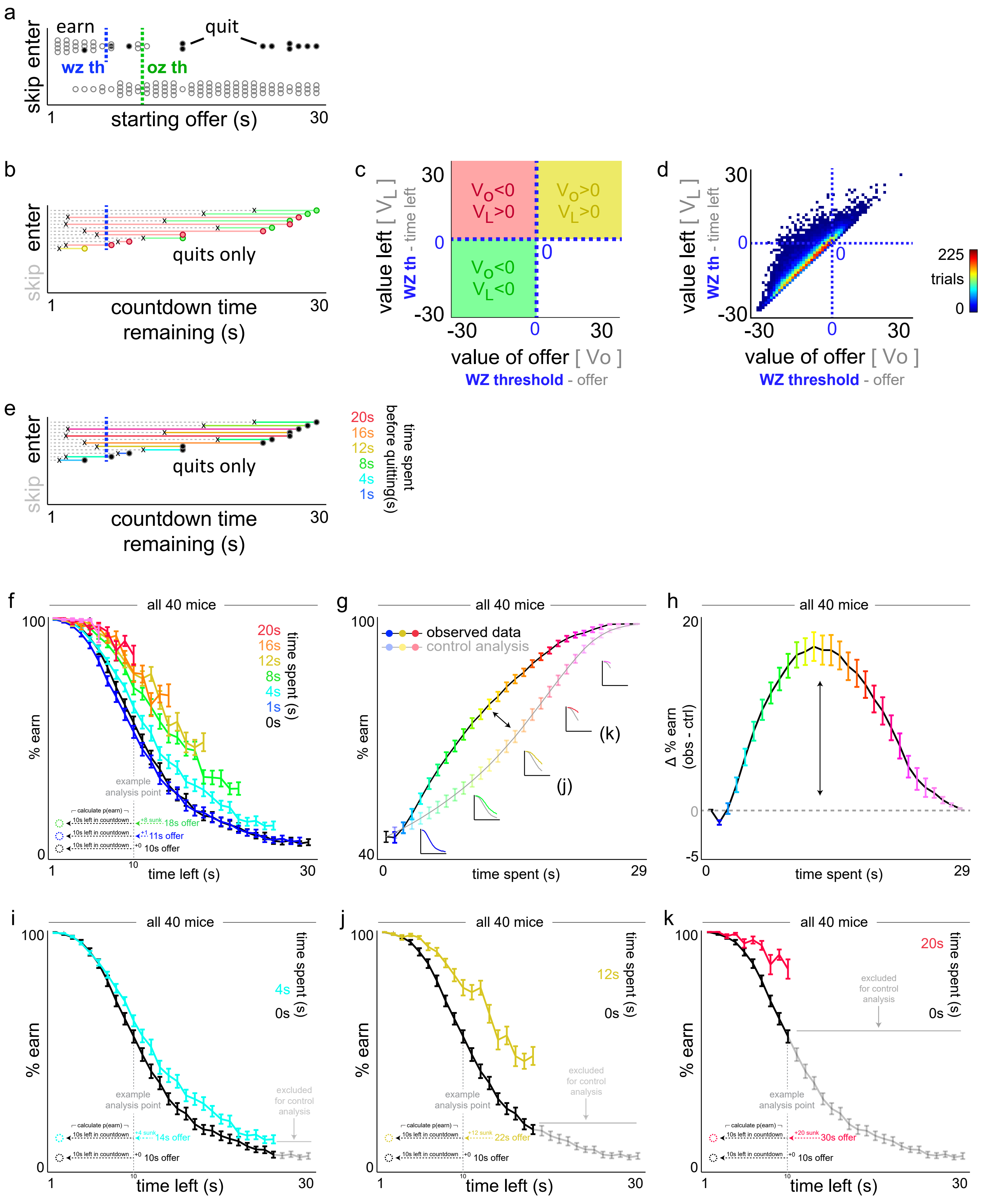

**Supplementary Fig. 8 | Visual explanation of characterizing quitting behavior. a** Example choice data from a single mouse, single session, in a single restaurant. Individual dots represent individual trials. Offer zone outcomes plotted on the Y-axis as a function of the cued offer at the start of each trial along the X-axis. Of the enter trials, wait zone outcomes represent earns as open circles and quits as closed circles. Vertical dashed green and blue lines indicate the offer zone and wait zone thresholds, respectively. Note: the only temporal data shown here is the starting delay at the onset of the trial, and how each trial terminates as either a skip, enter then earn, or enter then quit. Only panels b-e include temporal data about how long it took animals to quit, for instance, and conversely, how much time was remaining in the countdown at the moment of quitting. **b** Data from (a) showing only the quit trials. Horizontal solid-colored lines extending from each dot leftward represent how much time was spent waiting before the mouse decided to quit. The point at which the mouse quit is represented by the black "x" symbol. The remaining dashed gray lines extending from the "x" leftward thus represent the amount of time remaining in the countdown at the moment of quitting. The relationship between one's own wait zone threshold (vertical dashed blue line) and whether or not (i) the starting offer [circle] is to the right or the left of the wait zone threshold and (ii) the amount of time remaining at the moment of quitting ["x"] is to the right or the left of the wait zone threshold determines which quadrant each quit event belongs to in panels (c-d). Thus, each quit event is color coded based on its placement in the quadrants in (c-d). **c-d** Offer value plotted against value left can divide up this quit space into a two-by-two matrix based on the sign of each value term [i.e., if the starting offer was above or below one's own wait zone threshold [represented by the dashed blue 0 lines] and if the time left at the moment of quitting was above or below one's own wait zone threshold]. Data can only possibly exist in the three colored green, yellow, and red quadrants. Schematic in (c) and actual example data from main Fig. 5a re-displayed here. **e** Lastly, the time spent waiting before quitting is color-coded in yet another way, here based on how much time was already invested prior to the quit decision being made. Thus, this can separate rapid quit events from longer latencies to quit. These measures form one dimension of the data used in the sunk cost analysis, which also factors in varying amounts of time left remaining (horizontal gray dashed lines that represent the residual time at the moment of quitting). **f-k** Explanation of sunk cost and control analysis. (f-h) Redisplay of data from main Fig. 5f-h. Sunk cost analysis of staying behavior in the wait zone, demonstrated using all mice. (f) The likelihood of staying in the wait zone and earning a reward (e.g., not quit) is plotted as a function of time left in the countdown along the x-axis and time already spent waiting orthogonally in color. Note the black 0 s time spent curve represents animals having just entered the wait zone from the offer zone. Inset vertical dashed gray line illustrates an example analysis point comparing three sunk cost conditions originating from different starting offers but matched at 10 s left. Data from (f) dimensioned reduced in (g) collapsing across time left, instead highlighting the grand mean of each time spent sunk cost condition (color and x-axis). Insets depict data from curves in (f) are collapsed into the observed (sunk condition) and control (0 s condition) lines. Difference between curves in (g) are plotted in (h) in order to summarize the envelope of the overall effect of time already spent on escalating the commitment of staying in the wait zone. Horizontal dashed line represents a delta score of 0. (i-k) Redisplay of data from (f) but for three sunk cost conditions indicated in the insets in (g, 4 s cyan in i, 12 s gold in j, 20 s red in k) with the 0 sunk cost condition (black curve) repeated in each panel. Because each colored sunk cost curve derives from trials where the starting offer cost is higher than the matched time left value of the 0 s cost condition, some data points do not exist and are missing from the sunk cost curve on the rightward end. Thus, when reducing dimensions, to control for the inflated summary calculations of % earn due to missing data alone, the control analysis iteratively excludes matched data from the black 0 s sunk curve (highlighted in gray) before collapsing data to yield the resultant control curve in (g). Error bars represent ±1 SEM.

Supplementary Figure 9

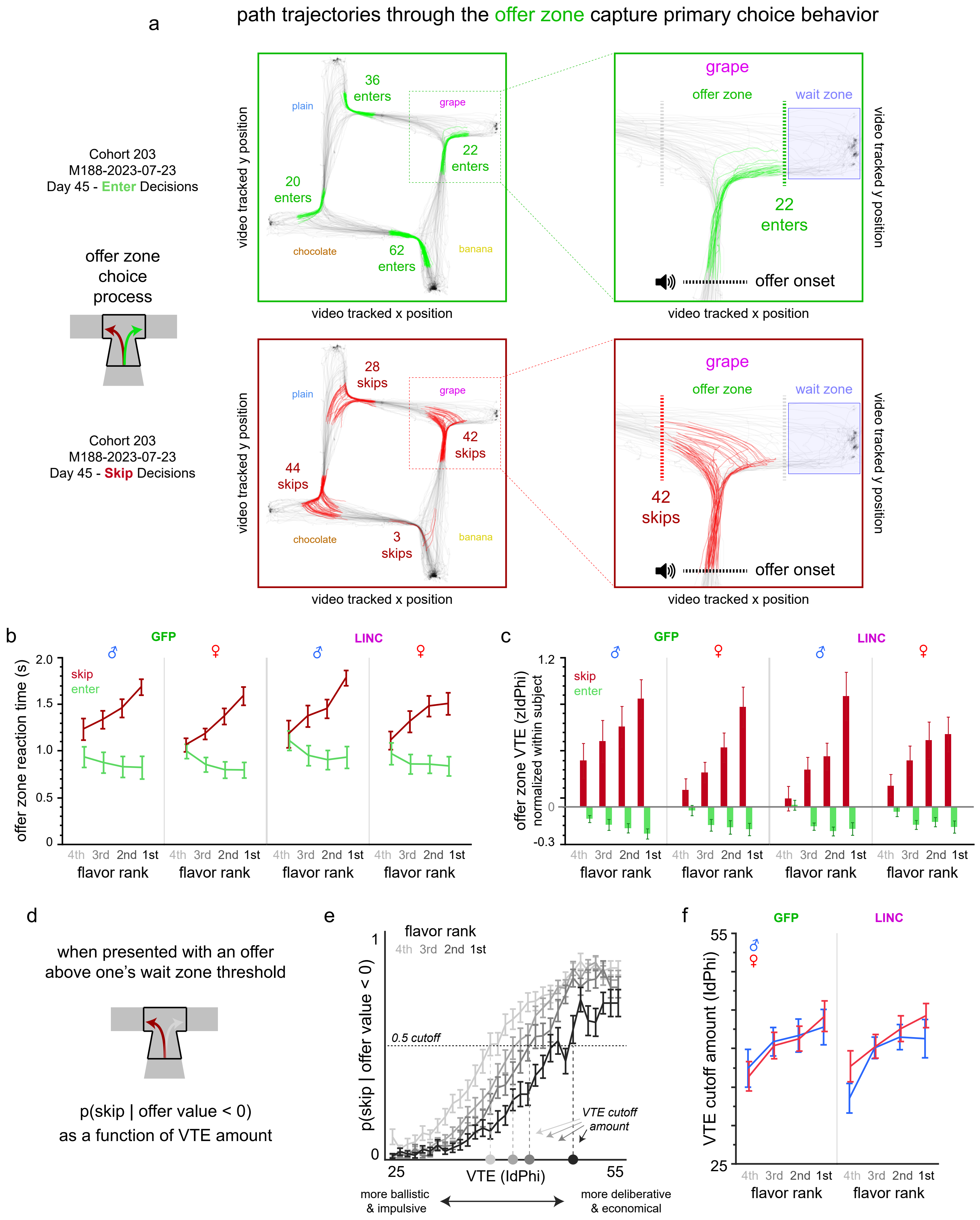

**Supplementary Fig. 9 | Video-tracking analysis of choice behavior in the offer zone. a** Example video tracking data during a full 30-min session obtained from one example mouse (M188) taken from day 45 of testing. Track plots display X-Y body centroid position data of the mouse at 30 fps as the animal navigates the maze arena performing the task. All path trajectories through the offer zone on each trial beginning at trial onset (triggered by body centroid position crossing the stem of the entrance into the restaurant, which also triggers the auditory cue onset) until animals turn right and enter the wait zone (triggered by body centroid position crossing the wait zone boundary, which then triggers the countdown onset) are color-coded in green. All other position data are color-coded in gray. The number of enter decisions made in each restaurant are displayed. Zoomed-in example shows a closer depiction of the grape restaurant. The same data is redisplayed again but for skip decisions color-coded in red. Note the significantly greater heterogeneity in path trajectories through the offer zone, including a greater mix of heading-direction reorientation events before ultimately registering a skip outcome (triggered by body centroid position crossing the left boundary of the offer zone as animals enter the hallway connecting to the next restaurant) compared to enter decisions that are more uniformly ballistic and without path reorientations. **b** Offer zone reaction time measures plotted as a function of restaurant flavor preferences ranked from least to most preferred and split by whether or not mice made a skip or enter decision. Latencies to skip or enter were measured from trial and cued offer onset (traversing the stem entrance of each restaurant) until body centroid position crosses the rightward wait zone entry point or leftward hallway entry point. Significant interaction between choice outcome (enter vs. skip) and flavor ranking ($F$=14.201, $p$<0.0001) but no significant effects of sex ($F$=1.170, $p$=0.280) or virus ($F$=0.253, $p$=0.615). Reaction time alone does not necessarily capture individual differences in how ballistic vs. tortuous path trajectories can be through the choice point. Vicarious trial and error (VTE) behavior has been previously shown to capture heading-direction reorientation events and correlates with neural representations of deliberative decision-making processes (high VTE events) vs. the lack thereof (low VTE events). The physical "hemming and hawing" characteristic of VTE is best measured by calculating changes in velocity vectors of discrete body $X$ and $Y$ positions over time as $dx$ and $dy$. From this, we can calculate the momentary change in angle, *Phi*, as *dPhi*. When this metric is integrated over the duration of the pass through the offer zone, VTE is measured as the absolute integrated angular velocity, or *IdPhi*, until either a skip or enter decision was made. This measure importantly is normalized within subject by z-scoring *IdPhi* values across all choices within a given session on a mouse-by-mouse basis, and then subsequently splitting the data by choice outcome and restaurant in order to appreciate within subject differences of how different types of choices and in different restaurants elicit varying amounts of either high or low VTE events. Significant interaction between choice outcome (enter vs. skip) and flavor ranking ($F$=19.819, $p$<0.0001) but no significant effects of sex ($F$=0.181, $p$=0.671) or virus ($F$=0.374, $p$=0.541). **d-f** Analysis of VTE behavior depicting the probability of skipping in the offer zone a negatively valued offer (offer above one's wait zone threshold) as a function of binned VTE amounts. (e) Skipping probability given offer value <0 as a function of VTE amount split by restaurants ranked by flavor preferences for all 40 mice. Horizontal dashed line indicates a probability cutoff of 0.5 used to determine the X-intercept (vertical dashed lines shaded per restaurant), or amount of VTE displayed in order to reliably skip an economically disadvantageous offer. (f) VTE cutoff amount split by flavor rank, sex, and LINC00473 treatment. Significant effect of flavor ranking ($F$=9.372, $p$<0.0001) but no significant effects of sex ($F$=0.464, $p$=0.497) or virus ($F$=1.330, $p$=0.251). Data in (b-f) collapsed across the 1-30 s epoch (days 18-45). These data indicate that LINC00473 treatment nor sex has an impact on offer zone choice processes, including leaving intact deliberative behaviors that interact with the type of decision being made and counteract pre-potent impulsive responses to enter depending on the degree of flavor preference.

Supplementary Figure 10

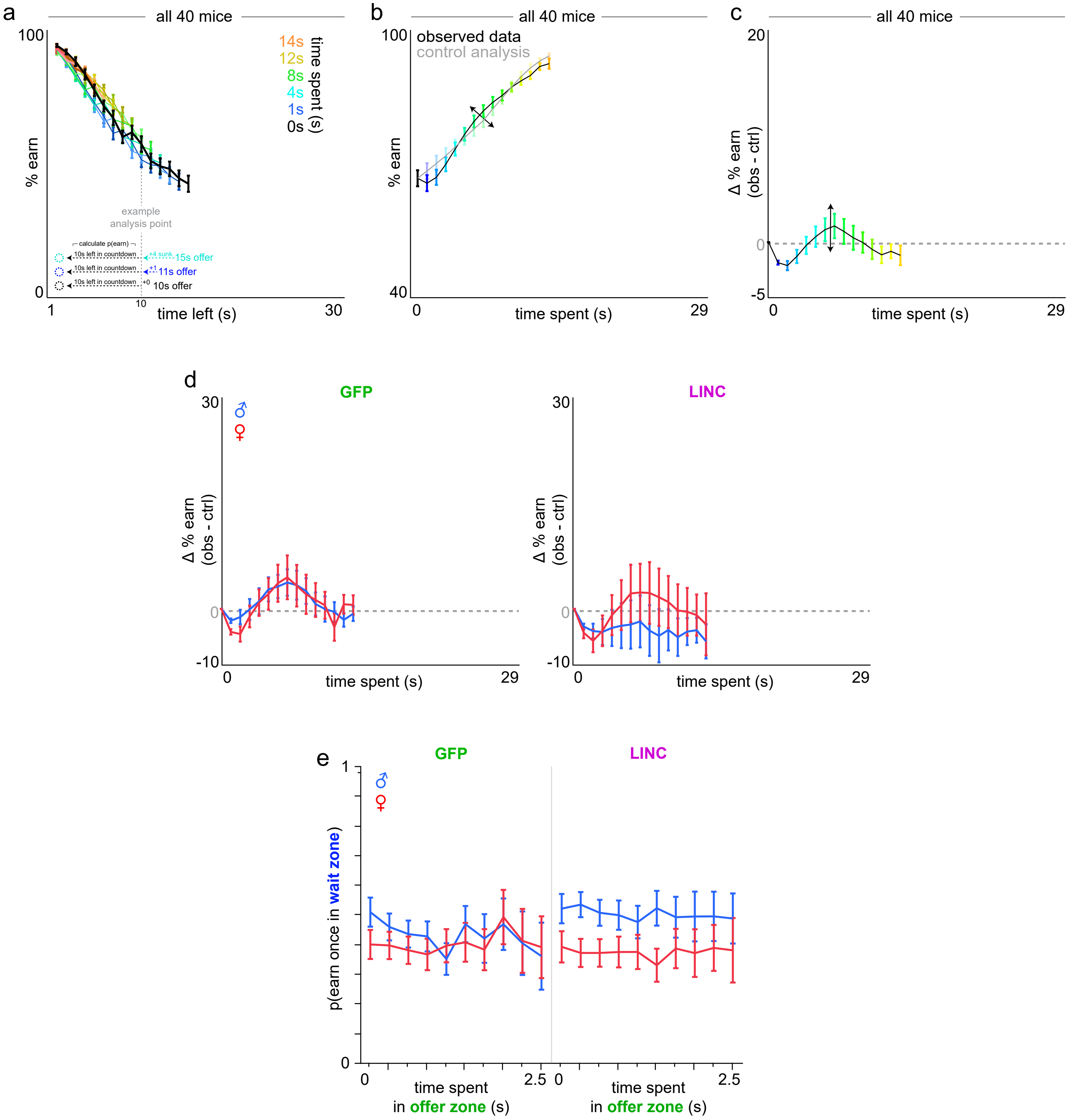

**Supplementary Fig. 10 | Sensitivity to sunk costs analysis during the 1-15 s testing epoch.** Mice do not display sensitivity to sunk costs during days 13-17 in a relatively reward rich environment compared to days 18+ (when offers range 1-30 s, as in main Fig. 5). **a-c** Sunk cost analysis of staying behavior in the wait zone, demonstrated using all 40 mice. (f) The likelihood of staying in the wait zone and earning a reward (e.g., not quit) is plotted as a function of time left in the countdown along the x-axis and time already spent waiting orthogonally in color. Note the black 0 s time spent curve represents animals having just entered the wait zone from the offer zone. Inset vertical dashed gray line illustrates an example analysis point comparing three sunk cost conditions originating from different starting offers but matched at 10 s left. Data from (a) dimensioned reduced in (b) collapsing across time left, highlighting the grand mean of each time spent sunk cost condition (color and x-axis). Difference between curves in (b) are plotted in (c) in order to summarize the envelope of the overall effect of time already spent on escalating the commitment of staying in the wait zone. Horizontal dashed line represents delta score of 0. **d** Data in (c) split by sex and virus treatment groups depicting no effect of time spent during the countdown in the wait zone on changing the probability of quitting vs. staying to earn a reward (time spent: $F$=0.522, $p$=0.470; sex*virus*time spent: $F$=0.809, $p$=0.369). **e** To compare to time spent waiting in the wait zone, data here depict a different type of time spent on the task: time spent in the offer zone prior to accepting an offer and entering the wait zone does not influence the probability of quitting vs. staying to earn a reward once in the wait zone (time spent in offer zone: $F$=1.574, $p$=0.210; sex*virus*time spent: $F$=0.916, $p$=0.339). These data are from the 1-30 s reward-scarce epoch (days 18-45) to demonstrate the unique value of time spent waiting in the wait zone uniquely conferring sunk cost sensitivity. Error bars represent ±1 SEM.

### Supplementary Figure 11

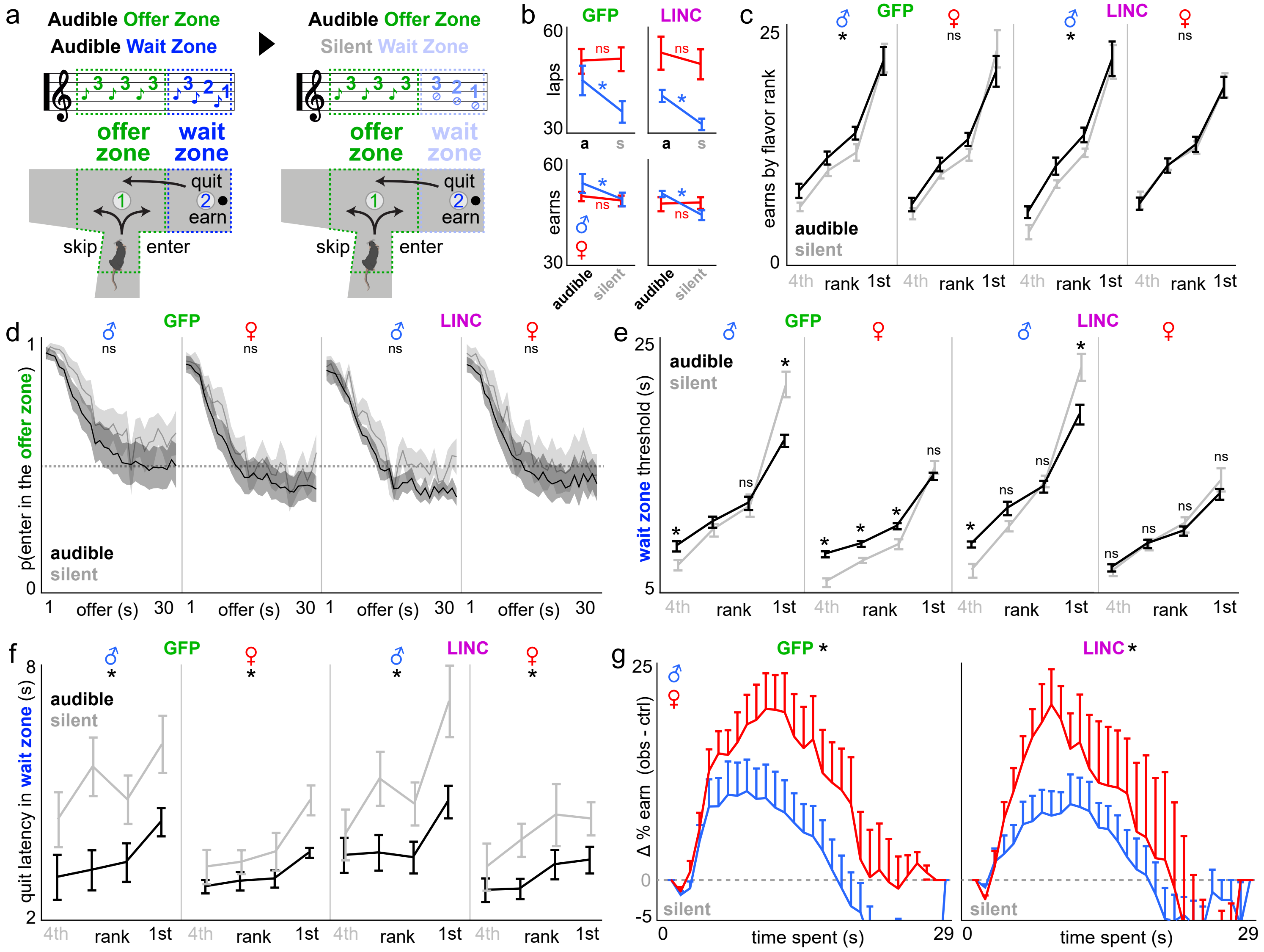

**Supplementary Fig. 11 | Sex- and LINC00473-dependent wait zone behaviors are differentially modulated by access to specific task information. a** Informational manipulation schematic: (left) standard Restaurant Row task followed by (right) two days of tones silenced only in the wait zone, but not offer zone (remains audible as before). **b** Total number of laps run in the correct direction (top) and total pellets earned aggregated among restaurants (bottom) across the audible and silent wait zone task conditions (x-axis). **c** Rewards earned in each restaurant ranked from least ($4^{th}$) to most preferred ($1^{st}$) across the audible (black) and silent wait zone (gray) task conditions. **d** Overall offer zone choice probabilities to make enter decisions as a function of audible cued offer costs collapsed across all flavors to illustrate no effect of task condition on auditory discriminatory behaviors. Horizontal dashed line indicates change at 0.5. **e** Wait zone thresholds in each restaurant across task conditions. **f** Latency to quit reaction times once in the wait zone in each restaurant across task conditions. **g** Sensitivity to sunk costs in the wait zone depicted here only in the silent condition as a function of time already spent (x-axis). Y-axis reflects delta earn probabilities as depicted in main Fig. 5h. Prominent sex-dependent effects of withholding wait zone information on overall task performance (observed in males), wait zone economic decision policies (bidirectional in males, unidirectional in GFP-females, and no effect in LINC00473-females), and sunk cost sensitivity (sex difference revealed for the first time GFP groups unlike in main Fig. 5). Horizontal dashed line indicates delta score of 0. ***** in (b-f) represent significant effects of task condition within group. * in (g) represent significant sex differences. Shading / error bars represent ±1 SEM. See **Supplementary Text**.

Supplementary Figure 12

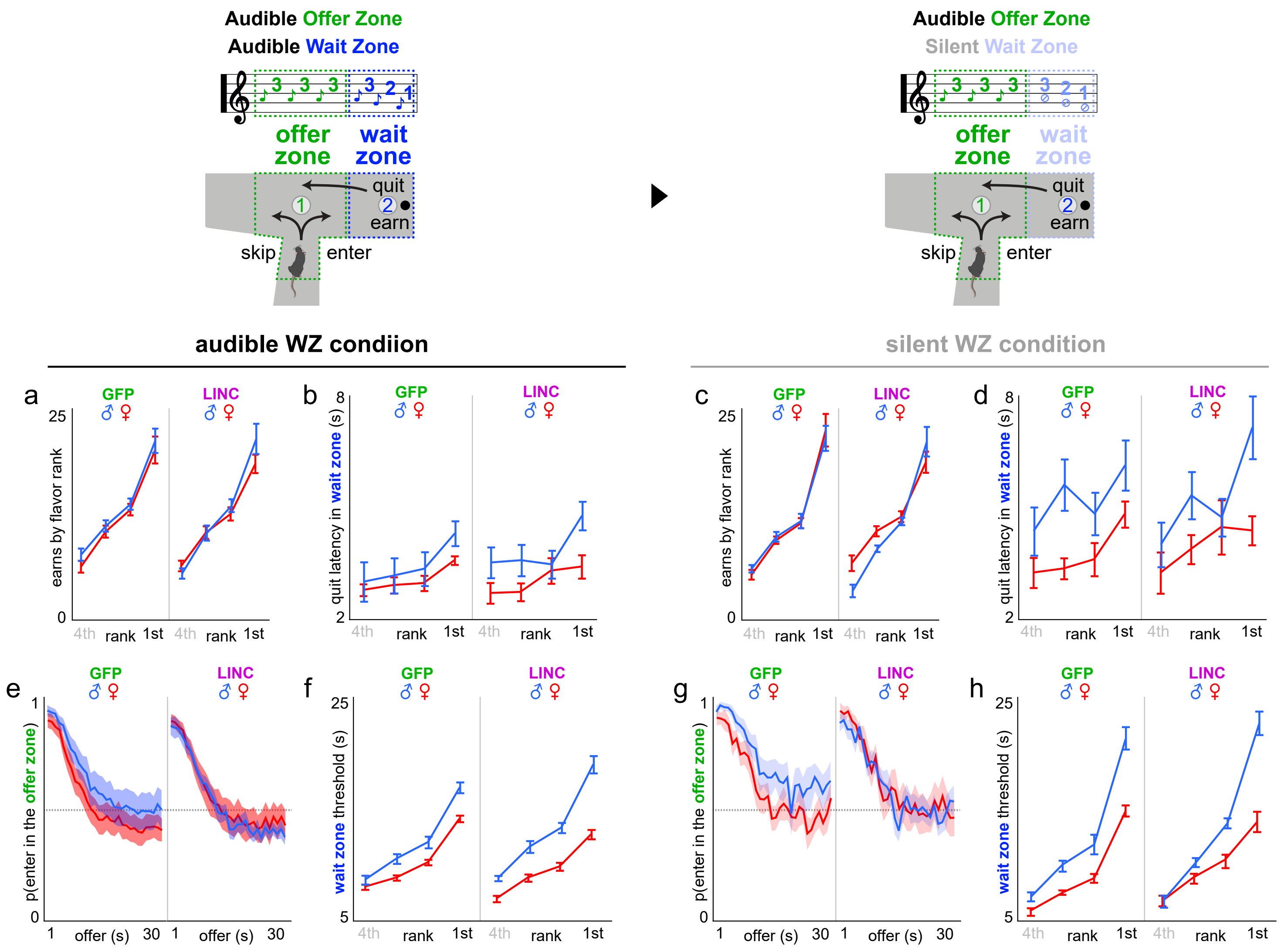

**Supplementary Fig. 12 | Redisplay of main metrics obtained from task information manipulation.** Figures redisplayed separating task manipulation conditions (baseline audible wait zone (WZ) vs. silent WZ conditions) and superimposing sex. Redisplay from Supplementary Fig. 11. **a,c** Average number of rewards earned split by flavors ranked from least to most preferred based on end-of-session totals. **b,d** Average latency to quit after accepting an offer once in the wait zone. **e,g** Average proportion of trials accepted by making an enter decision from the offer zone into the wait zone as a function of cued offer cost in the offer zone. **f,h** Average wait zone threshold calculated from the inflection point of Heaviside-step function fits to wait zone outcome (earn vs. quit) as a function of the starting delay of each trial. Error bars and shading represent ±1 SEM.

Supplementary Figure 13

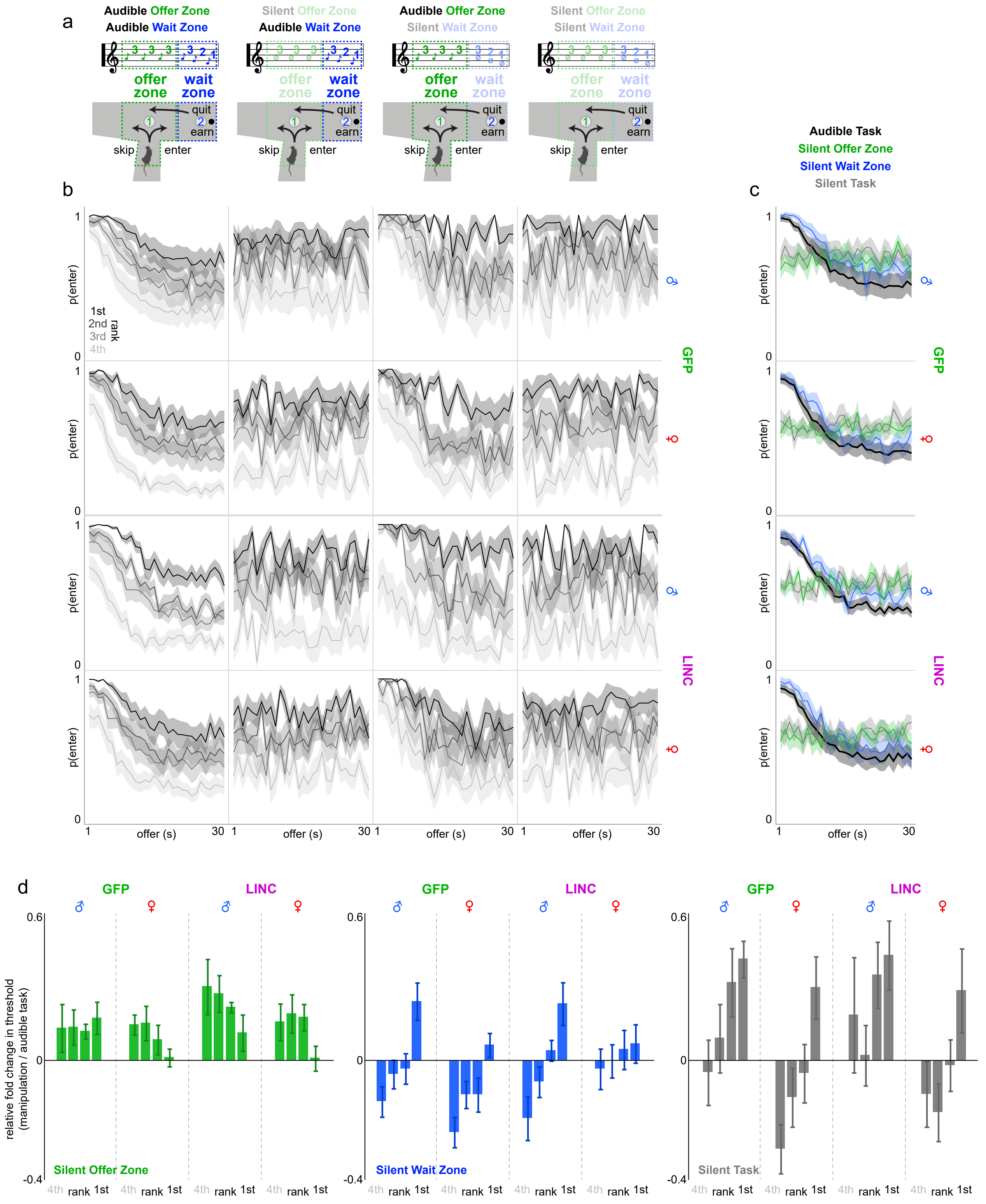

**Supplementary Fig. 13 | Full set of task information manipulations on Restaurant Row. a** After day 45, mice were challenged with special probe days on three different task informational manipulations each for 2 days. Manipulations took the form of silencing and withholding tone presentation either in the offer zone, wait zone, or both: (i) silent offer zone but audible wait zone [days 46-47], (ii) audible offer zone but silent wait zone [days 48-49], and (iii) silent offer zone and silent wait zone [days 50-51]. **b** Offer zone choice probability to enter as a function of offer cost split by restaurants ranked from least ($4^{th}$) to most ($1^{st}$) preferred separating sex and virus treatment groups (rows) across the 4 different task information conditions (columns). **c** Data in (b) collapsed across all four restaurants showing summary enter probabilities as a function of offer cost. Note that enter probabilities are largely unaffected when the offer zone tones are audible comparing baseline (black, regular task) to silent wait zone (blue) conditions, however, as intended, offer zone choice probabilities change into a flat horizontal line in both the silent offer zone (green) and silent task (gray) conditions, as animals do not have any tone information in the offer zone during these conditions (audible task: offer: $F$=2017.115, $p$<0.0001; sex*virus*offer: $F$=1.912, $p$=0.167; silent offer zone task: offer: $F$=1.783, $p$=0.182; sex*virus*offer: $F$=0.227, $p$=0.634; silent wait zone task: offer: $F$=1004.552, $p$<0.0001; sex*virus*offer: $F$=0.147, $p$=0.702; fully silent task: offer: $F$=0.723, $p$=0.395; sex*virus*offer: $F$=1.723, $p$=0.190). **d** Wait zone thresholds plotted as a relative fold change comparing each condition to the regular task audible baseline separated across the three information manipulations (silent offer zone, left, green; silent wait zone, middle, blue; silent task, right, gray) split by restaurants ranked from least ($4^{th}$) to most ($1^{st}$) preferred, sex, and virus treatment groups. Note the difference in patterns in changes in wait zone thresholds across restaurants between sex/virus groups and across task information manipulation conditions. Sign tests: silent offer zone: male GFP: least preferred: $t$=+1.365, $p$=0.103; most preferred: $t$=+2.639, $p$<0.05; male LNC: least preferred: $t$=+2.699, $p$<0.05; most preferred: $t$=+1.538, $p$=0.079; female GFP: least preferred: $t$=+3.638, $p$<0.01; most preferred: $t$=+0.360, $p$=0.363; female LNC: least preferred: $t$=+2.125, $p$=<0.05; most preferred: $t$=+0.188, $p$=0.427; silent wait zone: male GFP: least preferred: $t$=-2.684, $p$<0.05; most preferred: $t$=+3.191, $p$<0.01; male LNC: least preferred: $t$=-2.624, $p$<0.05; most preferred: $t$=+2.685, $p$<0.05; female GFP: least preferred: $t$=-4.733, $p$<0.001; most preferred: $t$=+1.325, $p$=0.109; female LNC: least preferred: $t$=-0.411, $p$=0.345; most preferred: $t$=+0.886, $p$=0.199; silent task: male GFP: least preferred: $t$=-0.358, $p$=0.364; most preferred: $t$=+5.554, $p$<0.001; male LNC: least preferred: $t$=+0.794, $p$=0.225; most preferred: $t$=+3.049, $p$<0.001; female GFP: least preferred: $t$=-3.584, $p$<0.01; most preferred: $t$=+2.345, $p$<0.05; female LNC: least preferred: $t$=-1.009, $p$=0.170**;** most preferred: $t$=+1.671, $p$=0.064. Shaded area and error bars represent ±1 SEM. See **Supplementary Text**.

Supplementary Figure 14

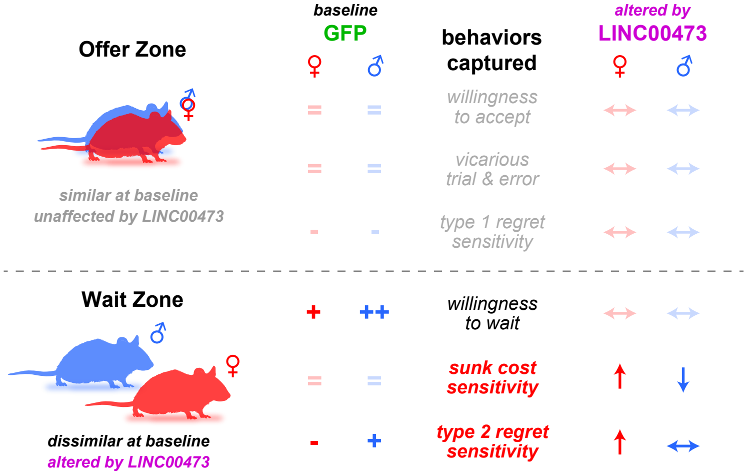

**Supplementary Fig. 14 | Expansion of summary of key findings.** Expanded table from main Fig. 6 describing in more detail the direction of baseline effects (GFP group) with equal signs (=) if no sex differences, plus signs (+ or ++) to represent a sex difference in magnitude, or minus signs (-) to indicate baseline insensitivity to a given behavioral metric. Metrics altered by mPFC LINC00473 expression are represented by arrow symbols showing either an increase (↑), decrease (↓), or no change (↔) relative to baseline. Bolded / opaque symbols highlight several key wait zone decision-making behaviors that are different between sexes at baseline or altered by LINC00473 expression, none of which appear in the offer zone.
